# Complex allometric relationships and ecological factors shape the development and evolution of eye size in the modular visual system of spiders

**DOI:** 10.1101/2023.12.28.573503

**Authors:** Kaylin Chong, Angelique Grahn, Craig D. Perl, Lauren Sumner-Rooney

## Abstract

Eye size affects many aspects of visual function, most notably including contrast sensitivity and achievable spatial resolution, but eyes are costly to grow and maintain. The allometry of eyes can provide insight to this trade-off in the context of visual ecology, but to date, this has mainly been explored in species that have a single pair of eyes of equal size. By contrast, animals possessing larger visual systems can exhibit variable eye sizes within individuals. Spiders have up to four pairs of eyes whose size often varies, but their ontogenetic, static, and evolutionary allometry has not yet been studied in a comparative context. We report variable evolutionary and developmental dynamics in eye size across 1098 individuals in 39 species and 8 families, indicating selective pressures and constraints driving the evolution of different eye pairs and lineages. We observed variation in the relationship between eye and body size not only between taxa and visual ecologies, but between eye pairs within species, indicating that growth dynamics are variable and can be divergently adapted in particular eye pairs. Supplementing our sampling with a phylogenetically comprehensive dataset recently published by Wolff et al. (2022), we confirmed these findings across more than 400 species, found that ecological factors such as visual hunting, web building, and diurnality can also impact eye diameter, and identified significant allometric shifts across spider phylogeny using an unbiased approach, many of which coincide with ecological changes such as visual pursuit hunting strategies. This study represents the first detailed comparative exploration of visual allometry in spiders, revealing striking differences in eye growth not only between families but also within species. Our findings shed light on the relationship between spider visual systems and their diverse ecologies, and how spiders exploit the modular nature of their visual systems to balance selective pressures and optical and energetic constraints.

## INTRODUCTION

Eyes are important sensory structures that allow organisms to spatially sample the light environment, and vision supports a wealth of behaviours ranging from the simple detection of shadows to complex communicative displays (Nilsson, 2009, 2013). Visual systems across the animal kingdom display astounding diversity, not only in the structure and function of individual eyes, but in their respective sizes, numbers, and placement (Cronin et al., 2014; Land, 2018; Land & Nilsson, 2012; Nilsson, 2021). Features of the eye can directly affect its ability to extract information from their environment, and these sometimes introduce conflict. Eyes must often balance contrast sensitivity against the ability to resolve spatial detail (acuity) and change over time (temporal resolution) (Dvorak & Snyder, 1978; Greiner, 2006; Klaus & Warrant, 2009; Land, 1984, 1997). The relative importance of these features vary depending on the visual environment and the ecology of the animal, and this may be reflected in eye morphology and development. For example, while wider photoreceptors can improve contrast sensitivity, this often comes at the cost of reduced spatial resolution (Robson, 1966; Owsley, 2003). One way to resolve this conflict is to make the eye larger: an eye with a wider aperture can collect more photons and support greater sensitivity, and a longer focal length can support greater spatial resolution (Caves et al., 2017, 2018; Land & Nilsson, 2012; Rutowski et al., 2009; Taylor et al., 2019). As a result, eye size can be a good indicator of both sensitivity and achievable resolution, and can provide insight to the relative importance of vision, particularly in dim light environments (Freelance et al., 2021; Land & Nilsson, 2012; Tierney et al., 2017). However, the potential benefits of larger eyes may not outweigh the required cost of investment, and the energetic budget allocated to the development of visual systems ultimately depends on this balance (Niven & Laughlin, 2008). Species that have relatively little use for vision, such as those living in caves and other dark environments, often exhibit reduced or even absent eyes that are more energetically economical (e.g. Arenz et al., 2018; Pérez-Moreno et al., 2018; Porter & Sumner-Rooney, 2018; Sumner-Rooney, 2018; Tierney et al., 2018).

In the context of visual ecology, allometry can provide insight to the trade-off in the growth and maintenance of eyes. Ontogenetic visual allometry describes the relationship between eye and body size within an individual’s lifetime. Generally, ontogenetic allometric relationships seem to be negative or hypoallometric, meaning the eyes grow more slowly than the body, especially in visually reliant species (Sakurai & Ikeda, 2022; Shrimpton et al., 2021; Stöckl et al., 2022). Ecological factors such as foraging mode and trophic level may correlate to these allometric relationships (Hall & Ross, 2007; Ross & Kirk, 2007). Phylogenetic visual allometry - the evolutionary relationship between eye and body size - has been explored in a range of species. Again, generalised relationships show eye size scaling hypoallometrically with body size or mass (e.g. in anurans, Huang et al., 2019; Thomas et al. 2020), but relative eye sizes also vary across phylogeny; for example, birds and primates tend to have larger eyes for a given body mass than rodents and reptiles (Howland et al., 2004), and within birds, the allometric relationship between head-space and body size differs between passerines and non-passerines (Burton, 2008). In addition, ecological factors such as habitat structure, predation pressure, and light availability, and other intrinsic aspects of morphology and behaviour, such as cognitive ability, bioluminescence and communication, have all been found to correlate to relative eye size in groups as diverse as birds, mammals, fishes, frogs, snakes, insects, and crustaceans (Caves et al., 2017; Howell et al., 2023; Huang et al., 2022; Jones et al., 2023; Liu et al., 2023; Schweikert et al., 2022; Talarico et al., 2007).

However, almost all species studied so far have a single pair of equally sized eyes, which is just one possible configuration for the visual system. Many animals have more than two eyes, expanding the opportunity to incorporate information from multiple sources simultaneously (Buschbeck & Bok, 2023; Sumner-Rooney, 2023). For example, errant polychaete worms generally have two pairs of cephalic eyes (Purschke et al., 2006; Suschenko & Purschke, 2009), box jellyfish have four rhopalia bearing six eyes each (Nilsson et al., 2005), while some chitons have a continually growing network of eyes embedded in their shells (Chappell et al., 2023; Sigwart & Sumner-Rooney, 2021; Speiser et al., 2011). In many of these cases, eye size can vary within an individual, potentially allowing for much more complex ontogenetic, static, and phylogenetic allometric relationships, but this has not yet been examined.

Spiders provide an excellent opportunity to explore the evolution and development of eye size in such visual systems. They exhibit substantial variation in their visual morphology, ecology, and behaviour, ranging from the relatively simple to the highly sophisticated (Morehouse et al., 2017; Winsor et al., 2023). For example, jumping spiders (Salticidae) and wolf spiders (Lycosidae) use vision for pursuit hunting and complex signalling to potential mates. Most spiders have four pairs of eyes, including one pair of principal or anterior median eyes (AMEs), which usually face forward, and three pairs of secondary eyes that often cover more peripheral fields of view: the anterior lateral (ALEs), posterior median (PMEs), and posterior lateral eyes (PLEs). Their size and structure can vary dramatically both between and within species, with the most extreme adaptations expected in lineages that rely more heavily on vision (Homann, 1971; Land, 1985). Ontogenetic allometry has been studied in one salticid species, *Phidippus audax*, which exhibited slight negative allometry, in line with reports from many vertebrates (Goté et al., 2019), as well as indications from gene expression studies that eye size is established during embryonic development (Baudouin Gonzalez et al., 2023). The authors reported stronger negative allometry in the AMEs than the ALEs or PLEs, suggesting some flexibility in growth dynamics between eye pairs. In addition, the potential complexity of phylogenetic visual allometry in spiders was recently highlighted in Deinopidae: in the context of reconstructing the evolution of their enlarged PMEs, Chamberland et al. (2022) also reported lower AME diameters which, though statistically non-significant, further hint at a role for modularity and possible trade-offs in the evolution of eye size. However, this has yet to be explored from a broader comparative perspective. Intriguingly, a comprehensive study of morphological evolution recently reported no difference in total eye size between different ecological guilds across the spider tree of life (Wolff et al., 2022), despite the highly variable behavioural and visual ecology of spiders and in contrast to the aforementioned studies of other taxa.

To investigate the evolutionary dynamics of eye diameter in spiders, we compiled two large comparative datasets from museum specimens and published trait data from Wolff et al. (2022). We (i) investigated the variation in the static allometric relationships between eye and body size across eight families, selected for their variable visual ecologies and morphologies and for phylogenetic representation. Within a phylogenetic framework, we (ii) explored the relationships between eye diameter, body size, and various ecological factors, (iii) estimated the values of ancestral eye diameters and rates of evolution through ancestral state reconstruction, (iv) predicted shifts in phylogenetic allometry between eye diameter and body size, and (v) explored the evolutionary rates affecting eye size where shifts were detected.

## METHODS

### a. Specimen sampling and morphological measurements

Ethanol-preserved specimens were taken from the British Araneae collection at the Oxford University Museum of Natural History (Oxford, United Kingdom). We selected eight families that encompassed a range of visual ecologies and phylogenetic positions. For each of these, we selected five species based on the availability and size range of specimens. In total, we used 1098 individuals from Araneidae (*Araneus diadematus* Clerck, 1757, *Araniella cucurbitina (Clerck, 1757), Larinioides cornutus (Clerck, 1757), Agalenatea redii (Scopoli, 1763), Zygiella atrica* (C. L. Koch, 1845)), Gnaphosidae (*Drassodes lapidosus (Walckenaer, 1802), Gnaphosa lugubris* (C. L. Koch, 1839), *Haplodrassus signifer* (C. L. Koch, 1839), *Scotophaeus blackwalli* (Thorell, 1871), *Zelotes latreillei (Simon, 1878))*, Lycosidae (*Arctosa perita* (Latreille, 1799), *Pardosa amentata* (Clerck, 1757), *Pirata piraticus* (Clerck, 1757), *Trochosa terricola* Thorell, 1856, *Xerolycosa miniata* (C. L. Koch, 1834)), Linyphiidae (*Centromerus sylvaticus* (Blackwall, 1841), *Erigone atra* Blackwall, 1833, *Lepthyphantes minutus (Blackwall, 1833)*, *Linyphia triangularis (Clerck, 1757)*, *Walckenaeria acuminata* Blackwall, 1833), Pholcidae (*Artema atlanta* Walckenaer, 1837, *Crossopriza lyoni* (Blackwall, 1867), *Pholcus phalangioides* (Fuesslin, 1775), *Smeringopus pallidus* (Blackwall, 1858)), Salticidae (*Euophrys frontalis* (Walckenaer, 1802), *Evarcha falcata* (Clerck, 1757), *Heliophanus flavipes* (Hahn, 1832), *Mogrus mathisi* (Berland & Millot, 1941), *Neon reticulatus* (Blackwall, 1853)), Theridiidae (*Parasteatoda tepidariorum* (C. L. Koch, 1841), *Episinus angulatus* (Blackwall, 1836), *Enoplognatha ovata* (Clerck, 1757), *Theridion ludius* Simon, 1880, *Steatoda albomaculata* (De Geer, 1778)), and Thomisidae (*Diaea dorsata* (Fabricius, 1777), *Misumena vatia* (Clerck, 1757), *Ozyptila atomaria* (Panzer, 1801), *Thomisus onustus* Walckenaer, 1805, *Xysticus cristatus* (Clerck, 1757)). Lot numbers are available in Supplementary Materials S1. Between 26-31 specimens per species were selected to provide the largest range of body sizes and temporarily held in conical microtubes filled with 70% ethanol. To avoid effects of sexual dimorphism, only females were included.

Specimens were imaged using a Discovery.V12 Zeiss stereomicroscope with an attached Zeiss Axiocam 105 colour digital camera via the Zeiss software ZEN. Images of submerged specimens were taken from the dorsal view with the addition of a digital scale. Ventral and/or lateral images were taken as needed to visualise all eye pairs.

Carapace width (CW) and eye diameter (ED) were measured from scaled images using Fiji (Version 2.10/1.53c) (Schindelin et al., 2012). The CW was taken to be the width of the cephalothorax at the position of the PLEs and was used as a proxy for body size (Goté et al., 2019). If the eyes were elliptical (e.g. PMEs, Gnaphosidae), the diameter was measured along the longest axis. Measurements are available in Supplementary Materials S2.

### b. Validating specimen measurements

ED and CW measurements were preliminarily plotted to visually check for outliers, which were then remeasured. Spot-checking was also conducted on five randomly selected specimens for each species.

### c. Published dataset: Wolff et al. 2022

To provide broader phylogenetic context, we also analysed a large comparative dataset recently published by Wolff et al. (2022). The original dataset contained morphometric data (n=1968 entries) including eye diameters and carapace widths, ecological data (n=829) including hunting guilds, and a time-calibrated maximum-likelihood phylogeny of 828 species. We extracted entries with all eye diameters, carapace width, and ecological data recorded. Where duplicate entries were present for a given species, we took mean eye diameters and carapace widths. Species missing from the phylogeny were removed. The resulting dataset and phylogeny contained single entries for 474 species. Only female individuals were included in this dataset by Wolff et al. (2022), for the same reasons as above. Curated datasets are available in Supplementary Materials S3-5.

### d. Static allometry

To confirm whether the identity of the eye pair (AME, ALE, PME, or PLE) affected its diameter within our selected families, we compared eye diameters using either ANOVA or Kruskall-Wallis test and Tukey’s Honestly Significant Difference (HSD) Test or Dunn post-hoc tests between the four eye pairs within each family.

For comparison with previous allometric studies, we first performed OLS-based regression analyses to examine the linear relationship between log-transformed CW and log-transformed ED. Each species and each eye pair was analysed separately to estimate the gradient and the significance of the allometric slope. The literature is divided between the use of ordinary least squares (OLS) or standardised major axis (SMA) regression in allometry (Jürgens, 1991; Kilmer & Rodríguez, 2017; R. J. Smith, 2009; Thomas et al., 2020; Warton et al., 2006). In our dataset, the variance in eye diameter was greater than the variance in carapace width; we therefore used OLS analysis (Egset et al., 2012; Warton et al., 2006).

To examine the effect of modularity in the spider visual system, we then constructed linear mixed models for the four eye pairs in each species using the R package lme4 v1.1-30 (Bates et al., 2015). The outcome variable was the log-transformed ED. The identity of the eye pair (i.e. AME, ALE, PME, PLE), log-transformed CW, and their interaction were included as fixed effects, and the individual measured was included as a random effect. The model fitted an allometric slope to each of the four eye pairs. Species were analysed separately. We compared the full and reduced models using likelihood ratio tests and used the model that returned the lower AIC score.

To further assess the quality of fit for each allometric slope, we extracted the fitted slopes from the resultant models and calculated the differences between observed and predicted ED at each observed CW. Absolute values for these estimated residuals were pooled within families and compared between eye pairs using Kruskal-Wallis and subsequent Dunn’s tests with Holm correction for multiple pairwise comparisons.

We used the emtrends and emmeans function from the emmeans package (Lenth et al., 2023) to perform pairwise slope and intercept comparisons between the different eye pairs within species, respectively. Tukey’s adjustment is used to correct for multiple pairwise comparisons. Where the relationship between CW and ED was not significant, pairwise comparisons were ignored.

To investigate variation within families, we also constructed a second set of linear mixed models for each family, incorporating both individual and species as random effects. Slope and intercept comparisons were again performed using emtrends and emmeans, respectively. See Supplementary Materials S6-7.

### e. Phylogenetic allometry and the impact of ecological factors

Phylogenetic generalised least-squares (PGLS) regressions with maximum likelihood estimates of Pagel’s *λ* were fitted to mean log-transformed ED against mean log-transformed CW for each species in the curated dataset from Wolff et al. (2022), using the R package Caper v1.0.1 (Orme et al., 2018). Where eye pairs were absent, these species were removed from the dataset before analysis (e.g. Segestriidae with absent AMEs were removed from the AME analysis). Due to the enlargement of different eye pairs in different groups, we also ranked eye pairs by diameter within each species and ran PGLS models for the largest and smallest eye pairs. This introduces the caveat that the compared ranked eyes are therefore not always homologous. Finally, we ran PGLS on the within-species variance between the relative diameter of the four eye pairs, to examine differential investment within the modular system.

The first set of PGLS analyses used only CW as an explanatory variable. After the first run, phylogenetic outliers were identified by inspection of studentised phylogenetic residuals (*|>3|*) and iteratively removed through five runs.

To explore the impact of ecological factors on eye size, residuals from the final model were used to assess phylogenetically corrected investment in each eye pair after Thomas et al. (2020). These residuals reflect how far a given eye departs from the expectation of the global model, given carapace width and accounting for phylogeny. Positive residuals suggest a larger investment in an eye pair given body size, and vice versa. Residuals were compared between the following ecological groupings using Kruskal-Wallis tests (Thomas et al., 2020): visual vs. non-visual hunting, diurnal vs. nocturnal activity, and web-building vs. non-web building (Wolff et al., 2022). As the role of vision in hunting varies between taxa, we divided this further into visual pursuit hunting (i.e. stalking and chasing prey) and hunting with any role for vision (e.g. sit-and-wait predators that detect and capture prey using visual cues).

We also examined the impact of these ecological factors by incorporating them, and their interaction with carapace width, as covariates in a subsequent PGLS (phylogenetic ANCOVA; after Thomas et al., 2020). See Supplementary Materials S8.

### f. Identification of allometric shifts

In addition to categorising species and comparing eye allometry across ecological factors, we also used reversible-jump Markov chain Monte Carlo (rjMCMC) to implement a bivariate Ornstein-Uhlenbeck (OU) model of evolution and identify evolutionary shifts in an unbiased way, using the R package Bayou v2.0 (Uyeda et al., 2020). This allowed us to detect predicted shifts in slope and intercept of evolutionary allometric relationships across the phylogeny. We combined 10 parallel chains of 1 million iterations, each with a burn-in proportion of 0.2. MCMC chain starting points were set by drawing a number of shifts from the prior poisson distribution and randomly assigning shifts to branches randomly drawn from the phylogeny which was the probability proportional to the size of the clade descended from the branch. The output of running the chains generated a best-fit allometric model highlighting shifts in slope and/or intercept and their assigned posterior probabilities. Based on the trace output, we were confident in the convergence of our models and used the model with a 20% chance of seeing a shift. We did not accept single-branch shifts. This model was fitted for all four eye pairs. We then visualised the shifts using PGLS.plotgrade (Smaers & Mongle, 2018).

We ran a univariate OU model to estimate the regime shifts in variance between eye pairs, independent of body size, to estimate where shifts in the mean variance occur in the phylogeny. We used the function estimate_shift_configuration using a modified Bayesian information criteria (mBIC) from the package l1ou (Khabbazian et al., 2016; Tung Ho & Ané, 2014). See Supplementary Materials S9.

### g. Assessing differential changes in mean eye or body size

Under the assumption that there was, overall, a positive correlation between eye and body size, we determined whether changes in allometry were driven primarily by changes in mean eye diameter or mean body size using the phylogenetic means from the initial PGLS based on methods from Smaers et al., (2021). The descendant grades used were based on the shifts predicted by the OU models generated using Bayou. To determine whether change had occurred, we looked at the ratio: 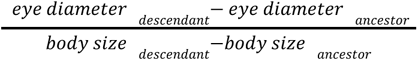 and compared this to the ancestral grade eye diameter:body size ratio (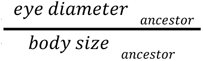). The upper and lower bounds of the 95% confidence interval of the ancestral grade body size ratio were used as a cutoff to determine if there was more of a change in the eye or body size, respectively, from the ancestor to descendant.

### h. Ancestral state reconstruction and evolutionary rate estimation

To better understand the evolution of ED and CW, we used the R function mvBM from the mvMORPH package (Clavel et al., 2015), which fits multivariate multiple rates of evolution under a BM model. We then used Bayesian MCMC using anc.Bayes from the phytools package (Revell, 2012) to sample from the posterior distribution for states at internal tree nodes.

In addition, based on the model produced by Bayou, we investigated how the rates of evolution of the predicted shifts would compare to the larger clade. Multivariate Brownian motion models were fitted to subsets of the data and phylogenetic tree for each of the four eye pairs using mvBM (Multivariate Brownian Motion) from the mvMORPH package (Clavel et al., 2015). The results included estimated branch lengths that can be used for Bayesian analysis required for ancestral state estimation using the anc.Bayes function from the phytools package (Revell, 2012). The evo.map function mvBM.getRate can then be used to calculate the rates for specific tree branches (Smaers & Mongle, 2018). This generates probability density distributions of evolutionary rates for given clades, whose overlap can be interpreted as the likelihood that they are the same. See Supplementary Materials S10.

### i. Reproducibility

All analyses were performed in Rstudio 4.2.1. Analyses and data can be found in the supplementary materials.

## RESULTS

### Eye diameter and carapace width

Measurements of 1098 individuals confirmed that eye diameter and carapace width vary between taxa and that eye diameter varies between eye pairs within many taxa. Some families (Araneidae, Gnaphosidae, Linyphiidae, Thomisidae, Theridiidae) generally have similarly sized eye pairs, others (Lycosidae, Salticidae, Pholcidae) demonstrated more variation (Figure 1A). The smallest and largest mean eye diameters for individuals were both measured from salticids: the largest was the *M. mathisi* AME, measuring 525.39 μm, and the smallest was from *N. reticulatus*, whose PME measured 39.66 μm. In terms of carapace width, linyphiids and thomisids were generally smaller than the other families (Supplementary figure SF1). The smallest species measured was the linyphiid *E. atra* with a mean carapace width of 371.97 μm, and the largest was *M. mathisi*, which measured 2280.31 μm on average. When taking carapace width into account, relative eye diameter reflected similar trends to absolute eye diameter (Figure 1B). The largest relative eye pair was found to be 0.430, the PME in the lycosid *P. piraticus*. The smallest relative eye pair was the PME of the salticid *M. mathisi*, at 0.0199.

**Figure 1.**
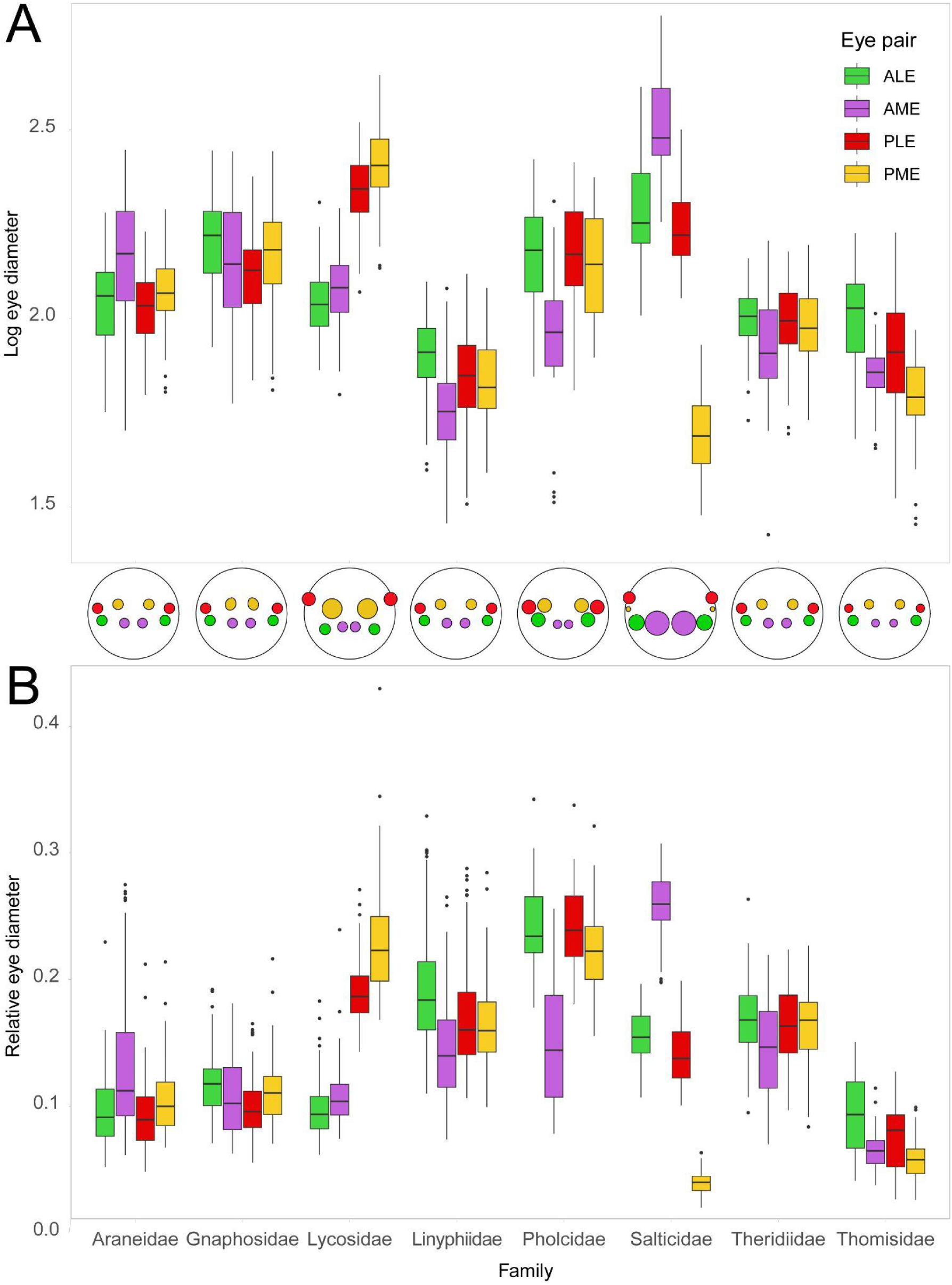
Absolute log-transformed and relative eye diameters vary substantially by family and by eye pair identity in spiders. Eye size can be relatively consistent across eye pairs (e.g. Gnaphosidae), vary by eye type (e.g. Pholcidae), or be divergent between specific eye pairs (e.g. Salticidae). Absolute (A) and relative (B) diameters exhibit similar patterns, with Linyphiidae having relatively larger eyes due to their small carapace width. Homologous eye pairs are coloured accordingly. Boxes represent the interquartile range, with solid bars at the median, and whiskers represent the lower and upper quartiles. All families include five pooled species (n=26-31 per species), except for pholcids (four pooled species, n=1-9 per species).

In Linyphiidae, ANOVA indicated that the overall effect of eye pair was statistically significant (F_3_=37.90, p<0.0001), and Tukey HSD reported significant differences between the ALEs and all other eye pairs (p_AME_<0.0001; p_PME_<0.0001; p_PLE_<0.0001), between the AMEs and the two posterior eye pairs (p_PME_<0.0001; p_PLE_<0.0001) (**Table S1**). In Lycosidae, the overall effect of eye pair was statistically significant (F_3_=591.19, p<0.0001), with significant differences between all four eye pairs (p<0.0001 (**Table S1**). ANOVA and subsequent Tukey’s HSD tests supported significant differences overall in diameter between eye pairs in Pholcidae (F_3_=8.02, p<0.0001), with the AMEs being significantly smaller than all other eye pairs (p_ALE_=0.0003; p_PME_=0.003; p_PLE_=0.0003) (**Table S1**).

Kruskal-Wallis and subsequent Dunn’s tests were used for the remaining families and showed there were significant differences between eye pairs in all families In Araneidae (χ^2^_3_=68.6, p<0.0001), the ALEs were smaller than the AMEs (Z=-6.177, p_adj_<0.0001) and the PMEs (Z=7.846, p_adj_<0.0001), while the PMEs were smaller than the AMEs (Z=4.222, p_adj_=0.0002) and larger than the PLEs (Z=-3.582, p_adj_=0.021) (**Table S2**). In Gnaphosidae (χ^2^_3_=29.8, p<0.0001) the ALEs were larger than the AMEs (Z=2.911, p_adj_=0.022) and PLEs (Z=5.318, p_adj_<0.0001) and the PMEs were larger than the PLEs (Z=-3.533, p_adj_=0.0025) (**Table S2**). In Theridiidae (χ^2^_3_=41.67, p<0.0001) the AMEs were smaller than the ALEs (Z=5.955, p_adj_<0.0001) and the PMEs (Z=-4.142, p_adj_<0.0001) and the ALEs were larger than the PMEs (Z =-5.063, p_adj_<0.0001) (**Table S2**). Finally, in Salticidae (χ^2^_3_=416.3, p<0.0001), the AMEs were larger than all other eye pairs (AME-ALE: Z=-8.136, p_adj_<0.0001; PLE-AME: Z=12.13, p_adj_<0.0001; PME-AME: Z=20.27, p_adj_<0.0001) and the PMEs were smaller than all other eye pairs (PME-ALE: Z=9.858, p_adj_<0.0001; PME-PLE: Z=10.41, p_adj_ <0.0001) (**Table S2**).

### Static Allometry

#### 1. OLS analysis

The vast majority of eye pairs, 123 out of 156 across 39 species, exhibited significant relationships with carapace width (Figure 2, **Table S3**). Of these, slope estimates ranged from 0.167 (*A. cucurbitina,* ALE) to 1.49 (*E. angulatus,* AME) (**Table S3)**. Among the eye pairs that exhibited a significant correlation to carapace width, 91 had an estimated slope significantly lower than 1 (i.e. negative allometry/hypomallometry), while 32 had a slope not significantly different from 1 (i.e. isometry), and two (*E. angulatus,* AME; *A. atlanta*, AME) had an estimated slope significantly higher than 1 (i.e. positive allometry/hyperallometry) (**Table S3, S4)**.

**Figure 2.**
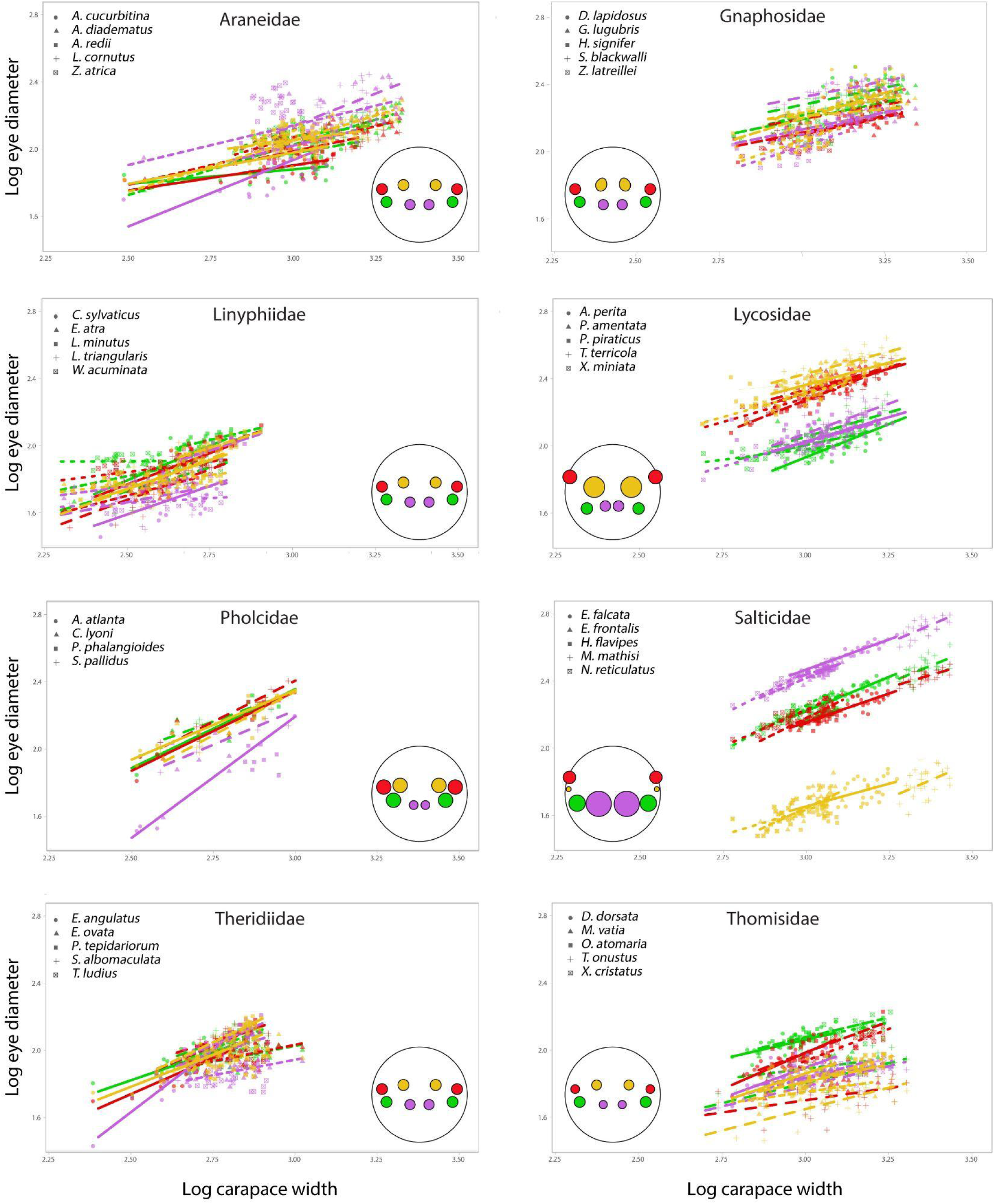
Static allometric relationships between eye diameter and carapace width in eight families of spiders. Lines indicate predicted allometric relationships where the relationship between carapace width and eye diameter is statistically significant. See Tables S3-S7 for analyses.

Although most eye pairs exhibited significant negative allometry, there was some variation in the slopes and strengths of allometric relationships across the eight families. Within Salticidae, the majority of relationships (11 out of 20) between eye diameter and carapace width were isometric, while in Lycosidae and Gnaphosidae, almost all were negatively allometric (19 and 18 out of 20, respectively) (**Table S3, S4**). Other families, particularly Theridiidae, were more variable, but no family demonstrated consistent patterns across all species. The identity of the eye pair also did not seem to consistently affect the slope, even in families that exhibit enlargement in a particular eye pair: both salticid AMEs and lycosid PMEs demonstrated variable relationships between ED and CW. R squared values generated by OLS, which give an indication of the strength of these linear relationships, varied between families ranging from 0.15 (*L. cornutus,* PLE) to 0.987 (*A. atlanta*, AME) (**Table S5**). Notably most salticids (4/6) had R squared values higher than 0.7 in both ALEs and AMEs (**Table S5**).

In some cases, the relationship between eye diameter and carapace width was found to be non-significant (33/156) (**Table S3, S4**). This occurred most commonly in Theridiidae and Gnaphosidae (6/20 eye pairs), and least commonly in Salticidae (1/20) and Thomisidae (2/20) (**Table S3, S4)**. Two species, *D. lapidosus* (Gnaphosidae) and *R. luidus* (Theridiidae), had no significant relationships between eye diameter and carapace width, and thus drove this trend in these families. The identity of the eye pairs that did not demonstrate a significant allometric relationship was roughly evenly split (AME: 8/33, ALE: 8/33, PME: 10/33, PLE: 7/33) (**Table S3, S4**).

#### 2. Mixed models

Based on linear mixed models constructed for each species, we generally, but not always, found significant relationships between carapace width and eye diameter overall. Exceptions included *A. cucurbitina* (Araneidae)*, L. cornutus* (Araneidae), *D. lapidosus* (Gnaphosidae), *Z. latreillei* (Gnaphosidae), *W. acuminata* (Linyphiidae), *P. amentata* (Lycosidae), *C. lyoni* (Pholcidae), *P. phalangioides* (Pholcidae), and *T. ludius* (Theridiidae) (**Table S6**). With the exception of *P. amentata* (Lycosidae), all visually hunting species demonstrated this significant relationship (**Table S6**).

In 26 out of 39 species, the identity of the eye pair was also found to be a significant fixed factor impacting eye diameter (**Table S6**). Of those, the interaction between carapace width and eye pair identity was significant in all except *D. lapidosus* (Gnaphosidae). See supplementary Table S6 for full model outputs.

Across these, 123/156 eye pairs had significant relationships to carapace width, of which 91/156 returned slopes <1 (negative allometry) and 33/156 returned slopes equal to 1 (isometry) (**Table S7**). Almost all eye pairs for lycosids had significant slopes that were less than one. The exceptions were *P. amentata* ALE and PLE as well as *P. piraticus*, PME which had non significant slopes. A similar pattern was observed in the thomisids, where all eye pairs and species except for *D. dorsata (*PLE) and *M. vatia* (PLE) had significant slopes less than 1 (**Table S7**). In Salticidae, allometric slopes were mostly equal to 1, with the exceptions of *E. falcata* (AME, PLE, PME) and *N. reticulatus* (PME), both of which had significant slopes less than 1, and *H. flavipes* PME, which did not exhibit a significant relationship to carapace width.

Through a series of pairwise comparisons, emtrends identified species from all eight families that demonstrated some differences between the allometric slopes of different eye pairs (i.e. p values adjusted for multiple comparisons were <0.05; **Table S8**).

Among the most common were differences in slope between the AMEs and all secondary eyes (*A. atlanta, A. cucurbitina, Z. atrica,* and *E. angulatus*), differences between the AMEs and one or both pairs of posterior eyes (*L. cornutus, L. triangularis, M. vatia, N. reticulatus, O. atomaria, S. blackwalli*), and differences between the PMEs and one or both other pairs of secondary eyes (*D. dorsata, H. signifer, E. atra, N. reticulatus, P. piraticus, S. blackwalli, X. miniata*). Individual species exhibited specific differences between one secondary eye pair and all other eyes (ALE, *X. miniata*; PME, *H. signifer*; PLE, *P. piraticus*), or between the two anterior (*P. amentata*) or the two lateral (*W. acuminata*) eye pairs.

Several species also demonstrated significant differences in overall eye diameter (i.e. intercepts) between eye pairs, as detected by emmeans. Similar to emtrends, the most common were differences in intercept between AMEs and all secondary eye pairs, which was observed in 29/39 species (*A. atlanta, A. diadematus, A. perita, A. redii, C. lyoni, C. sylvaticus, D. dorsata, D. lapidosus, E. atra, E. falcata, E. frontalis, E. ovata, G. lugubris, H. flavipes, H. signifer, L. cornutus, L. triangularis, M. mathisi, M. vatia, N. reticulatus, O. atomaria, P. amentata, P. phalangioides, P. piraticus, T. terricola, W. acuminata, X. cristatus, X. miniata, Z. atrica)* and differences in AMEs and one or both pairs of posterior eyes (10/39 species: *A. cucurbitina, E. angulatus, L. minutus, P. tepidariorum, S. albomaculata, S. blackwalli, S. pallidus, T. ludius, T. onustus, Z. latreillei)* (**Table S9)**.

#### 3. Data pooled by family

We found that carapace width was the only factor significant in predicting eye diameter across all families, while the interaction between carapace width and eye pair identity was never a significant factor in predicting eye diameter. Eye pair identity was only found to be a significant factor in thomisids. Species identity was significant in all families except araneids. The interactions between eye pair and species, and between carapace width, eye pair, and species, were significant in all families except linyphiids. A reduced model was used for the gnaphosid and salticid datasets, and full models for the other families (see Supplementary materials S6 for analysis and detailed model results). With the exception of salticid ALEs (slope =1), all other family eye pairs had significant slopes less than 1. Of these, salticid AMEs had the steepest slope (0.813), while gnaphosid PLEs had the shallowest significant slope (0.321) (**Table S10)**. When data were pooled at the family level, the patterns detected by emtrends reflected the general patterns in eye diameter observed across the families. Lycosids and salticids exhibited differences in slope between eye pairs that also differ in overall size: in Lycosidae, the slopes of the PLEs and PMEs both significantly differed from that of the ALEs (t_407_=-2.99, p=0.0156; t_403_=-2.61, p=0.046, respectively) (**Table S11)**. In the salticids, the allometric slopes of the PMEs and PLEs each differed from all other eye pairs: ALE-PLE (t_416_=3.06, p=0.0124), ALE-PME (t_417_=7.35, p<0.0001), AME-PLE (t_426_=5.01, p<0.0001), AME-PME (t_422_=9.28, p<0.0001), PLE-PME (t_422_=4.24, p=0.00016) (**Table S11)**. Among the other families, only the gnaphosids demonstrated significant differences in slope between eye pairs, between the PLEs and both the AMEs (t_400_=3.61, p=0.00195) and the PMEs (t_395_=-3.81, p=0.000937). The remaining families demonstrated no significant differences in the slopes between eye pairs (**Table S11**). With regards to intercept, almost all eye pairs were significantly different from each other in all families (**Table S12)**. However, there were a few exceptions in the pholcids, araneids, linyphiids and theridiids, where some or all of the secondary eyes were not significantly different from each other (**Table S12)**.

### Phylogenetic Allometry

PGLS reported significant relationships between eye diameter and carapace width across 458 species and 106 families (AME: slope est. 0.81±0.03, t_1,407_=25.62, R^2^ =0.616, F=656.6, p<0.0001, λ=1; ALE slope est. 0.697±0.027, t =26.2, R^2^ =0.606, F=686.7, p<0.0001, λ=0.771; PME slope est. 0.748±0.028, t =26.8, R^2^ =0.612, F=719.6, p<0.0001, λ=0.87; PLE slope est. 0.727±0.025, t_1,448_=28.72, R^2^ =0.647, p<0.0001, λ=0.756).

Both the largest and smallest eye pairs also demonstrated significant relationships with CW (Largest slope est. 0.733±0.027, t_1,443_=26.9, R^2^ =0.621, F=727.2, p<0.0001, λ=0.943; Smallest slope est. 0.79±0.027, t =29, R^2^ =0.652, F=841.7 p<0.0001, λ=0.8), as did variance (slope est. -0.0005±0.0001, t =-3.82, R^2^ =0.03, p=0.00015, λ=0.99).

#### Ecological factors

The impact of ecological factors was evaluated by comparing the relative investment in eyes in different ecological groupings. When homologous eyes were compared, visual pursuit hunters had larger AMEs (χ^2^_1_=24.3, p<0.0001), ALEs (χ^2^_1_=7.2, p=0.0071), and PLEs (χ^2^_1_=42, p<0.0001), but no difference in PME diameter (Figure 3). Overall, visual hunters also exhibited larger AMEs (χ^2^_1_=5.82, p=0.016), ALEs (χ^2^_1_=7.04, p=0.008) and PLEs (χ^2^_1_=22, p<0.0001), but smaller PMEs (χ^2^_1_=3.91, p=0.048). Web-building species had larger PMEs (χ^2^_1_=7.99, p=0.0047) but did not exhibit a clear difference in AME, ALE, or PLE diameter (Figure 4). Diurnal species generally had smaller PMEs (χ^2^_1_=15.43, p<0.0001; Figure 5), but the diameter of AMEs, ALEs, and PLEs were unaffected.

**Figure 3.**
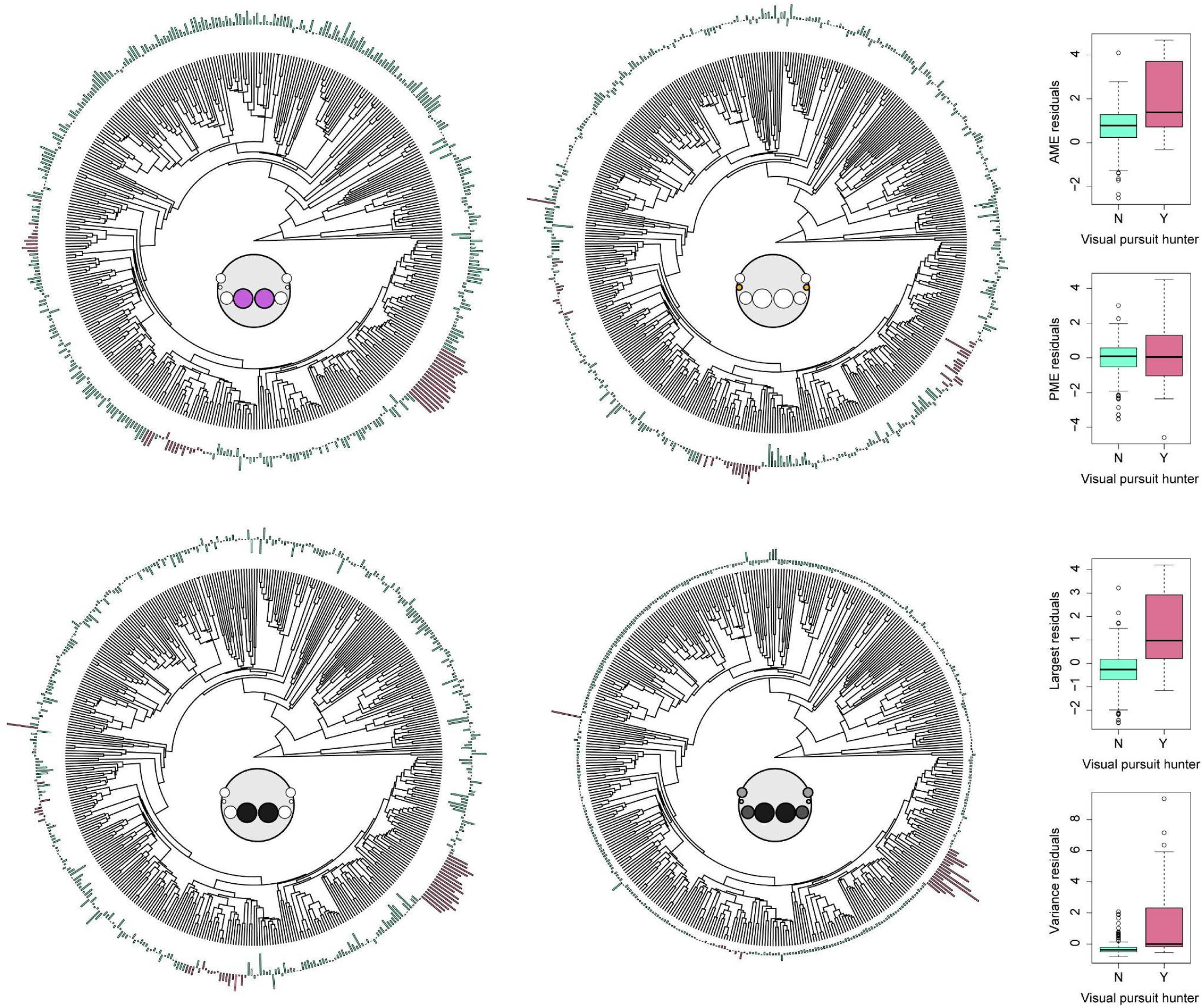
Visual pursuit hunting in spiders is associated with larger AMEs, greater enlargement of the largest eye pair, and increased variance in eye diameter, but does not affect PME diameter. Residuals from PGLS models, plotted around phylogenies, demonstrate that investment in certain homologous and ranked eye pairs is higher in visual pursuit hunters (pink bars) than other spiders (green bars) given their carapace width. Clockwise from top left: AMEs, PMEs, variance between eye pairs, and largest ranked eye pair. Box plots indicate residuals compared between visual pursuit hunters and other taxa, which were compared using Kruskal Wallis tests. See Supplementary Materials S8.

**Figure 4.**
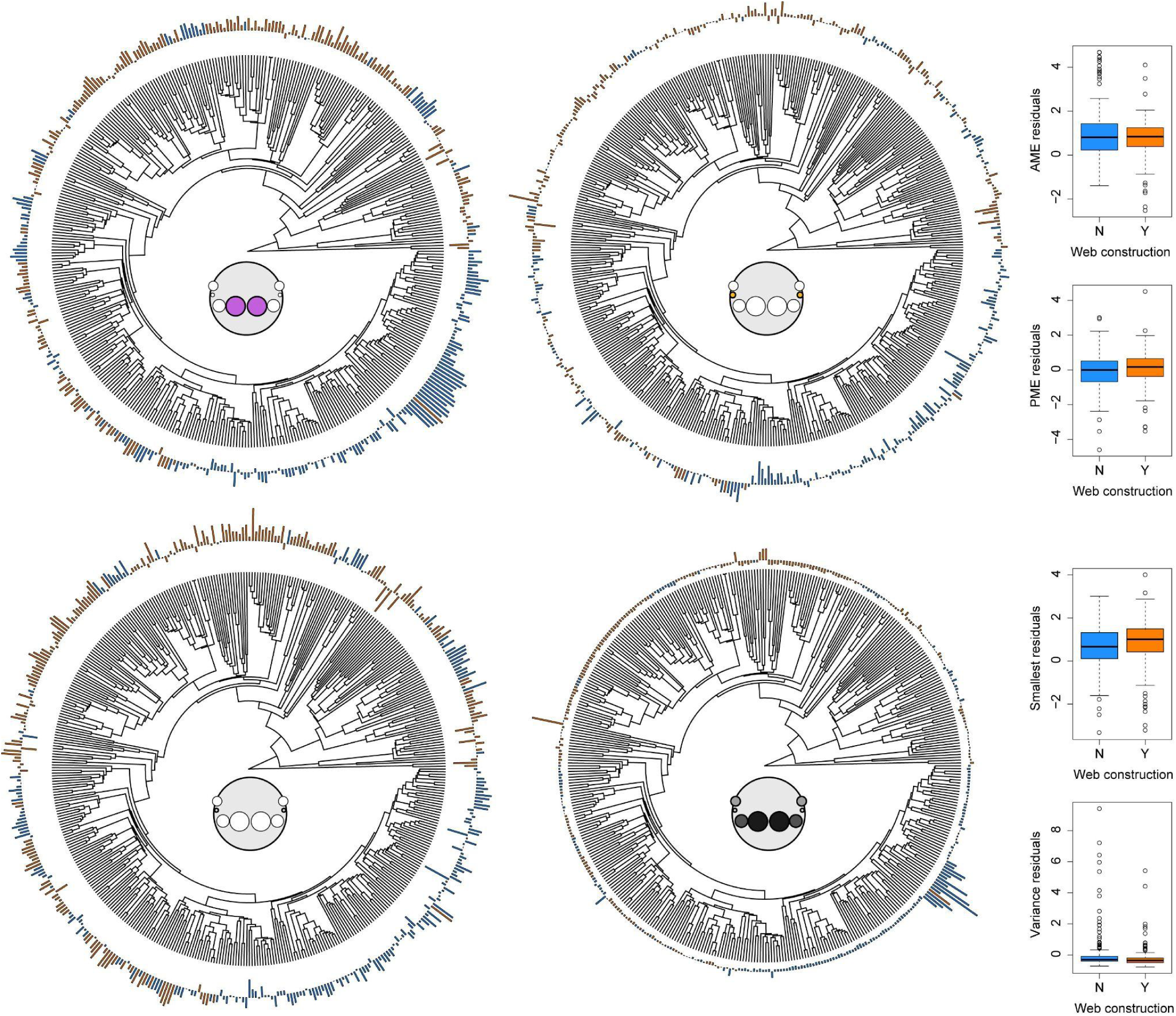
Web construction in spiders is associated with larger PMEs, greater investment in the smallest ranked eye pair, and decreased variance in eye diameter, but does not affect AME diameter. Residuals from PGLS models demonstrate that investment in certain homologous and ranked eye pairs is higher in web-building species (orange bars) than other spiders (blue bars) given their carapace width. Clockwise from top left: AMEs, PMEs, variance between eye pairs, and smallest ranked eye pair. Box plots indicate residuals compared between web builders and other taxa, which were compared using Kruskal Wallis tests. See Supplementary Materials S8.

**Figure 5.**
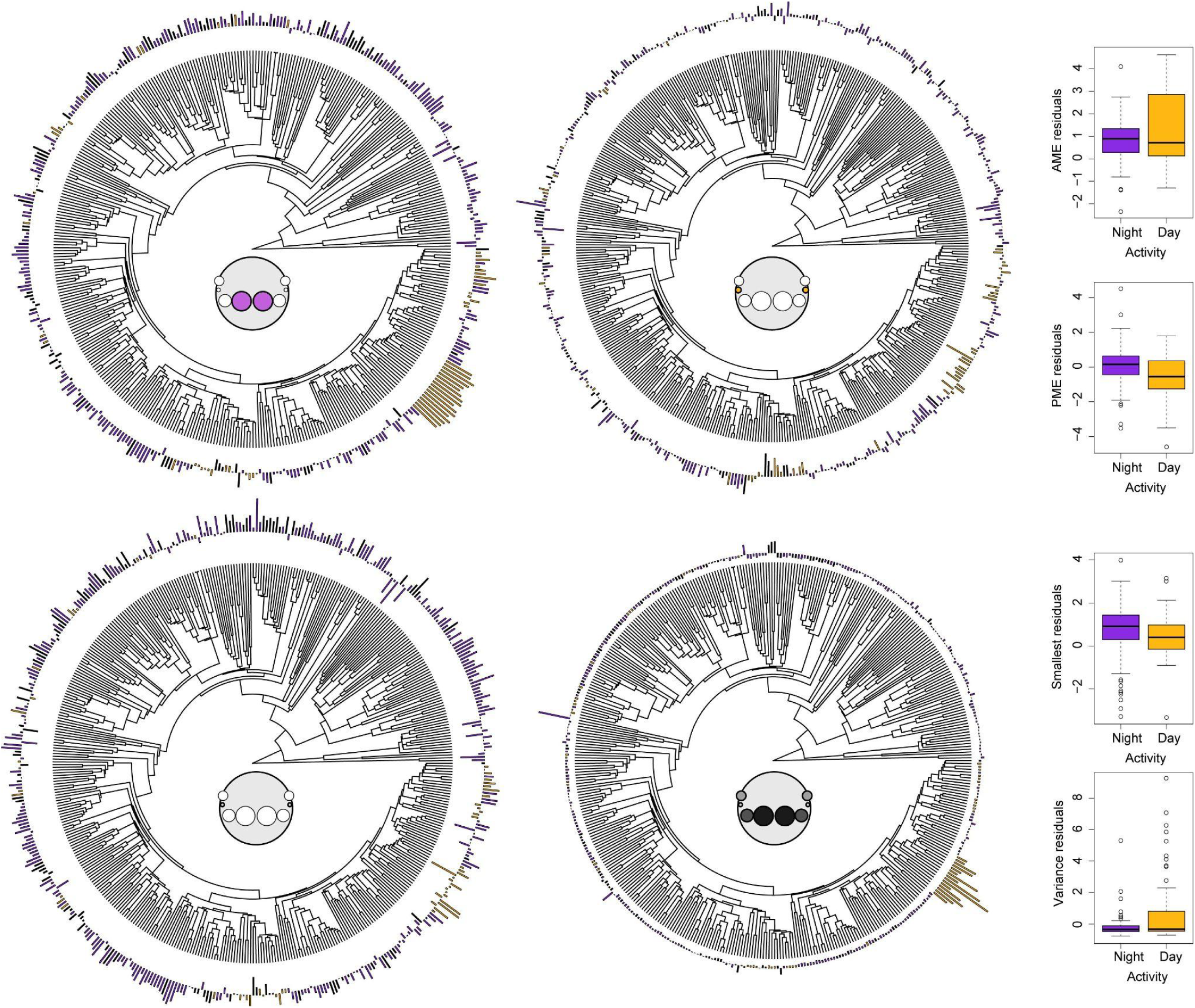
Diurnal activity in spiders is associated with greater variance in eye diameter, but decreased PME diameter and investment in the smallest eye pair. Residuals from PGLS models demonstrate that investment in certain homologous and ranked eye pairs differs between diurnally active species (yellow bars) and nocturnally active species (purple bars) given their carapace width. Clockwise from top left: AMEs, PMEs, variance between eye pairs, and smallest ranked eye pair. Box plots indicate residuals compared between nocturnal and diurnal taxa, which were compared using Kruskal Wallis tests. See Supplementary Materials S8.

When eye pairs were ranked, the diameter of the largest eye pair was significantly greater in visual hunters (visual pursuit: χ^2^_1_=51.91, p<0.0001; all visual: χ^2^_1_=31.82, p<0.0001; Figure 3), but was not affected by web building or activity (Figure 4). The diameter of the smallest eye pair was smaller in visual hunters (χ^2^_1_=11.25, p=0.0008), larger in web builders (χ^2^_1_=11, p=0.0009), and larger in nocturnal species (χ^2^_1_=11.81, p=0.0006; Figure 5). Variance between the eye pairs was significantly greater in visual hunters (visual pursuit: χ^2^ =59.7, p<0.0001; all visual: χ^2^ =50.9, p<0.0001; Figure 3) and significantly lower in web-building species (χ^2^ =13.13, p=0.0003; Figure 4), but unaffected by activity (Figure 5).

When these ecological factors were included in the PGLS model as additional explanatory parameters (phylogenetic ANCOVA), we found a significant effect of visual pursuit hunting on AME (t_3,405_=4.66, p<0.0001), ALE (t_3,443_=4.15, p<0.0001), PLE (t_3,452_=5.1, p<0.0001), and largest eye (t_3,441_=7.9, p<0.0001) diameters, and on variance (t_3,436_=10.47, p<0.0001). When all visual hunters were included we found significant effects of visual hunting on the AME (t_3,405_=2.9, p<0.0001), ALE (t_3,443_=4.16, p<0.0001), PME (t_3,447_=-2.7, p=0.0066), PLE (t_3,446_=2.7, p=0.0072), the largest (t_3,441_=6.45, p<0.0001) and smallest (t_3,445_=-2.48, p=0.013) eye diameters, and on variance (t_3,436_=9.51, p<0.0001). Diurnal and nocturnal activity had a significant effect on the diameter of the PMEs (t_3,344_=-2.31, p=0.021) and on variance (t_3,337_=4.01, p<0.0001). Web construction was not a significant factor.

##### Prediction of allometric shifts

The model detected multiple allometric shifts in all four eye pairs, in both the intercept of the allometric relationship (theta), and the slope (beta) (Figure 6). Full model outputs can be found in Supplementary Materials S9.

**Figure 6.**
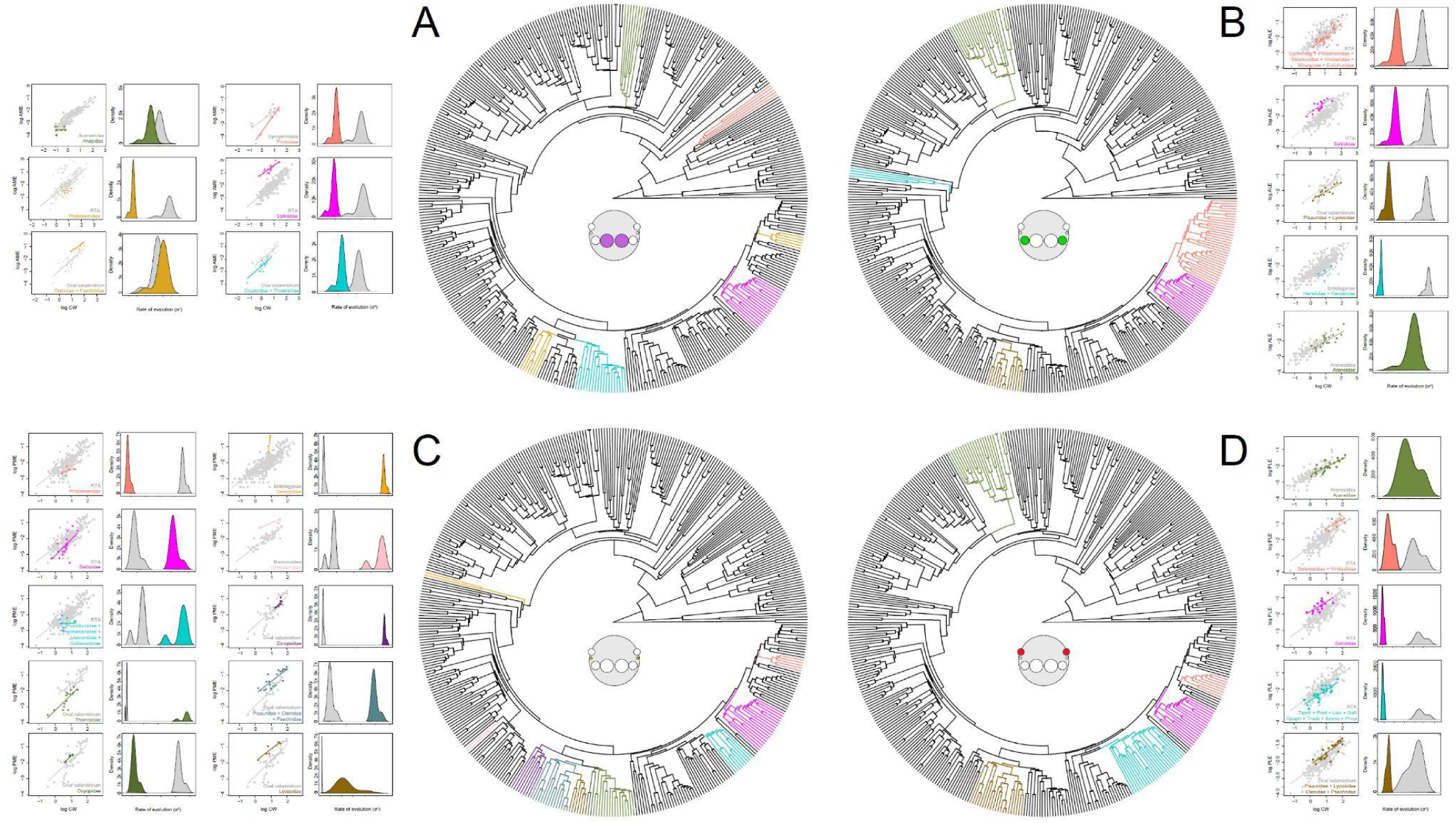
Predicted shifts in eye allometry occur in several clades, with rates of evolution often increasing or decreasing following a shift. Coloured branches indicate clades where a shift is predicted in the allometric relationship between eye diameter and carapace width. Adjacent graphs in corresponding colours depict the shifted allometries compared to the global model (scatter plots) and the predicted shift to evolutionary rate within that clade compared to its ancestral grade (probability density histograms). See Supplementary Materials S9.

In the AMEs (Figure 6A), decreases in intercept were predicted in Anapidae, Philodromidae, Pholcidae, and (Thomisidae+Oxyopidae), while increases in intercept were predicted in (Ctenidae+Psechridae) and Salticidae. Decreases in slope were predicted in Anapidae and Philodromidae, and increases in slope were predicted in (Ctenidae+Psechridae), Pholcidae, Salticidae, and (Thomisidae+Oxyopidae). In the ALEs (Figure 6B), decreases in the intercept were predicted in (Corinnidae+Philodromidae+Xenoctenidae+Selenopidae+Viridasiidae+Miturgidae+Eutichuridae), Araneidae, (Hersiliidae+Oecobiidae), and Lycosidae, and an increase in the intercept was predicted in Salticidae. In the PMEs (Figure 6C), the intercept was predicted to decrease in Philodromidae, (Prodidomidae+Trochanteriidae+Liocranidae+Gallieniellidae+Phrurolithidae), Salticidae, and Thomisidae. It was predicted to increase in Cyclotenidae, Deinopidae, Lycosidae, Oxyopidae, (Pisauridae+Ctenidae+Psechridae), and (Zoropsidae+Udubidae). Decreases in slope were predicted in Philodromidae and in (Prodidomidae+Trochanteriidae+Liocranidae+Gallieniellidae+Phrurolithidae), and increases were predicted in Cyclotenidae, Deinopidae, Lycosidae, Oxyopidae, (Pisauridae+Ctenidae+Psechridae), Salticidae, Thomisidae, and (Zoropsidae+Udubidae). Finally, in the PLEs (Figure 6D), a decrease in the intercept was predicted in (Prodidomidae+Trochanteriidae+Liocranidae+Gallieniellidae+Phrurolithidae+Gnaphosidae+Trochanteriidae+Lamponi dae+Trachelidae+Cithaeronidae+Ammoxenidae), and increases were predicted in (Lycosidae+Pisauridae+Ctenidae+Psechridae), Salticidae, and (Selenopidae+Viridasiidae). Decreased slope was predicted in Araneidae and in (Prodidomidae+Trochanteriidae+Liocranidae+Gallieniellidae+Phrurolithidae+Gnaphosidae+Trochanteriidae+Lamponi dae+Trachelidae+Cithaeronidae+Ammoxenidae), and increased slope was predicted in (Lycosidae+Pisauridae+Ctenidae+Psechridae) and (Selenopidae+Viridasiidae).

###### Differential changes in mean eye or body size

To assess whether these predicted allometric shifts were driven by changes to eye diameter or changes to carapace width, we examined the difference in the ratios of ED and CW within shifted clades, compared to their ancestral grades. We found that the allometric shifts predicted in the AMEs of Pholcidae, Thomisidae+Oxyopidae, Anapidae, and Salticidae were the result of greater change in mean AME diameter, rather than in mean CW, compared to their respective ancestral grades (**Table S13)**. Shifts in the allometry of salticid and lycosid ALEs were attributable to change in ALE diameter rather than mean CW, while those in Corinnidae+Miturgidae, Araneidae, and Hersiliidae+Oecobiidae resulted more from changes in mean CW (**Table S13)**. Thomisidae, Pisauridae+Psechridae, Cyclotenidae, and Deinopidae all exhibited greater change in PME size than in mean CW, while Oxyopidae and Lycosidae exhibited more change in mean CW than in PME diameter (**Table S13)**. The changing allometry of the PLEs in Selenopidae+Viridasiidae, Salticidae, Prodidomidae+Trochanteriidae, and (Lycosidae+Pisauridae+Ctenidae+Psechridae) was attributed to PLE diameter rather than CW, while Araneidae exhibited greater change in mean CW than in PLE diameter (**Table S13)**.

###### Ancestral state reconstructions

Ancestral state reconstructions were performed for both log-transformed absolute and for relative ED (Supplementary Materials S10, Supplementary Figures SF11-SF14). The common ancestor of all spiders was predicted to have smaller AMEs than secondary eyes; reconstructions of log-transformed eye diameters indicated that the secondary eyes were of similar size, but reconstructions of relative eye diameter suggest that the posterior pairs were smaller than the ALEs (**Table 1)**. Reconstructions of logAME evolution showed that AME diameter at the base of Mygalomorphae was higher than in the common spider ancestor, while the main early-diverging araneomorph clades, Synspermiata and Entelegynae, exhibited smaller AMEs (**Table 1**). Within araneomorphs, logAME decreased further at the base of the Araneoidea, but gradually increased through the more derived RTA, marronoid, and oval calamistrum clades (**Table 1**). The log-transformed diameters of the three secondary eye pairs all followed a similar pattern, remaining stable at the base of Mygalomorphae, but decreasing in all araneomorph clades, particularly in the araneoid ancestor, but recovering in the oval calamistrum clade (**Table 1**). The reconstruction of relative eye diameters indicated that AME size was stable at the base of Mygalomorphae compared to the common spider ancestor, and that they became relatively larger in Synspermiata and Entelegynae, increasing further in araneoids but decreasing again slightly in the RTA, marronoid, and oval calamistrum clades. The relative diameter of all three secondary eye pairs decreased in mygalomorphs but retained the rough proportions between them of the spider ancestor, with the posterior eyes being smaller than the ALEs. At the bases of the earlier-diverging araneomorph clades (Synspermiata, Entelegynae) and in Araneoidea, all three pairs increased and equalised in their relative size. However, this pattern does not persist in the ancestors of the marronoid, RTA, and oval calamistrum clades, wherein different pairs are enlarged and reduced compared to the common spider ancestor (**Table 1**).

**Table 1:**
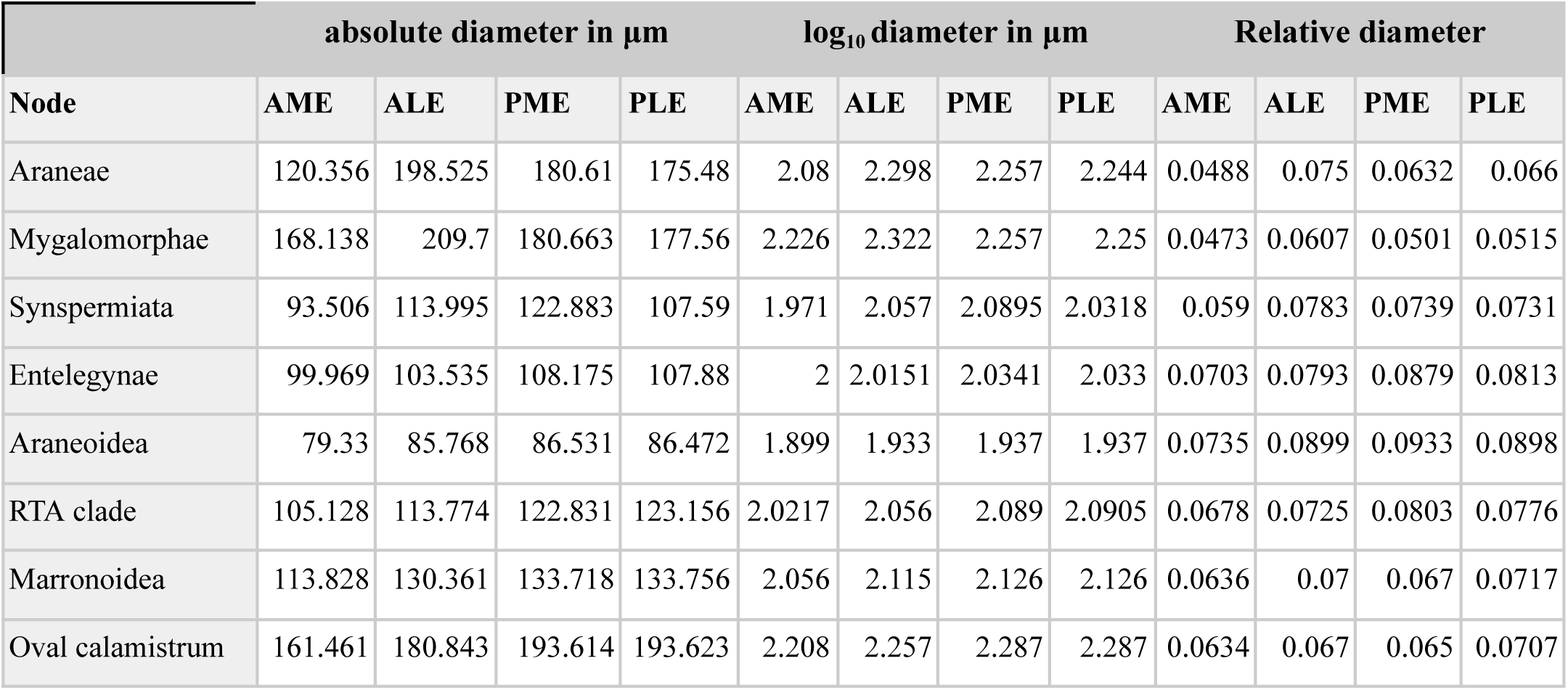
Reconstructed ancestral eye diameters for major clades. See Supplementary Materials S10, Supplementary Figures SF11-14, and Supplementary Table S14.

At finer resolution, ancestral state reconstructions for log-transformed eye diameters were largely congruent with the allometric shifts predicted by Bayou. Notable increases in logAME occurred within Salticidae, Ctenidae+Psechridae+Zoropsidae+Udubidae, Theraphosidae, and Sparassidae, and decreases in Linyphiidae, Anapidae+Mysmenidae, and Theridiosomatidae (Supplementary Figure SF11A). Increased logALE diameter was most apparent in the Salticidae, Ctenidae+Psechridae+Zoropsidae+Udubidae and mygalomorphs, while logALE was reduced in Anapidae and Theridiidae (SF12A). Increases in the log-transformed PME diameter were apparent in Lycosidae, Ctenidae, Psechridae, Zoropsidae, and Udubidae, while this decreased in Anapidae, Symphytognathidae, Mysmenidae, and Oonopidae (SF13A). The log-transformed PLE data showed the Ctenidae, Psechridae, Zoropsidae, and Udubidae growing larger and Anapidae having smaller PLEs (SF14A). Reconstructions of relative eye diameter revealed a marked increase in salticid AMEs (Supplementary Figure SF11B), in the ALEs of Salticidae and (Anapidae+Symphytognathidae+Mysmenidae+Theridiosomatidae)(SF12B), in the PMEs of Lycosidae, Deinopidae, and Symphytognathidae (SF13B), and in the PLEs of Salticidae, Uloboridae, and Anapidae (SF14B).

###### Rates of Evolution

The rates of eye diameter evolution were estimated for clades exhibiting allometric shifts and their ancestral grades using a multivariate Brownian motion mode with MCMC sampling. The reported rate is the mean of the probability distribution. In the AMEs, Salticidae (0.00214), Philodromidae (0.00191), Thomisidae+Oxyopidae (0.00181), and Pholcidae (0.00180) had lower rates of evolution compared to their ancestral grades (**Table S15)**. Corinnidae+Philodromidae+Xenoctenidae+Selenopidae+Viridasiidae+Miturgidae+Eutichuridae (0.00147), Salticidae (0.00145), Lycosidae (0.00145), and Hersiilidae (0.00285) all had lower rates of ALE diameter evolution compared to their ancestral grade. While Philodromidae (0.00132), Oxyopidae (0.00128) and Cyclotenidae (0.00538) had lower rates of PME evolution compared to their ancestral grades, Salticidae (0.001814), (Prodidomidae+Trochanteriidae+Liocranidae+Gallieniellidae+Phrurolithidae) (0.00245), Thomisidae (0.00821), Lycosidae (0.00540), (Pisauridae+Ctenidae+Psechridae) (0.00164), (Zoropsidae+Udubidae) (0.005), and Deinopidae (0.00197) had higher rates. Selenopidae + Viridasiidae (0.00372), Salticidae (0.00143), (Lycosidae+Pisauridae+Ctenidae+Psechridae) (0.00125), and (Prodidomidae+Trochanteriidae+Liocranidae+Gallieniellidae+Phrurolithidae+Gnaphosidae+Trochanteriidae+Lamponi dae+Trachelidae+Cithaeronidae+Ammoxenidae) (0.00123) all had lower rates of evolution in their PLEs compared to ancestral grades (**Table S15)**.

## Discussion

The factors influencing eye diameter in spiders are numerous and complex. Overall, eyes tend to scale hypoallometrically with carapace width, in terms of both static and phylogenetic allometry, broadly in line with patterns observed in other taxa. However, the relationships of different eye pairs to carapace width, ecological factors, and to one another can vary substantially both within and between species, families, and larger clades. Additionally, the evolutionary lability of selected taxonomic groups and eye pairs appears to vary; for example, salticids exhibited multiple allometric shifts with changing evolutionary rates and substantial differences between the four eye pairs, while across all spiders, the PMEs demonstrated the most variable static and phylogenetic allometries and evolutionary dynamics.

### Static Allometry

#### Growth dynamics vary between eye pairs, species, and families

Eye diameter and carapace width vary greatly, at both the species and the family level. This provides the basis for the observed diversity in allometric relationships found in different families and reflects the flexibility in developmental dynamics in eye development: we observed substantial variation in the elevation, gradient, and fit of allometric slopes across species and eye pairs. In addition, we had also found variation in the scaling relationships between eye pair and body size between families. As expected, visual hunters like salticids and lycosids had larger eyes than aerial web hunters such as araneids and theridiids - facilitating their well-documented contrast sensitivity and spatial resolution (Land, 1985). However, this only applied to certain eye pairs, whose identity differed between the families. Other eye pairs were more typically sized or even smaller than those of aerial web hunters, indicating that there are still limitations to total investment in the visual system in these groups. This resulted in greater variation in eye diameter in lycosids and salticids. By contrast, thomisids exhibited rather small eyes, despite their vision contributing to prey capture and habitat selection (Bhaskara et al., 2009; Insausti et al., 2012; Théry, 2007). Generally speaking, patterns were consistent between absolute and relative eye diameters, with the exception of Linyphiidae, which had very small eyes in absolute terms, but larger relative eye diameters owing to their small size overall. This may reflect size-related optical constraints or a minimum useful diameter that must be maintained even in smaller animals.

Although the variability of eye diameters is well-known in spiders, these data are among the first to test their allometric relationships directly. The correlation between eye diameter and carapace width exhibits substantial variability, including isometric, negative, and positive allometric relationships. While there is no consistent link between the slope estimates and ecology or eye type, salticid eyes are more likely to exhibit static isometry than other families (**Table S4**). Interestingly, this is at odds with a study of ontogenetic visual allometry in the jumping spider *Phidippus audax* by (Goté et al., 2019), which demonstrated that the lenses exhibited negative allometry through successive instars. Flexibility in growth dynamics may make an important contribution to adult eye size, which directly affects visual capability. Similar variability in eye allometry has been noted in other animals including anurans (Shrimpton et al., 2021) and deep-sea shrimp (Schweikert et al., 2022). However, the modular nature of the spider visual system provides an additional dimension for variation, and therefore, specialisation. In their study of *P. audax*, Goté et al. (2019) found that allometric slopes differed between the AMEs, ALEs, and PLEs. We find that this allometric divergence between eye pairs is not unique to salticids, but occurs in species across all eight families studied. In some cases, this manifests between the two eye types, principal and secondary; given the distinct developmental origins of these two types, their growth dynamics could be simply uncoupled or innately independent (Samadi et al., 2015; Schomburg et al., 2015). However, differences between the secondary eye pairs are potentially more complex. This seems to occur more commonly, but not exclusively, in groups such as lycosids and salticids that hunt outside an aerial web and may have more diverse visual needs including navigation, prey identification and capture, and courtship.

#### Visual hunters exhibit tighter relationships between their largest eye pairs and body size

While the majority of spider eyes demonstrated a significant correlation to carapace width, the strength of this relationship varied. For example, the residuals between fitted allometric relationships and the observed data were generally much higher in Araneidae than Salticidae (Supplementary Figure SF2). This variation is even apparent within families and species. In salticids, the relationships between AME, ALE, and PLE diameters and carapace width are very tight, with low residuals, while that between PME diameter and carapace width is much looser (Figure 2, SF2). This may reflect variable selective constraint on eye diameter; the AMEs are highly specialised for spatial acuity and play a critical role in prey tracking and visual communication, while the ALEs and PLEs are crucial for detecting movement and directing the gaze of the AMEs (Jakob et al., 2018; Land, 1985; Zurek et al., 2015; Zurek & Nelson, 2012). Their size is likely to be under strong selective pressure to sustain these visual capabilities. By contrast, the PMEs are often described as vestigial and probably contribute relatively little to visual behaviour, potentially being subject to lesser selective pressure and thus facilitating increased variation, as reflected by their increased evolutionary rates (Figure 6). In some cases, the relationships between eye diameter and carapace width were not significant; while this occurred in all families studied, it was most common in theridiids (aerial web hunters) and gnaphosids (wandering hunters), and least common in salticids (visual pursuit hunters) and thomisids (sit-and-wait visual hunters), further indicating that allometry is subject to tighter control in taxa more reliant on vision.

### Phylogenetic Allometry

#### Several ecological factors affect eye size and visual system configuration

##### Visual hunters have larger eyes and greater variation in eye size

Eye diameter can limit both acuity and sensitivity; larger eyes are likely to have a longer focal length and more photoreceptors, as well as larger apertures, allowing more light into the eye (Kiltie, 2000; Ross & Kirk, 2007; Veilleux & Kirk, 2014). Increased spatial resolution and contrast sensitivity are potentially beneficial to species relying on visual cues to identify, pursue, and capture prey, and increased eye size has been found in visually driven and active predators within many other groups, including fish (Caves et al., 2017), elasmobranchs (Lisney & Collin, 2007), birds (Burton, 2008; Garamszegi et al., 2002), mammals (Veilleux & Kirk, 2014) (and within primates (Ross & Kirk, 2007)), and amphipods (Glazier & Deptola, 2011). Indeed, our analyses reported larger AMEs, ALEs, and PLEs in visual hunters than in other taxa, and increases in eye diameter were detected in families known to use visual cues including Salticidae, Lycosidae, Deinopidae, Oxyopidae, Ctenidae (Fenk et al., 2010), and Pisauridae (Perevozkin et al., 2004). Interestingly, this unbiased approach also predicted increases in groups such as Cycloctenidae, Viridasiidae, Selenopidae, and Psechridae, which are not explicitly known for using vision in hunting. However, these sit within the RTA clade alongside many of the aforementioned families, and are mostly fast-moving, cursorial hunters. In several cases, these predicted allometric shifts were independent of their nearest known visually hunting relatives. These groups may represent exciting new avenues for research into uses of vision in spiders. By contrast, Thomisidae, which use vision to (at least) select hunting sites and mediate colour-changing behaviour (Bhaskara et al., 2009; Insausti et al., 2012), exhibited decreased eye size in both pairs of median eyes (Figure 6). The regular organisation of the retinae hints at other possible roles for vision in this group, but these species use a diurnal sit-and-wait strategy instead of wandering to forage (Bhaskara et al., 2009; Insausti et al., 2012). This distinction may reduce their need for higher acuity or sensitivity compared to visual pursuit predators. Conversely, members of the RTA clade that wander to hunt, but have not been observed to use vision in prey detection or capture, exhibited allometric shifts reducing the size of selected eye pairs, indicating that any contributions of vision to the wandering lifestyle also do not necessitate increased eye diameter.

Unlike these other groups, the spider visual system incorporates an additional layer of complexity due to its modular nature. When this is taken into consideration, the relationship between eye size and hunting behaviour becomes stronger and more complex. First, the largest eye pair (regardless of its identity) is significantly greater in visual hunters. Due to different groups investing in different eye pairs, this effect is dulled when only homologous pairs are compared - for example, the very largest eye diameters were the salticid AMEs and deinopid PMEs, which both play a critical role in prey capture. However, the salticid PMEs and deinopid AMEs are also dramatically reduced. Similarly, alongside increased diameter in selected eye pairs, decreases in other pairs were also predicted in Lycosidae, Pisauridae, and Oxyopidae (Figure 6). This highlights another key finding: visual hunters do not simply have larger eyes, they also exhibit greater variance in eye diameter. This may reflect limitations on overall investment in the visual system, wherein some eye pairs are enlarged, but not all; indeed, some analyses found that the smallest ranked eye pair was smaller in visual hunters than in other species, potentially indicating a trade-off in energy or space. Size variation between eye pairs is also likely to reflect functional divergence between them, which could result from the more diverse needs of visual hunters.

##### Web builders exhibit lower variance between eye pairs

Species hunting in aerial webs are likely to rely heavily on mechano- and chemoreception for prey identification and capture, courtship, and the evasion of threats (Barth, 1985, 2002). Vision may still play a role in the detection of approaching predators, habitat selection, and regulating activity, but these are relatively simple tasks that do not require high acuity or sensitivity, or regional specialisation (Winsor et al., 2023). Indeed, sheet-web building spiders are thought to have poor visual acuity (Clemente et al., 2010). Additionally, spiders occupying aerial webs may be at lower risk of predation than wandering spiders (Gunnarsson & Wiklander, 2015), and previous studies of mammals, birds, and crustaceans have found that animals subject to greater threat of predation are likely to invest more in vision (Beston et al., 2019; Glazier & Deptola, 2011; Møller & Erritzøe, 2010; Smith & Litvaitis, 1999). Combined with the energetic cost of eyes and visual processing in the central nervous system, we anticipated (like Wolff et al. 2022) that web builders would have smaller eyes. In fact, we found that web-building spiders had significantly larger PMEs than other species, and that their smallest ranked eye pair was larger than non-building species. However, these may be explained by the finding that web-building species exhibited less variance between eye pairs. If the visual system mainly fulfils tasks such as shadow detection and monitoring ambient light, there may be little need for elaboration or regional specialisation, and therefore differential investment in different eyes is not necessary. Conversely, uniform function and size may facilitate greater total field of view coverage through the arrangement of the eyes, which maximises the range of detection for such non-directional tasks (Buschbeck & Bok, 2023; Winsor et al., 2023). This may be particularly important to exposed aerial web hunters Such uniformity is easily innately achieved among the secondary eyes in particular, given their shared primordia, and almost certainly predates the innovation of aerial web hunting (Blackledge et al., 2009; Fernández et al., 2018; Wolff et al., 2019). Shifts reducing the size and allometric slope of specific eye pairs were, however, detected in the web-building Araneidae (ALEs and PLEs), Pholcidae (AMEs), and Anapidae (AMEs). The shifts in araneids were driven by increases in carapace width that were unmatched by corresponding increases in eye size. This suggests that there is no advantage to maintaining their relative diameter as body size increases, and that the visual needs met by these eye pairs are not sufficiently complex or important to merit the investment. The reduction in AME diameter detected in Anapidae and Pholcidae, by contrast, was driven by changing eye size and exhibited lower subsequent evolutionary rates than the surrounding lineages, which could reflect selective pressure to reduce the cost of the visual system in these groups.

##### Nocturnal spiders had significantly larger PMEs and minimum eye sizes than diurnal spiders

Greater eye aperture (pupil) diameter increases achievable contrast sensitivity, and many species that use vision in low-light environments exhibit larger eyes, including crustaceans (Howell et al., 2023), fish, insects (Freelance et al., 2021; Tierney et al., 2017; Warrant, 2008), geckos (Schmitz & Higham, 2018), frogs (Thomas et al., 2020), primates (Ross & Kirk, 2007), and birds (Brooke et al., 1999; Garamszegi et al., 2002; Hall & Ross, 2007; Liu et al., 2023). In spiders, the secondary eyes are well suited for low-light vision thanks to the reflective tapetum enabling improved photon capture and enhanced sensitivity, so the enlargement of the PMEs, as the only medially placed secondary eyes, is logical for species that rely on visual tasks at night. This is most strikingly apparent in Deinopidae, where the high predicted rate of evolution in the PMEs (Figure 6) results from the secondary reduction of PME diameter in *Menneus* (Chamberland et al., 2022). Despite speculation that this might enable niche separation between nocturnal *Deinopis* and crepuscular *Menneus*, both have been observed foraging at night (Chamberland et al., 2022). Additional structural changes to the eye further support lower sensitivity in *Menneus* PMEs (Blest et al., 1980), indicating a reduced contribution of vision to hunting in this genus. Other groups with enlarged PMEs (according to predicted allometric shifts, Figure 6, and ancestral state reconstructions, SF11-14) include lycosids, ctenids, and pisaurids, many - but not all - of which are nocturnal visual hunters (Suter & Benson, 2014), as well as oxyopids, which are diurnal but close relatives of the aforementioned families. It seems that a visual and cursorial element to hunting also contributes to this pattern, as allometric shifts increasing PME diameter were not predicted in nocturnal aerial web or wandering hunters such as gnaphosids or dysderids.

The higher minimum eye size among nocturnal species may arise from two factors. The first is a by-product of low variance in eye diameter that, as for web-builders, may reflect simplicity and uniformity in visual needs and capabilities in nocturnal spiders. The second is that the minimum aperture diameter required for an eye to achieve a given optical sensitivity is larger in low light (Frederiksen & Warrant, 2008). Thus, even if the smallest eye is not responsible for complex visual tasks, as the PMEs are in the deinopids and lycosids, its lower size limit to remain useful is likely to be higher than in diurnal species.

Conversely, nocturnal and other dark-living animals may limit their reliance on, and investment in, vision in favour of other sensory modes. This has been described in a range of taxa including snakes (Liu et al., 2012) and bats (Thiagavel et al., 2018), which use chemo- and thermoreception and echolocation, respectively, instead of vision to hunt and navigate. We did not detect evidence of this occurring in nocturnal species, although the reduction and loss of eyes has been documented in diverse lineages and habitats (Mammola & Isaia, 2017). Ancestral spiders were probably night-active and mainly reliant on non-visual cues already, as are modern mygalomorphs; visually driven hunting is likely to be a highly derived trait and perhaps more likely to have evolved in diurnal species where intensity is less likely to limit resolution and sensitivity.

#### Additional factors may contribute to the evolution of eye size

Besides visual hunting, web construction, and daily activity patterns, there are, of course, other factors that may interact with eye size and visual ecology that have not been tested here. For example, Leuckert’s law, which predicts that fast-moving animals will have larger eyes, has been discussed in mammals, birds, and fish, with mixed conclusions (Brooke et al., 1999; Caves et al., 2017; Hall & Heesy, 2011; Heard-Booth & Kirk, 2012). Although data are scant on the speeds of most spiders, some notably fast groups are among those that exhibit enlarged eyes; for example, both Cycloctenidae and Selenopidae are known for fast striking behaviour and high manoeuvrability, but not for their visual abilities (Kelly et al., 2023; Zeng & Crews, 2018). Nonetheless, increases in PME and PLE diameter were predicted in these families, respectively (Figure 6).

In addition, habitat structure and light conditions can exert strong selective pressures over visual system structure and function. These have been found to affect eye size in frogs (Thomas et al., 2020), fishes (Caves et al., 2017; Dobberfuhl et al., 2005; Ryer, 1988), birds (Jones et al., 2023; Liu et al., 2023), snakes (Huang et al., 2022), and many insects (Jelley & Barden, 2021; Land, 1989; Talarico et al., 2007; Warrant, 2001), with visually complex environments often demanding larger eyes, increased acuity, and often regional specialisation for increased resolution or sensitivity to specific areas of the visual scene. We might therefore expect spiders moving through highly structured habitats, such as leaf litter or foliage, to have larger eyes than those living in two-dimensional webs or open habitats. Although we only examined eye diameter, the modular nature of the spider visual system means that we are also able to capture some measure of regional specialisation in the calculated variance between eye pairs, which could plausibly be greater in more visually complex habitats.

Finally, the use of eye diameter alone does not reflect every aspect of size and its functional implications. Axial length, or depth of the eye, is also critical to achievable acuity as this determines the focal length and the angular spacing of photoreceptors. Other studies have found slightly different allometric relationships between axial length and corneal diameter (Hall & Ross, 2007; Ross & Kirk, 2007), and to corresponding ecological factors. In birds, for example, axial lengths were found to be greater in diurnal species that required greater acuity, while corneal diameters were greater in dark-living species that are likely to prioritise sensitivity (Hall & Ross, 2007; Ross & Kirk, 2007).

### Selection on eye size

As well as identifying variation in the allometric relationships of different eye pairs, and allometric shifts within homologous eye pairs across spider evolution, we also find evidence that these relationships are subject to different selective dynamics. Stabilising selection is evinced by reduced evolutionary rates of relative eye diameter (Figure 6). This appears to act on eye pairs that perform specialised visual functions, such as the AMEs and ALEs of salticids, whose tight static allometric relationships may further reflect constraint (Figure 2). Stabilising selection may also affect eye pairs that are reduced in size but probably still functional, such as the AMEs of pholcids, thomisids, and philodromids. Conversely, increased evolutionary rates may signal two possibilities. First, they can result from the relaxation of selection, which may also be apparent from looser static allometric relationships, as seen in the PMEs of salticids. Alternatively, increased rates of evolution may indicate directional selection within the clade, as discussed in the PMEs of *Deinopis* and *Menneus*. Although increased evolutionary rates were detected in the relative size of the PMEs in lycosids and oxyopids, which are critical to visual hunting, this was driven by changes to carapace width rather than eye diameter.

### Conclusions

Eye size has a wide range of implications for visual function and ecology. We find that, like many other animal groups, eyes tend to scale hypoallometrically in spiders, and that eyes are generally larger when they contribute to foraging, in both diurnal and nocturnal species. However, the modular nature of the spider visual system provides additional degrees of freedom regarding the investment of space and energy. This is apparent in the strong correlations between ranked eye pairs, and variance between eye pairs, and visual hunting, as well as variable relationships between different homologous eye pairs and ecological factors.

There are several attractive routes for future work. First, although only female individuals were included in this study, sexual dimorphism is a common and important evolutionary phenomenon in spiders. Second, while we have explored the static and phylogenetic allometry of eyes in a large number of species and families, comparative work on ontogenetic allometry is still lacking. It is striking that we detected static isometry in the majority of salticid species and eye pairs investigated, in contrast to the negative ontogenetic allometry apparent in *Phidippus audax* (Goté et al., 2019), but allometric shifts between ontogenetic life stages have also been reported in other taxa (Shrimpton et al., 2021). Finally, the inclusion of additional measurements reflecting visual function would be highly informative. As well as axial length, discussed above, volumetric assessments of the visual neuropils and higher processing centres such as the arcuate body could provide rich additional insight to the relative importance of, and investment in, the modular visual system of spiders. Work by Long et al. (2021) demonstrates that the size and number of visual neuropils is highest in families that use their vision in active hunting, in line with our findings that these species invest more in their eyes, but volumetric data for the different brain regions are available for relatively few species.

## Acknowledgements

This work was funded by the Oxford University Museum of Natural History summer studentship programme, the Deutsche Forschungsgemeinschaft Emmy Noether programme (SU 1336/1-1), and Harvard University. The authors warmly thank Zoë Simmons and Ricardo Pérez-de la Fuente (OUMNH) for access to specimens and equipment, and Jonas Wolff (U. Greifswald) for helpful discussions and the use of data from Wolff et al. 2022.

## Author contributions

The project was conceived by LSR and CDP and developed by LSR and KLC. Data collection was performed by KLC. Data analysis and visualisation was conducted by KLC and LSR, with advice from CDP. Data on diurnal and nocturnal activity patterns were collated by AG. KLC and LSR wrote the manuscript and prepared figures. All authors contributed to editing and approved the final manuscript.

## Data availability

All measurements generated for static allometry will be made available as part of the World Spider Traits database. Analyses and datasets are available in the supplementary materials.

