## Supplementary material for "Complex allometric relationships and ecological factors shape the development and evolution of eye size in the modular visual system of spiders": ancestral_state_analysis.Rmd

````
---
title: "Anc_state_2"
output: html_document
date: "2023-06-10"
editor_options: 
  chunk_output_type: console
---

```{r setup, include=FALSE}
knitr::opts_chunk$set(echo = TRUE)
```

**Setup**

```{r, echo = FALSE}
#Prepare analysis
library(tidyr)
library(caper)
library(geiger)
library(phytools)
library(ape)
library(geiger)
library(phytools)
library(nlme)
library(evomap)
library(phylolm)
library(adephylo)
library(phangorn)
library(geomorph)

orange<-rgb(238,99,0,alpha=0.8*255,maxColorValue=255)
red<-rgb(238,99,99,alpha=0.8*255,maxColorValue=255)
blue<-rgb(100,149,237,alpha=0.8*255,maxColorValue=255)
green<-rgb(162,205,90,alpha=0.8*255,maxColorValue=255)

WolffDataRedDupsGone<- read.delim("WolffDataNAs-Dec23.txt") 

#change characters to chars
class(WolffDataRedDupsGone$terminal) = "character"
class(WolffDataRedDupsGone$builder) = "character"
class(WolffDataRedDupsGone$orbweb) = "character"
class(WolffDataRedDupsGone$hanging) = "character"
class(WolffDataRedDupsGone$guild) = "character"

tree<-read.tree("WolffRed22Sept.nwk")

datadiff<-setdiff(WolffDataRedDupsGone$terminal, tree$tip.label)

#and remove these from the dataset
data<-WolffDataRedDupsGone[!WolffDataRedDupsGone$terminal %in% datadiff,]
#remove these from the tree
remove_tips <- c(datadiff)
tree <- drop.tip(tree, remove_tips)
data_all <- data
tree_all <- tree


```

**Plot the tree highlighting different clades**

```{r}
 pdf("Plots/wolff_tree.pdf",height=65,width=25)
#set up the different edges
edge_col<-rep("black",length(tree$edge.length))
edge_col[getEdges(tree,findMRCA(tree,c("Theraphosidae_Augacephalus_junodi", "Atypidae_Atypus_affinis")))]<-"red"
edge_col[getEdges(tree,findMRCA(tree,c("Pholcidae_Pholcus_kakum", "Trogloraptoridae_Trogloraptor_marchingtoni")))]<-"orange"
edge_col[getEdges(tree,findMRCA(tree,c("Corinnidae_Serendib_volans", "Eresidae_Stegodyphus_lineatus")))]<-"lightblue"
edge_col[getEdges(tree,findMRCA(tree,c("Linyphiidae_Microlinyphia_sp_GH39", "Theridiidae_Crustulina_sticta")))]<-"blue"
edge_col[getEdges(tree,findMRCA(tree,c("Corinnidae_Serendib_volans", "Uloboridae_Uloborus_glomosus")))]<-"green"
edge_col[getEdges(tree,findMRCA(tree,c("Desidae_Paramatachia_sp_BinnaBurra", "Amaurobiidae_Callobius_bennetti")))]<-"purple"
edge_col[getEdges(tree,findMRCA(tree,c("Thomisidae_Runcinia_albostriata", "Zoropsidae_Phanotea_sathegyna")))]<-"pink"
plot(tree,edge.col=edge_col)
dev.off()

#plot the different clades
 pdf("Plots/Mygalomorphae_tree.pdf",height=65,width=25)
edge_col<-rep("black",length(tree$edge.length))
edge_col[getEdges(tree,findMRCA(tree,c("Theraphosidae_Augacephalus_junodi", "Atypidae_Atypus_affinis")))]<-"red"
plot(tree,edge.col=edge_col)
dev.off()
pdf("Plots/Synspermiata_tree.pdf",height=65,width=25)
edge_col<-rep("black",length(tree$edge.length))
edge_col[getEdges(tree,findMRCA(tree,c("Pholcidae_Pholcus_kakum", "Trogloraptoridae_Trogloraptor_marchingtoni")))]<-"orange"
plot(tree,edge.col=edge_col)
dev.off()

pdf("Plots/Entelegynes_tree.pdf",height=65,width=25)
edge_col<-rep("black",length(tree$edge.length))
edge_col[getEdges(tree,findMRCA(tree,c("Corinnidae_Serendib_volans", "Eresidae_Stegodyphus_lineatus")))]<-"lightblue"
plot(tree,edge.col=edge_col)
dev.off()

pdf("Plots/Aranoidea_tree.pdf",height=65,width=25)
edge_col<-rep("black",length(tree$edge.length))
edge_col[getEdges(tree,findMRCA(tree,c("Linyphiidae_Microlinyphia_sp_GH39", "Theridiidae_Crustulina_sticta")))]<-"blue"
plot(tree,edge.col=edge_col)
dev.off()

pdf("Plots/RTA_tree.pdf",height=65,width=25)
edge_col<-rep("black",length(tree$edge.length))
edge_col[getEdges(tree,findMRCA(tree,c("Corinnidae_Serendib_volans", "Uloboridae_Uloborus_glomosus")))]<-"green"
plot(tree,edge.col=edge_col)
dev.off()

pdf("Plots/Marronoid_tree.pdf",height=65,width=25)
edge_col<-rep("black",length(tree$edge.length))
edge_col[getEdges(tree,findMRCA(tree,c("Desidae_Paramatachia_sp_BinnaBurra", "Amaurobiidae_Callobius_bennetti")))]<-"purple"
plot(tree,edge.col=edge_col)
dev.off()

pdf("Plots/oval_tree.pdf",height=65,width=25)
edge_col<-rep("black",length(tree$edge.length))
edge_col[getEdges(tree,findMRCA(tree,c("Thomisidae_Runcinia_albostriata", "Zoropsidae_Phanotea_sathegyna")))]<-"pink"
plot(tree,edge.col=edge_col)
dev.off()
```

## Plot each eye pair (abs + rel)

**logAME**

```{r}
#drop the rows in the data with no AMEs
data = data_all%>% drop_na(AME)
xx <- c(data$terminal)
common_names <- intersect(tree$tip.label, xx)
tree <- drop.tip(tree_all, tree$tip.label[!tree_all$tip.label %in% common_names])
rownames(data) <- data$terminal
Names <- rownames(data)

#AncEst - rate estmiation and run into the bayesian chain 
dat <- log(data$AMEmm)
names(dat) <- Names
	mvBMresults<-mvBM(dat,tree,ace(dat,tree,method="REML")$sigma2[1])
	# calculate rescaled branch lengths using mvBM
		tree_mvBM<-tree
		tree_mvBM$edge.length<-mvBMresults$rBL #create new tree with rescaled branch lengths
		plot(tree_mvBM,cex=0.2)

	iterations<-1000000
    model_mvBM<-anc.Bayes(tree_mvBM,dat,ngen=iterations)  
    # 'anc.Bayes' requires the 'phytools' package; this may take a few minutes
    MCMC_mvBM_sigma2<-model_mvBM$mcmc[(iterations*0.002):nrow(model_mvBM$mcmc),2] 
    MCMC_mvBM_anc<-colMeans(model_mvBM$mcmc[(iterations*0.002):nrow(model_mvBM$mcmc),])[3:(3+tree$Nnode-1)]   #These are the ancestral estimates
    MCMC_mvBM_logLik<-model_mvBM$mcmc[(iterations*0.002):nrow(model_mvBM$mcmc),tree$Nnode+3]

    hist(MCMC_mvBM_logLik)  
   # Plot Scattergram
	anc<-MCMC_mvBM_anc
            obj<-contMap(tree,dat,anc.states=anc,plot=FALSE,outline=FALSE,show.tip.label=FALSE,label.offset=50)
            n<-length(obj$cols)
            obj$cols[1:n]<-colorRampPalette(c("blue","blue","green","yellow","red"),space="Lab")(n)
 pdf("Plots/logame_tree.pdf",height=65,width=25)
            plot(obj,type="phylogram",legend=0.7*max(nodeHeights(tree)),fsize=c(0.7,0.9),label.offset=3,outline=FALSE,show.tip.label=FALSE,label.offset=50)
dev.off()
```

**relAME**

```{r}
#Run lambda model
data = data_all%>% drop_na(rAME)
xx <- c(data$terminal)
common_names <- intersect(tree_all$tip.label, xx)
tree <- drop.tip(tree_all, tree_all$tip.label[!tree_all$tip.label %in% common_names])
rownames(data) <- data$terminal
Names <- rownames(data)
#mvBM
Y<-"rAME"; X<-"CWmm"
data<-data[,c(which(colnames(data)==Y),which(colnames(data)==X)),drop=F]
colnames(data)<-c("Dependent","Independent") 
#log the data
data$Dependent <- log(data$Dependent)
data$Independent <- log(data$Independent)

pGLS<-phylolm(Dependent~Independent,data,phy=tree,model="lambda")
dat<-pGLS$residuals
mvBMresults<-mvBM(dat,tree,ace(dat,tree,method="REML")$sigma2[1]) # calculate rescaled branch lengths using mvBM
tree_mvBM<-tree
tree_mvBM$edge.length<-mvBMresults$rBL # create new tree with rescaled branch lengths
anc_mvBM<-ace(dat,tree_mvBM,method="REML")$ace

trait_all<-c(as.matrix(dat)[,1],anc_mvBM)

pdf("Plots/relAME_tree.pdf",height=65,width=25)
            plot(obj,type="phylogram",legend=0.7*max(nodeHeights(tree)),fsize=c(0.7,0.9),label.offset=3,outline=FALSE,show.tip.label=FALSE,label.offset=50)
dev.off()
```

**logALE**

```{r}
data = data_all%>% drop_na(ALE)
xx <- c(data$terminal)
common_names <- intersect(tree$tip.label, xx)
tree <- drop.tip(tree_all, tree$tip.label[!tree_all$tip.label %in% common_names])
rownames(data) <- data$terminal
Names <- rownames(data)
dat <- log(data$ALEmm)
names(dat) <- Names
	mvBMresults<-mvBM(dat,tree,ace(dat,tree,method="REML")$sigma2[1])
	# calculate rescaled branch lengths using mvBM
		tree_mvBM<-tree
		tree_mvBM$edge.length<-mvBMresults$rBL #create new tree with rescaled branch lengths
		plot(tree_mvBM,cex=0.2)

	iterations<-1000000
    model_mvBM<-anc.Bayes(tree_mvBM,dat,ngen=iterations)  
    # 'anc.Bayes' requires the 'phytools' package; this may take a few minutes
    MCMC_mvBM_sigma2<-model_mvBM$mcmc[(iterations*0.002):nrow(model_mvBM$mcmc),2] 
    MCMC_mvBM_anc<-colMeans(model_mvBM$mcmc[(iterations*0.002):nrow(model_mvBM$mcmc),])[3:(3+tree$Nnode-1)]   #These are the ancestral estimates
    MCMC_mvBM_logLik<-model_mvBM$mcmc[(iterations*0.002):nrow(model_mvBM$mcmc),tree$Nnode+3]

    hist(MCMC_mvBM_logLik)  
   # Plot Scattergram
	anc<-MCMC_mvBM_anc
            obj<-contMap(tree,dat,anc.states=anc,plot=FALSE,outline=FALSE,show.tip.label=FALSE,label.offset=50)
            n<-length(obj$cols)
            obj$cols[1:n]<-colorRampPalette(c("blue","blue","green","yellow","red"),space="Lab")(n)
 pdf("Plots/scattergram_ale_tree.pdf",height=65,width=25)
            plot(obj,type="phylogram",legend=0.7*max(nodeHeights(tree)),fsize=c(0.7,0.9),label.offset=3,outline=FALSE,show.tip.label=FALSE,label.offset=50)
dev.off()
```

**relALE**

```{r}
#Run lambda model
data = data_all%>% drop_na(rALE)
xx <- c(data$terminal)
common_names <- intersect(tree_all$tip.label, xx)
tree <- drop.tip(tree_all, tree_all$tip.label[!tree_all$tip.label %in% common_names])
rownames(data) <- data$terminal
Names <- rownames(data)
#mvBM
Y<-"rALE"; X<-"CWmm"
data<-data[,c(which(colnames(data)==Y),which(colnames(data)==X)),drop=F]
colnames(data)<-c("Dependent","Independent") 
#log the data
data$Dependent <- log(data$Dependent)
data$Independent <- log(data$Independent)

pGLS<-phylolm(Dependent~Independent,data,phy=tree,model="lambda")
dat<-pGLS$residuals
mvBMresults<-mvBM(dat,tree,ace(dat,tree,method="REML")$sigma2[1]) # calculate rescaled branch lengths using mvBM
tree_mvBM<-tree
tree_mvBM$edge.length<-mvBMresults$rBL # create new tree with rescaled branch lengths
anc_mvBM<-ace(dat,tree_mvBM,method="REML")$ace

trait_all<-c(as.matrix(dat)[,1],anc_mvBM)

pdf("Plots/relALE_tree.pdf",height=65,width=25)
            plot(obj,type="phylogram",legend=0.7*max(nodeHeights(tree)),fsize=c(0.7,0.9),label.offset=3,outline=FALSE,show.tip.label=FALSE,label.offset=50)
dev.off()
```

**logPLE**

```{r}
data = data_all%>% drop_na(PLE)
xx <- c(data$terminal)
common_names <- intersect(tree$tip.label, xx)
tree <- drop.tip(tree_all, tree$tip.label[!tree_all$tip.label %in% common_names])
rownames(data) <- data$terminal
Names <- rownames(data)
dat <- log(data$PLEmm)
names(dat) <- Names
mvBMresults<-mvBM(dat,tree,ace(dat,tree,method="REML")$sigma2[1])
# calculate rescaled branch lengths using mvBM
tree_mvBM<-tree
tree_mvBM$edge.length<-mvBMresults$rBL #create new tree with rescaled branch lengths
plot(tree_mvBM,cex=0.2)

iterations<-1000000
model_mvBM<-anc.Bayes(tree_mvBM,dat,ngen=iterations)  
# 'anc.Bayes' requires the 'phytools' package; this may take a few minutes
MCMC_mvBM_sigma2<-model_mvBM$mcmc[(iterations*0.002):nrow(model_mvBM$mcmc),2] 
MCMC_mvBM_anc<-colMeans(model_mvBM$mcmc[(iterations*0.002):nrow(model_mvBM$mcmc),])[3:(3+tree$Nnode-1)]   #These are the ancestral estimates
MCMC_mvBM_logLik<-model_mvBM$mcmc[(iterations*0.002):nrow(model_mvBM$mcmc),tree$Nnode+3]

hist(MCMC_mvBM_logLik)  
# Plot Scattergram
anc<-MCMC_mvBM_anc
obj<-contMap(tree,dat,anc.states=anc,plot=FALSE,outline=FALSE,show.tip.label=FALSE,label.offset=50)
n<-length(obj$cols)
obj$cols[1:n]<-colorRampPalette(c("blue","blue","green","yellow","red"),space="Lab")(n)
pdf("Plots/scattergram_ple_tree.pdf",height=65,width=25)
plot(obj,type="phylogram",legend=0.7*max(nodeHeights(tree)),fsize=c(0.7,0.9),label.offset=3,outline=FALSE,show.tip.label=FALSE,label.offset=50)
dev.off()
trait_all<-c(dat,anc)
```

**relPLE**

```{r}
#Run lambda model
data = data_all%>% drop_na(rPLE)
xx <- c(data$terminal)
common_names <- intersect(tree_all$tip.label, xx)
tree <- drop.tip(tree_all, tree_all$tip.label[!tree_all$tip.label %in% common_names])
rownames(data) <- data$terminal
Names <- rownames(data)
#mvBM
Y<-"rPLE"; X<-"CWmm"
data<-data[,c(which(colnames(data)==Y),which(colnames(data)==X)),drop=F]
colnames(data)<-c("Dependent","Independent") 
#log the data
data$Dependent <- log(data$Dependent)
data$Independent <- log(data$Independent)

pGLS<-phylolm(Dependent~Independent,data,phy=tree,model="lambda")
dat<-pGLS$residuals
mvBMresults<-mvBM(dat,tree,ace(dat,tree,method="REML")$sigma2[1]) # calculate rescaled branch lengths using mvBM
tree_mvBM<-tree
tree_mvBM$edge.length<-mvBMresults$rBL # create new tree with rescaled branch lengths
anc_mvBM<-ace(dat,tree_mvBM,method="REML")$ace

trait_all<-c(as.matrix(dat)[,1],anc_mvBM)

pdf("Plots/relPLE_tree.pdf",height=65,width=25)
            plot(obj,type="phylogram",legend=0.7*max(nodeHeights(tree)),fsize=c(0.7,0.9),label.offset=3,outline=FALSE,show.tip.label=FALSE,label.offset=50)
dev.off()
      #col=cols,
```

**logPME**

```{r}
data = data_all%>% drop_na(PME)
xx <- c(data$terminal)
common_names <- intersect(tree$tip.label, xx)
tree <- drop.tip(tree_all, tree$tip.label[!tree_all$tip.label %in% common_names])
rownames(data) <- data$terminal
Names <- rownames(data)
dat <- log(data$PMEmm)
names(dat) <- Names
mvBMresults<-mvBM(dat,tree,ace(dat,tree,method="REML")$sigma2[1])
# calculate rescaled branch lengths using mvBM
tree_mvBM<-tree
tree_mvBM$edge.length<-mvBMresults$rBL #create new tree with rescaled branch lengths
plot(tree_mvBM,cex=0.2)

iterations<-1000000
model_mvBM<-anc.Bayes(tree_mvBM,dat,ngen=iterations)  
# 'anc.Bayes' requires the 'phytools' package; this may take a few minutes
MCMC_mvBM_sigma2<-model_mvBM$mcmc[(iterations*0.002):nrow(model_mvBM$mcmc),2] 
MCMC_mvBM_anc<-colMeans(model_mvBM$mcmc[(iterations*0.002):nrow(model_mvBM$mcmc),])[3:(3+tree$Nnode-1)]   #These are the ancestral estimates
MCMC_mvBM_logLik<-model_mvBM$mcmc[(iterations*0.002):nrow(model_mvBM$mcmc),tree$Nnode+3]

hist(MCMC_mvBM_logLik)  
# Plot Scattergram
anc<-MCMC_mvBM_anc
obj<-contMap(tree,dat,anc.states=anc,plot=FALSE,outline=FALSE,show.tip.label=FALSE,label.offset=50)
n<-length(obj$cols)
obj$cols[1:n]<-colorRampPalette(c("blue","blue","green","yellow","red"),space="Lab")(n)
pdf("Plots/scattergram_pme_tree.pdf",height=65,width=25)
plot(obj,type="phylogram",legend=0.7*max(nodeHeights(tree)),fsize=c(0.7,0.9),label.offset=3,outline=FALSE,show.tip.label=FALSE,label.offset=50)
dev.off()
```

**relPME**

```{r}
#Run lambda model
data = data_all%>% drop_na(rPME)
xx <- c(data$terminal)
common_names <- intersect(tree_all$tip.label, xx)
tree <- drop.tip(tree_all, tree_all$tip.label[!tree_all$tip.label %in% common_names])
rownames(data) <- data$terminal
Names <- rownames(data)
#mvBM
Y<-"rPME"; X<-"CWmm"
data<-data[,c(which(colnames(data)==Y),which(colnames(data)==X)),drop=F]
colnames(data)<-c("Dependent","Independent") 
#log the data
data$Dependent <- log(data$Dependent)
data$Independent <- log(data$Independent)

pGLS<-phylolm(Dependent~Independent,data,phy=tree,model="lambda")
dat<-pGLS$residuals
mvBMresults<-mvBM(dat,tree,ace(dat,tree,method="REML")$sigma2[1]) # calculate rescaled branch lengths using mvBM
tree_mvBM<-tree
tree_mvBM$edge.length<-mvBMresults$rBL # create new tree with rescaled branch lengths
anc_mvBM<-ace(dat,tree_mvBM,method="REML")$ace

trait_all<-c(as.matrix(dat)[,1],anc_mvBM)

pdf("Plots/relPME_tree.pdf",height=65,width=25)
            plot(obj,type="phylogram",legend=0.7*max(nodeHeights(tree)),fsize=c(0.7,0.9),label.offset=3,outline=FALSE,show.tip.label=FALSE,label.offset=50)
dev.off()
```

**logCW**

```{r}
#drop the rows in the data with no CWmm
data = data_all%>% drop_na(CWmm)
xx <- c(data$terminal)
common_names <- intersect(tree_all$tip.label, xx)
tree <- drop.tip(tree_all, tree_all$tip.label[!tree_all$tip.label %in% common_names])
rownames(data) <- data$terminal
Names <- rownames(data)

#AncEst - rate estmiation and run into the bayesian chain 
dat <- log(data$CWmm)
names(dat) <- Names
	mvBMresults<-mvBM(dat,tree,ace(dat,tree,method="REML")$sigma2[1])
	# calculate rescaled branch lengths using mvBM
		tree_mvBM<-tree
		tree_mvBM$edge.length<-mvBMresults$rBL #create new tree with rescaled branch lengths
		plot(tree_mvBM,cex=0.2)

	iterations<-1000000
    model_mvBM<-anc.Bayes(tree_mvBM,dat,ngen=iterations)  
    # 'anc.Bayes' requires the 'phytools' package; this may take a few minutes
    MCMC_mvBM_sigma2<-model_mvBM$mcmc[(iterations*0.002):nrow(model_mvBM$mcmc),2] 
    MCMC_mvBM_anc<-colMeans(model_mvBM$mcmc[(iterations*0.002):nrow(model_mvBM$mcmc),])[3:(3+tree$Nnode-1)]   #These are the ancestral estimates
    MCMC_mvBM_logLik<-model_mvBM$mcmc[(iterations*0.002):nrow(model_mvBM$mcmc),tree$Nnode+3]

    hist(MCMC_mvBM_logLik)  
   # Plot Scattergram
	anc<-MCMC_mvBM_anc
            obj<-contMap(tree,dat,anc.states=anc,plot=FALSE,outline=FALSE,show.tip.label=FALSE,label.offset=50)
            n<-length(obj$cols)
            obj$cols[1:n]<-colorRampPalette(c("blue","blue","green","yellow","red"),space="Lab")(n)
 pdf("Plots/scattergram_CW_tree.pdf",height=65,width=25)
            plot(obj,type="phylogram",legend=0.7*max(nodeHeights(tree)),fsize=c(0.7,0.9),label.offset=3,outline=FALSE,show.tip.label=FALSE,label.offset=50)
dev.off()
```

**variance**

```{r}
data = data_all%>% drop_na(var)
xx <- c(data$terminal)
common_names <- intersect(tree_all$tip.label, xx)
tree <- drop.tip(tree_all, tree_all$tip.label[!tree_all$tip.label %in% common_names])
rownames(data) <- data$terminal
Names <- rownames(data)
dat <-data$var
names(dat) <- Names
dat <- dat[is.finite(dat)]
common_names <- intersect(tree$tip.label, names(dat))
tree <- drop.tip(tree, tree$tip.label[!tree$tip.label %in% common_names])
#AncEst - rate estmiation and run into the bayesian chain 
mvBMresults<-mvBM(dat,tree,ace(dat,tree,method="REML")$sigma2[1])
# calculate rescaled branch lengths using mvBM
tree_mvBM<-tree
tree_mvBM$edge.length<-mvBMresults$rBL #create new tree with rescaled branch lengths
plot(tree_mvBM,cex=0.2)

iterations<-1000000
model_mvBM<-anc.Bayes(tree_mvBM,dat,ngen=iterations)  
# 'anc.Bayes' requires the 'phytools' package; this may take a few minutes
MCMC_mvBM_sigma2<-model_mvBM$mcmc[(iterations*0.002):nrow(model_mvBM$mcmc),2] 
MCMC_mvBM_anc<-colMeans(model_mvBM$mcmc[(iterations*0.002):nrow(model_mvBM$mcmc),])[3:(3+tree$Nnode-1)]   #These are the ancestral estimates
MCMC_mvBM_logLik<-model_mvBM$mcmc[(iterations*0.002):nrow(model_mvBM$mcmc),tree$Nnode+3]

hist(MCMC_mvBM_logLik)  
# Plot Scattergram
pdf("Plots/variance_tree.pdf",height=65,width=25)
anc<-MCMC_mvBM_anc
obj<-contMap(tree,dat,anc.states=anc,plot=FALSE,outline=FALSE,show.tip.label=FALSE,label.offset=50)
n<-length(obj$cols)
obj$cols[1:n]<-colorRampPalette(c("blue","blue","green","yellow","red"),space="Lab")(n)
plot(obj,type="phylogram",legend=0.7*max(nodeHeights(tree)),fsize=c(0.7,0.9),label.offset=3,outline=FALSE,show.tip.label=FALSE,label.offset=50)
dev.off()
```

## Calculating evolutionary rates 

Allocate ancestral clades

```{r}
All<-c(1:length(tree$tip.label))
Mygalomorphae  <-getTips(tree,findMRCA(tree,c("Theraphosidae_Augacephalus_junodi", "Atypidae_Atypus_affinis")))
Synspermiata <-getTips(tree,findMRCA(tree,c("Pholcidae_Pholcus_kakum", "Caponiidae_Nops_sp_Jaragua")))
Entelegynes<-getTips(tree,findMRCA(tree,c("Corinnidae_Serendib_volans", "Eresidae_Stegodyphus_lineatus")))
Aranoidea<-getTips(tree,findMRCA(tree,c("Linyphiidae_Microlinyphia_sp_GH39", "Theridiidae_Crustulina_sticta")))
RTA <-getTips(tree,findMRCA(tree,c("Corinnidae_Serendib_volans", "Uloboridae_Uloborus_glomosus")))
Marronoid<-getTips(tree,findMRCA(tree,c("Desidae_Paramatachia_sp_BinnaBurra", "Amaurobiidae_Callobius_bennetti")))
Oval<-getTips(tree,findMRCA(tree,c("Thomisidae_Runcinia_albostriata", "Zoropsidae_Phanotea_sathegyna")))
```

#### **logAME**

```{r}
#allocate groups as noted from bayou:
pholcid <-getTips(tree,findMRCA(tree,c("Pholcidae_Pholcus_kakum", "Pholcidae_Artema_atlanta")))
anapid<-getTips(tree,findMRCA(tree,c("Anapidae_Acrobleps_hygrophilus", "Anapidae_Conculus_lyugadinus")))
ctenid_psechrid<-getTips(tree,findMRCA(tree,c("Psechridae_Fecenia_protensa", "Ctenidae_Ctenus_crulsi")))
thomisid_oxyopid<-getTips(tree,findMRCA(tree,c("Oxyopidae_Peucetia_viridans", "Thomisidae_Runcinia_albostriata")))
salticid<-getTips(tree,findMRCA(tree,c("Salticidae_Lyssomanes_viridis", "Salticidae_Salticus_scenicus")))
philodromid<-getTips(tree,findMRCA(tree,c("Philodromidae_Thanatus_formicinus", "Philodromidae_Philodromus_aureolus")))
#drop the rows in the data with no AMEs
data = data_all%>% drop_na(AME)
xx <- c(data$terminal)
common_names <- intersect(tree$tip.label, xx)
tree <- drop.tip(tree_all, tree$tip.label[!tree_all$tip.label %in% common_names])
rownames(data) <- data$terminal
Names <- rownames(data)
#AncEst - rate estmiation and run into the bayesian chain 
dat <- log(data$AMEmm)
names(dat) <- Names
	mvBMresults<-mvBM(dat,tree,ace(dat,tree,method="REML")$sigma2[1])
	# calculate rescaled branch lengths using mvBM
		tree_mvBM<-tree
		tree_mvBM$edge.length<-mvBMresults$rBL #create new tree with rescaled branch lengths
		plot(tree_mvBM,cex=0.2)
#setting seed
set.seed(0)
	iterations<-1000000
    model_mvBM<-anc.Bayes(tree_mvBM,dat,ngen=iterations)  
    # 'anc.Bayes' requires the 'phytools' package; this may take a few minutes
    MCMC_mvBM_sigma2<-model_mvBM$mcmc[(iterations*0.002):nrow(model_mvBM$mcmc),2] 
    MCMC_mvBM_anc<-colMeans(model_mvBM$mcmc[(iterations*0.002):nrow(model_mvBM$mcmc),])[3:(3+tree$Nnode-1)]   #These are the ancestral estimates
    MCMC_mvBM_logLik<-model_mvBM$mcmc[(iterations*0.002):nrow(model_mvBM$mcmc),tree$Nnode+3]

    #save pdf of the rates calculated for each shift compared to the larger clade as found by Bayou
pdf("AME_ancestral_tree.pdf",height=32,width=45)
par(mfrow = c(2, 3))
Branches_synsperimata<-tail(getEdges(tree,findMRCA(tree,setdiff(Synspermiata, pholcid))),n=1)
Branches_pholcid<-tail(getEdges(tree,findMRCA(tree,pholcid)),n=1)
    mvBM.getRate(tree,tree_mvBM,branches=Branches_synsperimata,sigma2Distr = MCMC_mvBM_sigma2)
    mvBM.getRate(tree,tree_mvBM,branches=Branches_pholcid,sigma2Distr = MCMC_mvBM_sigma2)
    mvBM.plotRate(tree,tree_mvBM,branches=Branches_synsperimata,sigma2Distr = MCMC_mvBM_sigma2,col="lightgrey",adjust=6,main="", xlim=c(0.001, 0.0041), ylim = c(0, 3100))
    par(new=TRUE)
    mvBM.plotRate(tree,tree_mvBM,branches=Branches_pholcid,sigma2Distr = MCMC_mvBM_sigma2,col="salmon",adjust=6,main="",xlim=c(0.001, 0.0041), ylim = c(0, 3100))
legend(x = "topleft", legend = c("Synspermiata", "Pholcidae"), col = c("lightgrey", "salmon"), lty = 10, lwd = 10, bty = "n")

Branches_thomisid_oxyopid<-tail(getEdges(tree,findMRCA(tree,setdiff(Oval, thomisid_oxyopid))),n=1)
Branches_oval<-tail(getEdges(tree,findMRCA(tree,thomisid_oxyopid)),n=1)
mvBM.getRate(tree,tree_mvBM,branches=Branches_oval,sigma2Distr = MCMC_mvBM_sigma2)
    mvBM.getRate(tree,tree_mvBM,branches=Branches_thomisid_oxyopid,sigma2Distr = MCMC_mvBM_sigma2)
    
    mvBM.plotRate(tree,tree_mvBM,branches=Branches_oval,sigma2Distr = MCMC_mvBM_sigma2,col="lightgrey",adjust=6,main="",xlim=c(0.001, 0.0035), ylim = c(0, 3000))
    par(new=TRUE)
    mvBM.plotRate(tree,tree_mvBM,branches=Branches_thomisid_oxyopid,sigma2Distr = MCMC_mvBM_sigma2,col="cyan3",adjust=6,main="",xlim=c(0.001, 0.0035), ylim = c(0, 3000))
legend(x = "topleft", legend = c("Oval", "Thomisidae & Oxyopidae"), col = c("lightgrey", "cyan3"), lty = 10, lwd = 10, bty = "n", cex = 0.5)

Branches_anapid<-tail(getEdges(tree,findMRCA(tree,anapid)),n=1)
Branches_aranoidea<-tail(getEdges(tree,findMRCA(tree,setdiff(Aranoidea, anapid))),n=1)
mvBM.getRate(tree,tree_mvBM,branches=Branches_aranoidea,sigma2Distr = MCMC_mvBM_sigma2)
    mvBM.getRate(tree,tree_mvBM,branches=Branches_anapid,sigma2Distr = MCMC_mvBM_sigma2)

    mvBM.plotRate(tree,tree_mvBM,branches=Branches_aranoidea,sigma2Distr = MCMC_mvBM_sigma2,col="lightgrey",adjust=6,main="", xlim=c(0.0012, 0.0030), ylim = c(0, 3000))
    par(new=TRUE)
    mvBM.plotRate(tree,tree_mvBM,branches=Branches_anapid,sigma2Distr = MCMC_mvBM_sigma2,col="darkolivegreen4",adjust=6,main="",xlim=c(0.0012, 0.0030), ylim = c(0, 3000))
      legend(x = "topright", legend = c("Aranoidea", "Anapidae"), col = c("lightgrey", "darkolivegreen4"), lty = 10, lwd = 10,  bty = "n")

Branches_salticid<-tail(getEdges(tree,findMRCA(tree,salticid)),n=1)
Branches_RTA<-tail(getEdges(tree,findMRCA(tree,setdiff(RTA, c(salticid)))),n=1)
mvBM.getRate(tree,tree_mvBM,branches=Branches_salticid,sigma2Distr = MCMC_mvBM_sigma2)
    mvBM.getRate(tree,tree_mvBM,branches=Branches_RTA,sigma2Distr = MCMC_mvBM_sigma2)

    mvBM.plotRate(tree,tree_mvBM,branches=Branches_RTA,sigma2Distr = MCMC_mvBM_sigma2,col="lightgrey",adjust=6,main="", xlim=c(0.0018, 0.006), ylim = c(0, 2000))
    par(new=TRUE)
    mvBM.plotRate(tree,tree_mvBM,branches=Branches_salticid,sigma2Distr = MCMC_mvBM_sigma2,col="magenta",adjust=6,main="",xlim=c(0.0018, 0.006), ylim = c(0, 2000))
legend(x = "topright", legend = c("RTA", "Salticidae"), col = c("lightgrey", "magenta"), lty = 10, lwd = 10, bty = "n")

ctenid_psechrid<-getTips(tree,findMRCA(tree,c("Psechridae_Fecenia_protensa", "Ctenidae_Ctenus_crulsi")))
philodromid<-getTips(tree,findMRCA(tree,c("Philodromidae_Thanatus_formicinus", "Philodromidae_Philodromus_aureolus")))

Branches_ctenid_psechrid<-tail(getEdges(tree,findMRCA(tree,ctenid_psechrid)),n=1)
Branches_oval<-tail(getEdges(tree,findMRCA(tree,setdiff(Oval, ctenid_psechrid))),n=1)
mvBM.getRate(tree,tree_mvBM,branches=Branches_oval,sigma2Distr = MCMC_mvBM_sigma2)
mvBM.getRate(tree,tree_mvBM,branches=Branches_ctenid_psechrid,sigma2Distr = MCMC_mvBM_sigma2)
mvBM.plotRate(tree,tree_mvBM,branches=Branches_oval,sigma2Distr =MCMC_mvBM_sigma2,col="lightgrey",adjust=6,main="", xlim=c(0.0011, 0.0026), ylim = c(0, 2950))
    par(new=TRUE)
    mvBM.plotRate(tree,tree_mvBM,branches=Branches_anapid,sigma2Distr = MCMC_mvBM_sigma2,col="darkolivegreen4",adjust=6,main="",xlim=c(0.0011, 0.0026), ylim = c(0, 2950))
      legend(x = "topright", legend = c("Oval", "Ctenid_psechrid"), col = c("lightgrey", "darkolivegreen4"), lty = 10, lwd = 10,  bty = "n", cex = 0.5)

Branches_philodromid<-tail(getEdges(tree,findMRCA(tree,philodromid)),n=1)
Branches_RTA<-tail(getEdges(tree,findMRCA(tree,setdiff(RTA, philodromid))),n=1)
mvBM.getRate(tree,tree_mvBM,branches=Branches_RTA,sigma2Distr = MCMC_mvBM_sigma2)
mvBM.getRate(tree,tree_mvBM,branches=Branches_philodromid,sigma2Distr = MCMC_mvBM_sigma2)
mvBM.plotRate(tree,tree_mvBM,branches=Branches_RTA,sigma2Distr =MCMC_mvBM_sigma2,col="lightgrey",adjust=6,main="",xlim=c(0.0011, 0.006), ylim = c(0, 3000))
    par(new=TRUE)
    mvBM.plotRate(tree,tree_mvBM,branches=Branches_philodromid,sigma2Distr = MCMC_mvBM_sigma2,col="darkolivegreen4",adjust=6,main="",xlim=c(0.0011, 0.006), ylim = c(0, 3000))
      legend(x = "topright", legend = c("RTA", "philodromid"), col = c("lightgrey", "darkolivegreen4"), lty = 10, lwd = 10,  bty = "n")
    dev.off()
```

Repeat for all other eye pairs

#### **logALE**

```{r}
data = data_all%>% drop_na(ALE)
xx <- c(data$terminal)
common_names <- intersect(tree_all$tip.label, xx)
tree <- drop.tip(tree_all, tree_all$tip.label[!tree_all$tip.label %in% common_names])
rownames(data) <- data$terminal
Names <- rownames(data)
dat <- log(data$ALEmm)
names(dat) <- Names
	mvBMresults<-mvBM(dat,tree,ace(dat,tree,method="REML")$sigma2[1])
	# calculate rescaled branch lengths using mvBM
		tree_mvBM<-tree
		tree_mvBM$edge.length<-mvBMresults$rBL #create new tree with rescaled branch lengths
		plot(tree_mvBM,cex=0.2)
#setting seed
set.seed(0)
	iterations<-1000000
    model_mvBM<-anc.Bayes(tree_mvBM,dat,ngen=iterations)  
    # 'anc.Bayes' requires the 'phytools' package; this may take a few minutes
    MCMC_mvBM_sigma2<-model_mvBM$mcmc[(iterations*0.002):nrow(model_mvBM$mcmc),2] 
    MCMC_mvBM_anc<-colMeans(model_mvBM$mcmc[(iterations*0.002):nrow(model_mvBM$mcmc),])[3:(3+tree$Nnode-1)]   #These are the ancestral estimates
    MCMC_mvBM_logLik<-model_mvBM$mcmc[(iterations*0.002):nrow(model_mvBM$mcmc),tree$Nnode+3]
```

**Plotting rates**

```{r}
corinnid_miturgid<-getTips(tree,findMRCA(tree,c("Corinnidae_Serendib_volans", "Miturgidae_Syspira_sp_cf_longipes")))
salticid<-getTips(tree,findMRCA(tree,c("Salticidae_Lyssomanes_viridis", "Salticidae_Salticus_scenicus")))
lycosid <- getTips(tree,findMRCA(tree,c("Lycosidae_Schizocosa_ocreata","Pisauridae_Bradystichus_crispatus")))
hersillid_oecobiid <- getTips(tree,findMRCA(tree,c("Hersiliidae_Iviraiva_argentina","Oecobiidae_Uroctea_durandi")))
araneidae <- getTips(tree,findMRCA(tree,c("Araneidae_Eriophora_ravilla", "Araneidae_Phonognatha_graeffei")))

lightgrey <- rgb(210, 210, 210,alpha=0.8*255,maxColorValue=255)

pdf("ALE_ancestral_tree.pdf",height=30,width=45)
par(mfrow = c(2, 3))

#plot 1
Branches_RTA<-tail(getEdges(tree,findMRCA(tree,setdiff(RTA, c(corinnid_miturgid)))),n=1)
Branches_corinnid_miturgid<-tail(getEdges(tree,findMRCA(tree,corinnid_miturgid)),n=1)
    mvBM.getRate(tree,tree_mvBM,branches=Branches_RTA,sigma2Distr = MCMC_mvBM_sigma2)
    mvBM.getRate(tree,tree_mvBM,branches=Branches_corinnid_miturgid,sigma2Distr = MCMC_mvBM_sigma2)
    mvBM.plotRate(tree,tree_mvBM,branches=Branches_RTA,sigma2Distr = MCMC_mvBM_sigma2,col="lightgrey",adjust=6,main="", xlim=c(0.00131, 0.0063), ylim = c(0, 13000))
    par(new=TRUE)
    mvBM.plotRate(tree,tree_mvBM,branches=Branches_corinnid_miturgid,sigma2Distr = MCMC_mvBM_sigma2,col="#09ef09",adjust=6,main="",xlim=c(0.00131, 0.0063), ylim = c(0, 13000) )
legend(x = "topleft", legend = c("RTA", "corinnid_miturgid"), col = c("lightgrey", "#09ef09"), lty = 10, lwd = 10, bty = "n")

#Plot 2
Branches_salticid<-tail(getEdges(tree,findMRCA(tree,salticid)),n=1)
Branches_RTA<-tail(getEdges(tree,findMRCA(tree,setdiff(RTA, c(salticid)))),n=1)
mvBM.getRate(tree,tree_mvBM,branches=Branches_salticid,sigma2Distr = MCMC_mvBM_sigma2)
    mvBM.getRate(tree,tree_mvBM,branches=Branches_RTA,sigma2Distr = MCMC_mvBM_sigma2)
    mvBM.plotRate(tree,tree_mvBM,branches=Branches_RTA,sigma2Distr = MCMC_mvBM_sigma2,col="lightgrey",adjust=6,main="", xlim=c(0.0013, 0.0063), ylim = c(0, 12500))
    par(new=TRUE)
    mvBM.plotRate(tree,tree_mvBM,branches=Branches_salticid,sigma2Distr = MCMC_mvBM_sigma2,col="#09ef09",adjust=6,main="",xlim=c(0.0013, 0.0063), ylim = c(0, 12500))
legend(x = "topright", legend = c("RTA", "Salticidae"), col = c("lightgrey", "#09ef09"), lty = 10, lwd = 10, bty = "n")

#Plot 3
Branches_lycosid<-tail(getEdges(tree,findMRCA(tree,lycosid)),n=1)
Branches_Oval<-tail(getEdges(tree,findMRCA(tree,setdiff(Oval, c(lycosid)))),n=1)
mvBM.getRate(tree,tree_mvBM,branches=Branches_lycosid,sigma2Distr = MCMC_mvBM_sigma2)
mvBM.getRate(tree,tree_mvBM,branches=Branches_Oval,sigma2Distr = MCMC_mvBM_sigma2)

mvBM.plotRate(tree,tree_mvBM,branches=Branches_Oval,sigma2Distr = MCMC_mvBM_sigma2,col="lightgrey",adjust=6,main="", xlim=c(0.0013, 0.00205), ylim = c(0, 12500))
    par(new=TRUE)
mvBM.plotRate(tree,tree_mvBM,branches=Branches_lycosid,sigma2Distr = MCMC_mvBM_sigma2,col="#09ef09",adjust=6,main="",xlim=c(0.0013, 0.00205), ylim = c(0, 12500))
legend(x = "topright", legend = c("Oval", "lycosid"), col = c("lightgrey", "#09ef09"), lty = 10, lwd = 10, bty = "n")

#Plot 4
Branches_hersillid_oecobiid<-tail(getEdges(tree,findMRCA(tree,hersillid_oecobiid)),n=1)
Branches_Entelegynes<-tail(getEdges(tree,findMRCA(tree,setdiff(Entelegynes, c(hersillid_oecobiid)))),n=1)
mvBM.getRate(tree,tree_mvBM,branches=Branches_hersillid_oecobiid,sigma2Distr = MCMC_mvBM_sigma2)
mvBM.getRate(tree,tree_mvBM,branches=Branches_Entelegynes,sigma2Distr = MCMC_mvBM_sigma2)

mvBM.plotRate(tree,tree_mvBM,branches=Branches_hersillid_oecobiid,sigma2Distr = MCMC_mvBM_sigma2,col="lightgrey",adjust=6,main="", xlim=c(0.00130, 0.00305), ylim = c(0, 13000))
    par(new=TRUE)
mvBM.plotRate(tree,tree_mvBM,branches=Branches_Entelegynes,sigma2Distr = MCMC_mvBM_sigma2,col="#09ef09",adjust=6,main="",xlim=c(0.00130, 0.00305), ylim = c(0, 13000))
legend(x = "topright", legend = c("Entelegynes", "hersillid_oecobiid"), col = c("lightgrey", "#09ef09"), lty = 10, lwd = 10, bty = "n")

#Plot 5
Branches_Aranoidea<-tail(getEdges(tree,findMRCA(tree,Aranoidea)),n=1)
Branches_araneidae<-tail(getEdges(tree,findMRCA(tree,setdiff(Aranoidea, c(araneidae)))),n=1)
mvBM.getRate(tree,tree_mvBM,branches=Branches_Aranoidea,sigma2Distr = MCMC_mvBM_sigma2)
mvBM.getRate(tree,tree_mvBM,branches=Branches_araneidae,sigma2Distr = MCMC_mvBM_sigma2)

mvBM.plotRate(tree,tree_mvBM,branches=Branches_Aranoidea,sigma2Distr = MCMC_mvBM_sigma2,col="lightgrey",adjust=6,main="", xlim=c(0.00164, 0.00194), ylim = c(0, 10000))
    par(new=TRUE)
mvBM.plotRate(tree,tree_mvBM,branches=Branches_araneidae,sigma2Distr = MCMC_mvBM_sigma2,col="#09ef09",adjust=6,main="", xlim=c(0.00164, 0.00194), ylim = c(0, 10000))
legend(x = "topright", legend = c("Aranoidea", "araneidae"), col = c("lightgrey", "#09ef09"), lty = 10, lwd = 10, bty = "n")
    dev.off()
```

#### **logPLE**

```{r}
data = data_all%>% drop_na(PLE)
xx <- c(data$terminal)
common_names <- intersect(tree_all$tip.label, xx)
tree <- drop.tip(tree_all, tree_all$tip.label[!tree_all$tip.label %in% common_names])
rownames(data) <- data$terminal
Names <- rownames(data)
dat <- log(data$PLEmm)
names(dat) <- Names
mvBMresults<-mvBM(dat,tree,ace(dat,tree,method="REML")$sigma2[1])
# calculate rescaled branch lengths using mvBM
tree_mvBM<-tree
tree_mvBM$edge.length<-mvBMresults$rBL #create new tree with rescaled branch lengths
plot(tree_mvBM,cex=0.2)
#setting seed
set.seed(0)
iterations<-1000000
model_mvBM<-anc.Bayes(tree_mvBM,dat,ngen=iterations)  
MCMC_mvBM_sigma2<-model_mvBM$mcmc[(iterations*0.002):nrow(model_mvBM$mcmc),2] 
MCMC_mvBM_anc<-colMeans(model_mvBM$mcmc[(iterations*0.002):nrow(model_mvBM$mcmc),])[3:(3+tree$Nnode-1)]   #These are the ancestral estimates
MCMC_mvBM_logLik<-model_mvBM$mcmc[(iterations*0.002):nrow(model_mvBM$mcmc),tree$Nnode+3]
```

**Plotting rates**

```{r}
pdf("PLE_ancestral_tree.pdf",height=30,width=45)
par(mfrow = c(2, 3))
selenopid_viridasiid<-getTips(tree,findMRCA(tree,c("Selenopidae_Selenops_nesophilus","Viridasiidae_Mahafalytenus_isalo")))
prodidomid_trochanteriid<-getTips(tree,findMRCA(tree,c("Prodidomidae_Molycria_stanisici","Trochanteriidae_Tinytrema_sandy")))
salticid<-getTips(tree,findMRCA(tree,c("Salticidae_Lyssomanes_viridis", "Salticidae_Salticus_scenicus")))
lycosid <- getTips(tree,findMRCA(tree,c("Lycosidae_Schizocosa_ocreata","Psechridae_Fecenia_protensa")))
araneidae <- getTips(tree,findMRCA(tree,c("Araneidae_Eriophora_ravilla", "Araneidae_Phonognatha_graeffei")))
#plot 1
Branches_selenopid_viridasiid<-tail(getEdges(tree,findMRCA(tree,selenopid_viridasiid)),n=1)
Branches_RTA<-tail(getEdges(tree,findMRCA(tree,setdiff(RTA, c(selenopid_viridasiid)))),n=1)
mvBM.getRate(tree,tree_mvBM,branches=Branches_selenopid_viridasiid,sigma2Distr = MCMC_mvBM_sigma2)
mvBM.getRate(tree,tree_mvBM,branches=Branches_RTA,sigma2Distr = MCMC_mvBM_sigma2)

mvBM.plotRate(tree,tree_mvBM,branches=Branches_RTA,sigma2Distr = MCMC_mvBM_sigma2,col="lightgrey",adjust=6,main="", xlim=c(0.003666, 0.00667), ylim = c(0, 25000))
    par(new=TRUE)
mvBM.plotRate(tree,tree_mvBM,branches=Branches_selenopid_viridasiid,sigma2Distr = MCMC_mvBM_sigma2,col="#e20505",adjust=6,main="", xlim=c(0.003666, 0.00667), ylim = c(0, 25000))
legend(x = "topright", legend = c("RTA", "selenopid_viridasiid"), col = c("lightgrey", "#e20505"), lty = 10, lwd = 10, bty = "n")

#plot 2
Branches_salticid<-tail(getEdges(tree,findMRCA(tree,salticid)),n=1)
Branches_RTA<-tail(getEdges(tree,findMRCA(tree,setdiff(RTA, c(salticid)))),n=1)
mvBM.getRate(tree,tree_mvBM,branches=Branches_salticid,sigma2Distr = MCMC_mvBM_sigma2)
mvBM.getRate(tree,tree_mvBM,branches=Branches_RTA,sigma2Distr = MCMC_mvBM_sigma2)

mvBM.plotRate(tree,tree_mvBM,branches=Branches_RTA,sigma2Distr = MCMC_mvBM_sigma2,col="lightgrey",adjust=6,main="", xlim=c(0.0014, 0.00667), ylim = c(0, 66000))
    par(new=TRUE)
mvBM.plotRate(tree,tree_mvBM,branches=Branches_salticid,sigma2Distr = MCMC_mvBM_sigma2,col="#e20505",adjust=6,main="", xlim=c(0.0014, 0.00667), ylim = c(0, 66000))
legend(x = "topright", legend = c("RTA", "salticid"), col = c("lightgrey", "#e20505"), lty = 10, lwd = 10, bty = "n")

#plot 3
Branches_prodidomid_trochanteriid<-tail(getEdges(tree,findMRCA(tree,prodidomid_trochanteriid)),n=1)
Branches_RTA<-tail(getEdges(tree,findMRCA(tree,setdiff(RTA, c(prodidomid_trochanteriid)))),n=1)
mvBM.getRate(tree,tree_mvBM,branches=Branches_prodidomid_trochanteriid,sigma2Distr = MCMC_mvBM_sigma2)
mvBM.getRate(tree,tree_mvBM,branches=Branches_RTA,sigma2Distr = MCMC_mvBM_sigma2)

mvBM.plotRate(tree,tree_mvBM,branches=Branches_RTA,sigma2Distr = MCMC_mvBM_sigma2,col="lightgrey",adjust=6,main="", xlim=c(0.001210, 0.00667), ylim = c(0, 75000))
    par(new=TRUE)
mvBM.plotRate(tree,tree_mvBM,branches=Branches_prodidomid_trochanteriid,sigma2Distr = MCMC_mvBM_sigma2,col="#e20505",adjust=6,main="", xlim=c(0.001210, 0.00667), ylim = c(0, 75000))
legend(x = "topright", legend = c("RTA", "prodidomid_trochanteriid"), col = c("lightgrey", "#e20505"), lty = 10, lwd = 10, bty = "n")

#Plot 4
Branches_lycosid<-tail(getEdges(tree,findMRCA(tree,lycosid)),n=1)
Branches_Oval<-tail(getEdges(tree,findMRCA(tree,setdiff(Oval, c(lycosid)))),n=1)
mvBM.getRate(tree,tree_mvBM,branches=Branches_lycosid,sigma2Distr = MCMC_mvBM_sigma2)
mvBM.getRate(tree,tree_mvBM,branches=Branches_Oval,sigma2Distr = MCMC_mvBM_sigma2)

mvBM.plotRate(tree,tree_mvBM,branches=Branches_Oval,sigma2Distr = MCMC_mvBM_sigma2,col="lightgrey",adjust=6,main="", xlim=c(0.00162, 0.001675), ylim = c(0, 61000))
    par(new=TRUE)
mvBM.plotRate(tree,tree_mvBM,branches=Branches_lycosid,sigma2Distr = MCMC_mvBM_sigma2,col="#e20505",adjust=6,main="", xlim=c(0.001465, 0.001675), ylim = c(0, 61000))
legend(x = "topright", legend = c("Oval", "lycosid"), col = c("lightgrey", "#e20505"), lty = 10, lwd = 10, bty = "n")

#Plot 5
Branches_Aranoidea<-tail(getEdges(tree,findMRCA(tree,Aranoidea)),n=1)
Branches_araneidae<-tail(getEdges(tree,findMRCA(tree,setdiff(Aranoidea, c(araneidae)))),n=1)
mvBM.getRate(tree,tree_mvBM,branches=Branches_Aranoidea,sigma2Distr = MCMC_mvBM_sigma2)
mvBM.getRate(tree,tree_mvBM,branches=Branches_araneidae,sigma2Distr = MCMC_mvBM_sigma2)

mvBM.plotRate(tree,tree_mvBM,branches=Branches_Aranoidea,sigma2Distr = MCMC_mvBM_sigma2,col="lightgrey",adjust=6,main="", xlim=c(0.001315, 0.001355), ylim = c(0, 70000))
    par(new=TRUE)
mvBM.plotRate(tree,tree_mvBM,branches=Branches_araneidae,sigma2Distr = MCMC_mvBM_sigma2,col="#e20505",adjust=6,main="", xlim=c(0.001315, 0.001355), ylim = c(0, 70000))
legend(x = "topright", legend = c("Aranoidea", "araneidae"), col = c("lightgrey", "#e20505"), lty = 10, lwd = 10, bty = "n")
    dev.off()

```

#### **logPME**

```{r}
data = data_all%>% drop_na(PME)
xx <- c(data$terminal)
common_names <- intersect(tree_all$tip.label, xx)
tree <- drop.tip(tree_all, tree_all$tip.label[!tree_all$tip.label %in% common_names])
rownames(data) <- data$terminal
Names <- rownames(data)
dat <- log(data$PMEmm)
names(dat) <- Names
mvBMresults<-mvBM(dat,tree,ace(dat,tree,method="REML")$sigma2[1])
# calculate rescaled branch lengths using mvBM
tree_mvBM<-tree
tree_mvBM$edge.length<-mvBMresults$rBL #create new tree with rescaled branch lengths
plot(tree_mvBM,cex=0.2)
#setting seed
set.seed(0)
iterations<-1000000
model_mvBM<-anc.Bayes(tree_mvBM,dat,ngen=iterations)  
# 'anc.Bayes' requires the 'phytools' package; this may take a few minutes
MCMC_mvBM_sigma2<-model_mvBM$mcmc[(iterations*0.002):nrow(model_mvBM$mcmc),2] 
MCMC_mvBM_anc<-colMeans(model_mvBM$mcmc[(iterations*0.002):nrow(model_mvBM$mcmc),])[3:(3+tree$Nnode-1)]   #These are the ancestral estimates
MCMC_mvBM_logLik<-model_mvBM$mcmc[(iterations*0.002):nrow(model_mvBM$mcmc),tree$Nnode+3]

```

**Plotting rates**

```{r}
philodromid<-getTips(tree,findMRCA(tree,c("Philodromidae_Philodromus_aureolus","Philodromidae_Thanatus_formicinus")))
salticid<-getTips(tree,findMRCA(tree,c("Salticidae_Lyssomanes_viridis", "Salticidae_Salticus_scenicus")))
prodidomid_phrurolithid <-getTips(tree,findMRCA(tree,c("Prodidomidae_Molycria_stanisici","Phrurolithidae_Otacilia_sp_HaTinh")))
thomisid<-getTips(tree,findMRCA(tree,c("Thomisidae_Runcinia_albostriata","Thomisidae_Borboropactus_bituberculatus")))
oxyopid <- getTips(tree,findMRCA(tree,c("Oxyopidae_Oxyopes_salticus","Oxyopidae_Peucetia_viridans")))
lycosid <- getTips(tree,findMRCA(tree,c("Lycosidae_Schizocosa_ocreata","Lycosidae_Pirata_subpiraticus")))
pisaurid_psechrid <- getTips(tree,findMRCA(tree,c("Pisauridae_Hala_paulyi","Psechridae_Fecenia_protensa")))
zoropsid <- getTips(tree,findMRCA(tree,c("Zoropsidae_Uliodon_sp_CG127","Zoropsidae_Phanotea_sathegyna")))
cycloctenid <- getTips(tree,findMRCA(tree,c("Cycloctenidae_Cycloctenus_fugax","Cycloctenidae_Toxopsiella_minuta")))
deinopid <- getTips(tree,findMRCA(tree,c("Deinopidae_Menneus_camelus","Deinopidae_Deinopis_spinosa")))
pdf("PME_ancestral_tree.pdf",height=30,width=45)
par(mfrow = c(3, 4))
#plot 1
Branches_philodromid<-tail(getEdges(tree,findMRCA(tree,philodromid)),n=1)
Branches_RTA<-tail(getEdges(tree,findMRCA(tree,setdiff(RTA, c(philodromid)))),n=1)
mvBM.getRate(tree,tree_mvBM,branches=Branches_philodromid,sigma2Distr = MCMC_mvBM_sigma2)
mvBM.getRate(tree,tree_mvBM,branches=Branches_RTA,sigma2Distr = MCMC_mvBM_sigma2)

mvBM.plotRate(tree,tree_mvBM,branches=Branches_RTA,sigma2Distr = MCMC_mvBM_sigma2,col="lightgrey",adjust=6,main="", xlim=c(0.001295, 0.00174), ylim = c(0, 41000))
    par(new=TRUE)
mvBM.plotRate(tree,tree_mvBM,branches=Branches_philodromid,sigma2Distr = MCMC_mvBM_sigma2,col="#eded14",adjust=6,main="",xlim=c(0.001295,  0.00174), ylim = c(0, 41000))
legend(x = "topright", legend = c("RTA", "philodromid"), col = c("lightgrey", "#eded14"), lty = 10, lwd = 10, bty = "n")
#plot 2
Branches_salticid<-tail(getEdges(tree,findMRCA(tree,salticid)),n=1)
Branches_RTA<-tail(getEdges(tree,findMRCA(tree,setdiff(RTA, c(salticid)))),n=1)
mvBM.getRate(tree,tree_mvBM,branches=Branches_salticid,sigma2Distr = MCMC_mvBM_sigma2)
mvBM.getRate(tree,tree_mvBM,branches=Branches_RTA,sigma2Distr = MCMC_mvBM_sigma2)

mvBM.plotRate(tree,tree_mvBM,branches=Branches_RTA,sigma2Distr = MCMC_mvBM_sigma2,col="lightgrey",adjust=6,main="", xlim=c(0.00164, 0.00188), ylim = c(0, 32100))
    par(new=TRUE)
mvBM.plotRate(tree,tree_mvBM,branches=Branches_salticid,sigma2Distr = MCMC_mvBM_sigma2,col="#eded14",adjust=6,main="",xlim=c(0.00164, 0.00188), ylim = c(0, 32100))
legend(x = "topright", legend = c("RTA", "salticid"), col = c("lightgrey", "#eded14"), lty = 10, lwd = 10, bty = "n")
#Plot 3
Branches_prodidomid_phrurolithid<-tail(getEdges(tree,findMRCA(tree,prodidomid_phrurolithid)),n=1)
Branches_RTA<-tail(getEdges(tree,findMRCA(tree,setdiff(RTA, c(prodidomid_phrurolithid)))),n=1)
mvBM.getRate(tree,tree_mvBM,branches=Branches_prodidomid_phrurolithid,sigma2Distr = MCMC_mvBM_sigma2)
mvBM.getRate(tree,tree_mvBM,branches=Branches_RTA,sigma2Distr = MCMC_mvBM_sigma2)

mvBM.plotRate(tree,tree_mvBM,branches=Branches_RTA,sigma2Distr = MCMC_mvBM_sigma2,col="lightgrey",adjust=6,main="", xlim=c(0.00164, 0.00255), ylim = c(0, 33000))
    par(new=TRUE)
mvBM.plotRate(tree,tree_mvBM,branches=Branches_prodidomid_phrurolithid,sigma2Distr = MCMC_mvBM_sigma2,col="#eded14",adjust=6,main="", xlim=c(0.00164, 0.00255), ylim = c(0, 33000))
legend(x = "topright", legend = c("RTA", "prodidomid_phrurolithid"), col = c("lightgrey", "#eded14"), lty = 10, lwd = 10, bty = "n")

#Plot 4
Branches_thomisid<-tail(getEdges(tree,findMRCA(tree,thomisid)),n=1)
Branches_Oval<-tail(getEdges(tree,findMRCA(tree,setdiff(Oval, c(thomisid)))),n=1)
mvBM.getRate(tree,tree_mvBM,branches=Branches_thomisid,sigma2Distr = MCMC_mvBM_sigma2)
mvBM.getRate(tree,tree_mvBM,branches=Branches_Oval,sigma2Distr = MCMC_mvBM_sigma2)

mvBM.plotRate(tree,tree_mvBM,branches=Branches_Oval,sigma2Distr = MCMC_mvBM_sigma2,col="lightgrey",adjust=6,main="", xlim=c(0.00141, 0.0085), ylim = c(0, 38500))
    par(new=TRUE)
mvBM.plotRate(tree,tree_mvBM,branches=Branches_thomisid,sigma2Distr = MCMC_mvBM_sigma2,col="#eded14",adjust=6,main="",xlim=c(0.00141, 0.0085), ylim = c(0, 38500))
legend(x = "topright", legend = c("Oval", "thomisid"), col = c("lightgrey", "#eded14"), lty = 10, lwd = 10, bty = "n")

#Plot 5
Branches_oxypodid<-tail(getEdges(tree,findMRCA(tree,oxyopid)),n=1)
Branches_Oval<-tail(getEdges(tree,findMRCA(tree,setdiff(Oval, c(oxyopid)))),n=1)
mvBM.getRate(tree,tree_mvBM,branches=Branches_oxypodid,sigma2Distr = MCMC_mvBM_sigma2)
mvBM.getRate(tree,tree_mvBM,branches=Branches_Oval,sigma2Distr = MCMC_mvBM_sigma2)

mvBM.plotRate(tree,tree_mvBM,branches=Branches_Oval,sigma2Distr = MCMC_mvBM_sigma2,col="lightgrey",adjust=6,main="", xlim=c(0.00125, 0.00150), ylim = c(0, 42000))
    par(new=TRUE)
mvBM.plotRate(tree,tree_mvBM,branches=Branches_oxypodid,sigma2Distr = MCMC_mvBM_sigma2,col="#eded14",adjust=6,main="",xlim=c(0.00125, 0.0015), ylim = c(0, 42000))
legend(x = "topright", legend = c("Oval", "oxypodid"), col = c("lightgrey", "#eded14"), lty = 10, lwd = 10, bty = "n")

#Plot 6
Branches_lycosid<-tail(getEdges(tree,findMRCA(tree,lycosid)),n=1)
Branches_Oval<-tail(getEdges(tree,findMRCA(tree,setdiff(Oval, c(lycosid)))),n=1)
mvBM.getRate(tree,tree_mvBM,branches=Branches_lycosid,sigma2Distr = MCMC_mvBM_sigma2)
mvBM.getRate(tree,tree_mvBM,branches=Branches_Oval,sigma2Distr = MCMC_mvBM_sigma2)

mvBM.plotRate(tree,tree_mvBM,branches=Branches_Oval,sigma2Distr = MCMC_mvBM_sigma2,col="lightgrey",adjust=6,main="", xlim=c(0.00141, 0.0056), ylim = c(0, 38000))
    par(new=TRUE)
mvBM.plotRate(tree,tree_mvBM,branches=Branches_lycosid,sigma2Distr = MCMC_mvBM_sigma2,col="#eded14",adjust=6,main="", xlim=c(0.00529, 0.0056), ylim = c(0, 38000))
legend(x = "topright", legend = c("Oval", "lycosid"), col = c("lightgrey", "#eded14"), lty = 10, lwd = 10, bty = "n")

#Plot 7
Branches_pisaurid_psechrid<-tail(getEdges(tree,findMRCA(tree,pisaurid_psechrid)),n=1)
Branches_Oval<-tail(getEdges(tree,findMRCA(tree,setdiff(Oval, c(pisaurid_psechrid)))),n=1)
mvBM.getRate(tree,tree_mvBM,branches=Branches_pisaurid_psechrid,sigma2Distr = MCMC_mvBM_sigma2)
mvBM.getRate(tree,tree_mvBM,branches=Branches_Oval,sigma2Distr = MCMC_mvBM_sigma2)

mvBM.plotRate(tree,tree_mvBM,branches=Branches_Oval,sigma2Distr = MCMC_mvBM_sigma2,col="lightgrey",adjust=6,main="", xlim=c(0.00141, 0.00169), ylim = c(0, 37000))
    par(new=TRUE)
mvBM.plotRate(tree,tree_mvBM,branches=Branches_pisaurid_psechrid,sigma2Distr = MCMC_mvBM_sigma2,col="#eded14",adjust=6,main="",xlim=c(0.00141, 0.00169), ylim = c(0, 37000))
legend(x = "topright", legend = c("Oval", "pisaurid_psechrid"), col = c("lightgrey", "#eded14"), lty = 10, lwd = 10, bty = "n")

#Plot 8
Branches_zoropsid<-tail(getEdges(tree,findMRCA(tree,zoropsid)),n=1)
Branches_Oval<-tail(getEdges(tree,findMRCA(tree,setdiff(Oval, c(zoropsid)))),n=1)
mvBM.getRate(tree,tree_mvBM,branches=Branches_zoropsid,sigma2Distr = MCMC_mvBM_sigma2)
mvBM.getRate(tree,tree_mvBM,branches=Branches_Oval,sigma2Distr = MCMC_mvBM_sigma2)

mvBM.plotRate(tree,tree_mvBM,branches=Branches_Oval,sigma2Distr = MCMC_mvBM_sigma2,col="lightgrey",adjust=6,main="", xlim=c(0.0027, 0.00517), ylim = c(0, 20000))
    par(new=TRUE)
mvBM.plotRate(tree,tree_mvBM,branches=Branches_zoropsid,sigma2Distr = MCMC_mvBM_sigma2,col="#eded14",adjust=6,main="",xlim=c(0.0027, 0.00517), ylim = c(0, 20000))
legend(x = "topright", legend = c("Oval", "zoropsid"), col = c("lightgrey", "#eded14"), lty = 10, lwd = 10, bty = "n")

#plot 9
Branches_cycloctenid<-tail(getEdges(tree,findMRCA(tree,cycloctenid)),n=1)
Branches_Marronoid<-tail(getEdges(tree,findMRCA(tree,setdiff(Marronoid, c(cycloctenid)))),n=1)
mvBM.getRate(tree,tree_mvBM,branches=Branches_cycloctenid,sigma2Distr = MCMC_mvBM_sigma2)
mvBM.getRate(tree,tree_mvBM,branches=Branches_Marronoid,sigma2Distr = MCMC_mvBM_sigma2)

mvBM.plotRate(tree,tree_mvBM,branches=Branches_Marronoid,sigma2Distr = MCMC_mvBM_sigma2,col="lightgrey",adjust=6,main="", xlim=c(0.00525, 0.0178), ylim = c(0, 10000))
    par(new=TRUE)
mvBM.plotRate(tree,tree_mvBM,branches=Branches_cycloctenid,sigma2Distr = MCMC_mvBM_sigma2,col="#eded14",adjust=6,main="", xlim=c(0.00525, 0.0178), ylim = c(0, 10000))
legend(x = "topright", legend = c("Marronoid", "cycloctenid"), col = c("lightgrey", "#eded14"), lty = 10, lwd = 10, bty = "n")
#Plot 10
Branches_deinopid<-tail(getEdges(tree,findMRCA(tree,deinopid)),n=1)
Branches_Entelegynes<-tail(getEdges(tree,findMRCA(tree,setdiff(Entelegynes, c(deinopid)))),n=1)
mvBM.getRate(tree,tree_mvBM,branches=Branches_deinopid,sigma2Distr = MCMC_mvBM_sigma2)
mvBM.getRate(tree,tree_mvBM,branches=Branches_Entelegynes,sigma2Distr = MCMC_mvBM_sigma2)
mvBM.plotRate(tree,tree_mvBM,branches=Branches_Entelegynes,sigma2Distr = MCMC_mvBM_sigma2,col="lightgrey",adjust=6,main="",xlim=c(0.00130, 0.00204), ylim = c(0, 41000))
    par(new=TRUE)
mvBM.plotRate(tree,tree_mvBM,branches=Branches_deinopid,sigma2Distr = MCMC_mvBM_sigma2,col="#eded14",adjust=6,main="",xlim=c(0.00130, 0.00204), ylim = c(0, 41000))
legend(x = "topright", legend = c("Entelegynes", "deinopid"), col = c("lightgrey", "#eded14"), lty = 10, lwd = 10, bty = "n")
    dev.off()

```
````
