## Supplementary material for "Complex allometric relationships and ecological factors shape the development and evolution of eye size in the modular visual system of spiders": Chong et al. supplementary data.docx

**Chong et al. Supplementary materials	(* indicates files uploaded separately)**

**Datasets and analyses**

*S1. Specimen numbers*

*S2. Ln_total.csv**

Raw data for static allometry.

*S3. WolffData-Dec23.txt**

Curated dataset from Wolff et al. 2022 (Systematic Biology), used in phylogenetic allometric analyses.

*S4. WolffDataNAs-Dec23.txt**

Curated dataset from Wolff et al. 2022 (Systematic Biology), used in phylogenetic allometric analyses.

*S5. WolffRed22Sept.nwk**

Curated phylogeny from Wolff et al. 2022 (Systematic Biology), used in phylogenetic allometric analyses.

*S6. static_allometry.Rmd**

R analysis for raw data and static allometry results.

*S7. Residuals.csv**

Calculated residuals from static allometric models generated for each eye pair of each species.

*S8. PGLS.Rmd**

R analysis of phylogenetic allometry using curated dataset from Wolff et al. 2022.

*S9. Bayou Rmd.zip**

Contains 4 folders (AME, ALE, PME, PLE) containing R markdowns to run Bayesian multi-regime OU modelling.

*S10. ancestral_state_analysis.Rmd**

R analysis for ancestral state reconstruction results.

**Figures**

**Static allometry**

*SF1. Carapace.pdf*

Carapace widths measured across 1098 spiders from eight families and 39 species. Measurements are pooled by family.

*SF2. Residuals.pdf*

Residuals from static allometric models generated for each eye pair of each species, pooled across families.

**Phylogenetic allometry**

SF3-SF6

Bayesian multi-regime OU modelling trace outputs for all four eye pair models generated from 10 chains. Curated dataset from Wolff et al. (2022).

*SF3. bayouAllometry_withPrimer_AME_CW_1e+06_trace.pdf*

*SF4. bayouAllometry_withPrimer_ALE_CW_1e+06_trace.pdf*

*SF5. bayouAllometry_withPrimer_PME_CW_1e+06_trace.pdf*

*SF6. bayouAllometry_withPrimer_PLE_CW_1e+06_trace.pdf*

SF7-SF10

Phylogenetic tree predicted through Bayesian multi-regime Ornstein–Uhlenbeck (OU) modelling with a posterior probability of 0.2. Curated dataset from Wolff et al. (2022).

*SF7. bayouAllometry_AME_CW_1e+06_PP_0.2.pdf**

*SF8. bayouAllometry_ALE_CW_1e+06_PP_0.2.pdf**

*SF9. bayouAllometry_PLE_CW_1e+06_PP_0.2.pdf**

*SF10. bayouAllometry_PME_CW_1e+06_PP_0.2.pdf**

**Ancestral state reconstructions**

SF11-SF14 Ancestral state reconstructions of log-transformed absolute (left) and relative (right) eye diameters across 474 species from a comparative dataset extracted from Wolff et al. 2022 (Systematic Biology).

*SF11. ASR-AME.pdf*

*SF12. ASR-ALE.pdf*

*SF13. ASR-PME.pdf*

*SF14. ASR-PLE.pdf*

**Supplementary Tables (Supplementary_tables.xlsx)**

Table S1 Results of Tukey HSD tests comparing absolute eye diameter between eye pairs within three families: Linyphiidae, Lycosidae, and Pholcidae.

Table S2 Results of Dunn’s tests comparing absolute eye diameter between eye pairs within five families: Araneidae, Gnaphosidae, Salticidae, Theridiidae, and Thomisidae.

Table S3 Results of OLS analyses of eye diameter vs. carapace width for four eye pairs across 39 species of spider.

Table S4 Summary of OLS analyses of eye diameter vs. carapace width for 39 species of spiders. Colours indicate significance and gradient of allometric slopes.

Table S5 R-squared values for slopes generated by OLS analyses of eye diameter vs. carapace width for 39 species of spiders.

Table S6 Results of linear mixed models of static visual allometry in 39 species of spiders. Carapace width, eye pair identity, and their interaction were included as fixed effects, and the individual measured was included as a random effect.

Table S7 Allometric slope estimates for each eye pair, generated by linear mixed models of static visual allometry in 39 species of spiders. Carapace width, eye pair identity, and their interaction were included as fixed effects, and the individual measured was included as a random effect.

Table S8 Pairwise comparisons of allometric slopes generated by linear mixed models of static visual allometry in 39 species of spiders, performed using emtrends. Carapace width, eye pair identity, and their interaction were included as fixed effects, and the individual measured was included as a random effect.

Table S9 Pairwise comparisons of intercepts generated by linear mixed models of static visual allometry in 39 species of spiders, performed using emmeans. Carapace width, eye pair identity, and their interaction were included as fixed effects, and the individual measured was included as a random effect.

Table S10 Results of linear mixed models of static visual allometry in eight families of spiders. Carapace width, eye pair identity, and their interaction were included as fixed effects, and the species and individual measured were included as random effects.

Table S11 Pairwise comparisons of allometric slopes generated by linear mixed models of static visual allometry in eight families of spiders, performed using emtrends. Carapace width, eye pair identity, and their interaction were included as fixed effects, and the species and individual measured were included as random effects.

Table S12 Pairwise comparisons of intercepts generated by linear mixed models of static visual allometry in 39 species of spiders, performed using emmeans. Carapace width, eye pair identity, and their interaction were included as fixed effects, and the species and individual measured were included as random effects.

Table S13 Comparison of carapace width and eye diameter before and after allometric shifts as predicted by Bayou, with estimations as to which of these drove the predicted shift.

Table S14 Reconstructed ancestral relative and log-transformed eye diameters at the base of selected major clades of spiders.

Table S15 Predicted evolutionary rates of eye diameter, before and after allometric shifts within clades as predicted by Bayou.

***Supplementary materials***

***S1. Specimen numbers***

**Araneidae**

*Agalenatea redii*	OUMNH British Spider Colln Jar #3200

*Araneus diadematus*	OUMNH British Spider Colln Jar #3100

*Araniella curcurbita*	OUMNH British Spider Colln Jar #3240

*Larinioides cornutus*	OUMNH British Spider Colln Jar #3150

*Zygiella atrica*	OUMNH British Spider Colln Jar #3370

**Gnaphosidae**

*Drassodes lapidosus*	OUMNH British Spider Colln Jar #350

*Gnaphosa lugubris*	OUMNH British Spider Colln Jar #560

*Haplodrassus signifer*	OUMNH British Spider Colln Jar #390

*Scotophaeus blackwalli*	OUMNH British Spider Colln Jar #440

*Zelotes latreilli*	OUMNH British Spider Colln Jar #520

**Pholcidae**

*Artema atlantica*	OUMNH World Spider Colln Jar #43

*Crossopriza lyoni*	OUMNH World Spider Colln Jar #45

*Pholcus phalangioides*	OUMNH World Spider Colln Jar #47,
OUMNH British Spider Colln Jar #330

*Smeringopus pallidus*	OUMNH World Spider Colln Jar #49

**Linyphiidae**

*Centromerus sylvaticus*	OUMNH British Spider Colln Jar #5250

*Erigone atra*	OUMNH British Spider Colln Jar #4760

*Lephthyphantes minutus*	OUMNH British Spider Colln Jar #5660

*Linyphius triangularis*	OUMNH British Spider Colln Jar #5850

*Walckenaeria acuminata*	OUMNH British Spider Colln Jar #3460

**Lycosidae**

*Arctosa perita*	OUMNH British Spider Colln Jar #2050

*Pardosa amentata*	OUMNH British Spider Colln Jar #1860

*Pirata piraticus*	OUMNH British Spider Colln Jar #2090

*Trochosa terricola*	OUMNH British Spider Colln Jar #2020

*Xerolycosa miniata*	OUMNH British Spider Colln Jar #1950

**Salticidae**

*Evarcha fulcata*	OUMNH British Spider Colln Jar #1680,
OUMNH World Spider Colln Jar #458

*Euophrys frontalis*	OUMNH British Spider Colln Jar #1560

*Heliophanus flavipes*	OUMNH British Spider Colln Jar #1470

*Mogrus mathisi*	OUMNH-2005-030, OUMNH World Spider Colln Jar #474

*Neon reticulatus*	OUMNH British Spider Colln Jar #1540

**Theridiidae**

*Parasteatoda tepidariorum*	OUMNH British Spider Colln Jar #2650

**Thomisidae**

*Diaea dorsata*	OUMNH British Spider Colln Jar #1060

*Misumena vatia*	OUMNH British Spider Colln Jar #1070

*Ozyptila atomaria*	OUMNH British Spider Colln Jar #1280

*Thomisus onustus*	OUMNH British Spider Colln Jar #1050

*Xysticus cristatus*	OUMNH British Spider Colln Jar #1090

*[ Ln_total.csv]*

**S2 Raw data for static allometry**. Measurements of eye diameter and carapace width taken from specimens drawn from the British Spider Collection at the Oxford University Museum of Natural History (see S1 for accession numbers).

*[Residuals.csv]*

**S3 Calculated residuals from static allometric models generated for each eye pair of each species.** Allometric slopes were fitted to each dataset using OLS analysis. Residuals for each observed datapoint were calculated based on its distance from the fitted slope.

*[WolffDataNAs-Dec23.txt]*

**S4 Curated dataset from Wolff et al. 2022 (Systematic Biology), used in phylogenetic allometric analyses.** The original dataset contained morphometric data (n=1968 entries) including eye diameters and carapace widths, ecological data (n=829) including hunting guilds. We extracted entries with all eyes, carapace width, and ecological data recorded. Where duplicate entries were present for a given species, we took mean eye diameters and carapace widths. Species missing from the phylogeny were removed. The resulting dataset contains single entries for 474 species.

*[WolffRed22Sept.nwk]*

**S5 Curated phylogeny from Wolff et al. 2022 (Systematic Biology), used in phylogenetic allometric analyses.** The original phylogeny was a time-calibrated maximum-likelihood tree of 828 species. Taxa missing from the curated dataset (see S5) and corresponding internal nodes were removed using the *drop tips* function in R. The resulting phylogeny contains 474 species.

*[S6-S10 R markdown files for relevant analyses. See README.txt*]*

***Supplementary figures***

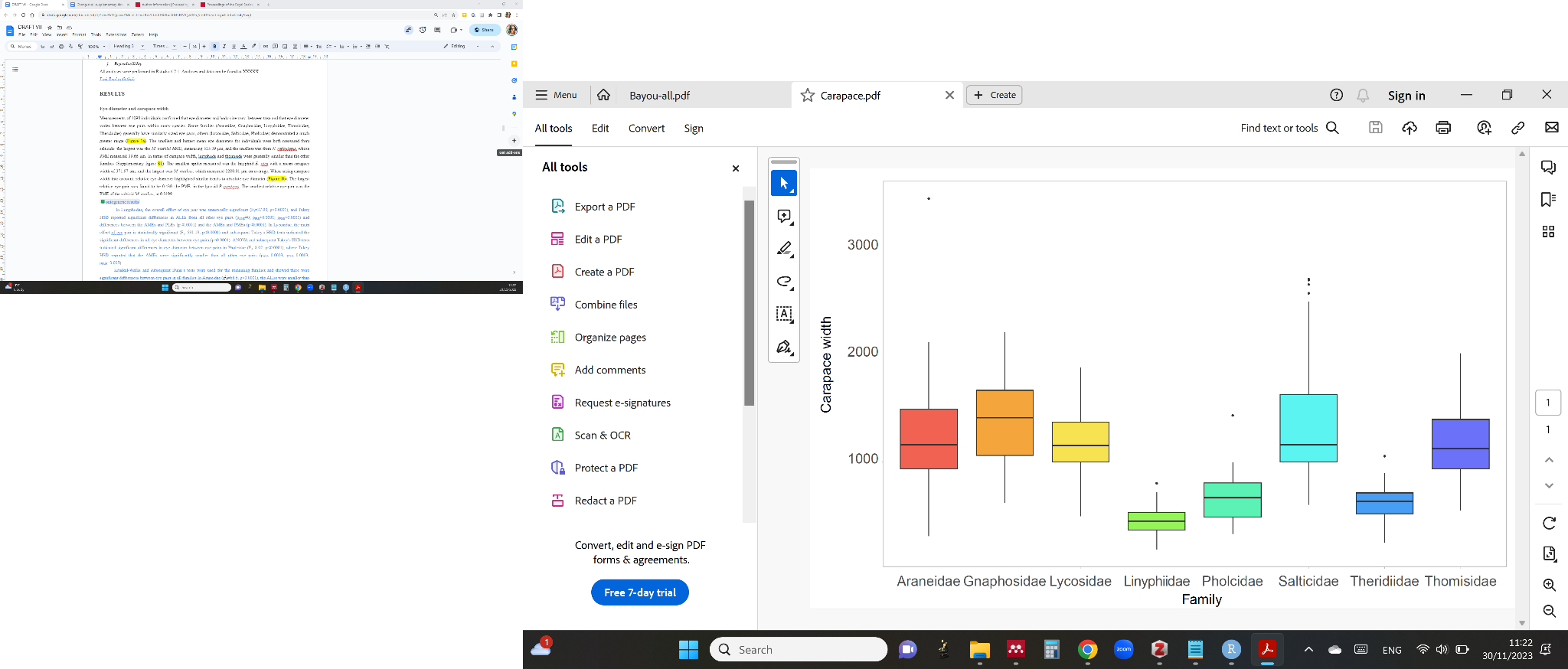

**Figure SF1 Carapace width measurements (in μm) of 1098 individual spiders from 39 species, pooled by family.**

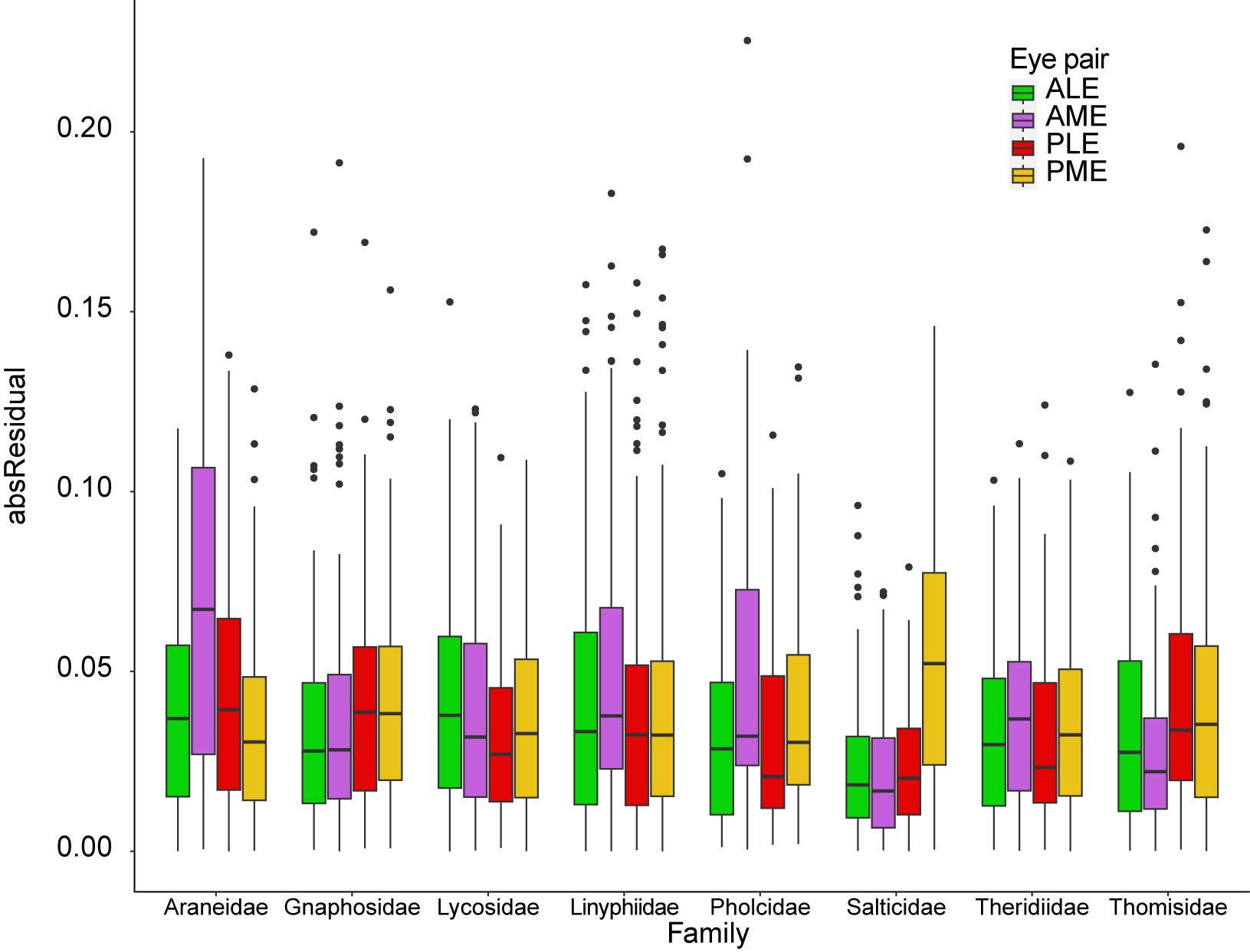

**Figure SF2 Absolute residuals for static allometric models, pooled by family. Residuals were extracted from individual OLS analyses for each eye pair and each species.**

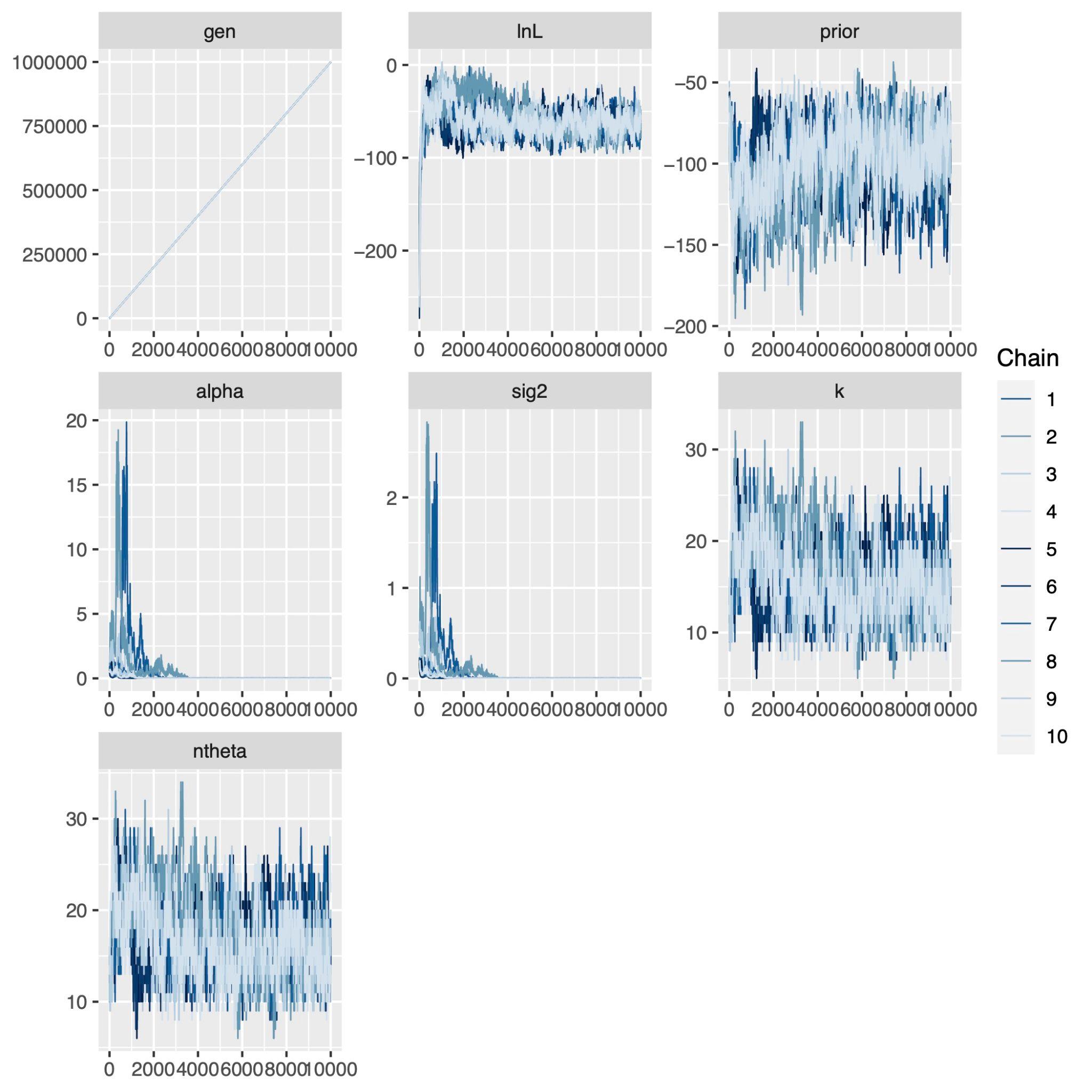

**Figure SF3 Trace output for the AME model using Bayesian multiregime OU modeling, generated from 10 chains.** Curated dataset from Wolff et al. (2022).

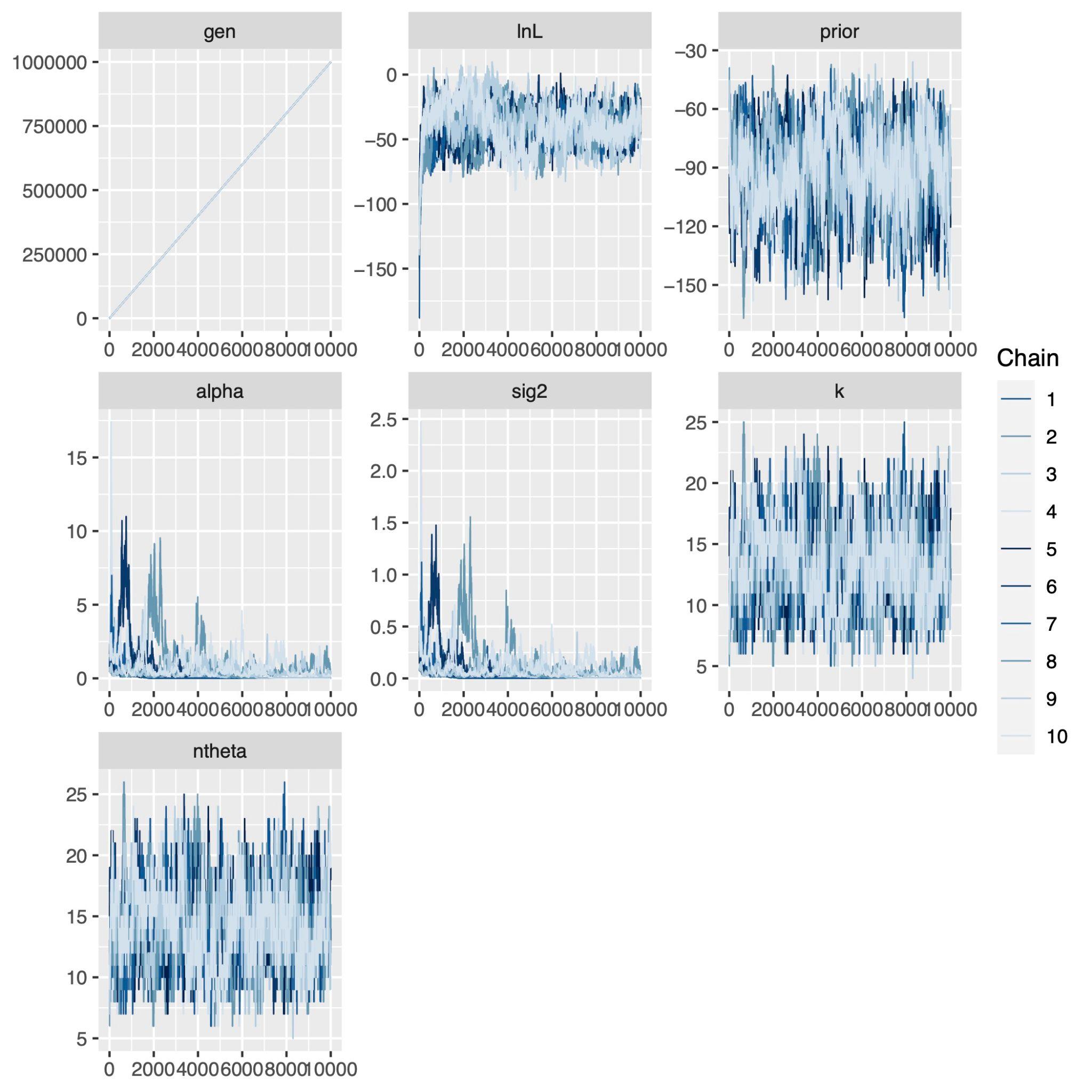

**Figure SF4 Trace output for the ALE model using Bayesian multiregime OU modeling, generated from 10 chains.** Curated dataset from Wolff et al. (2022).

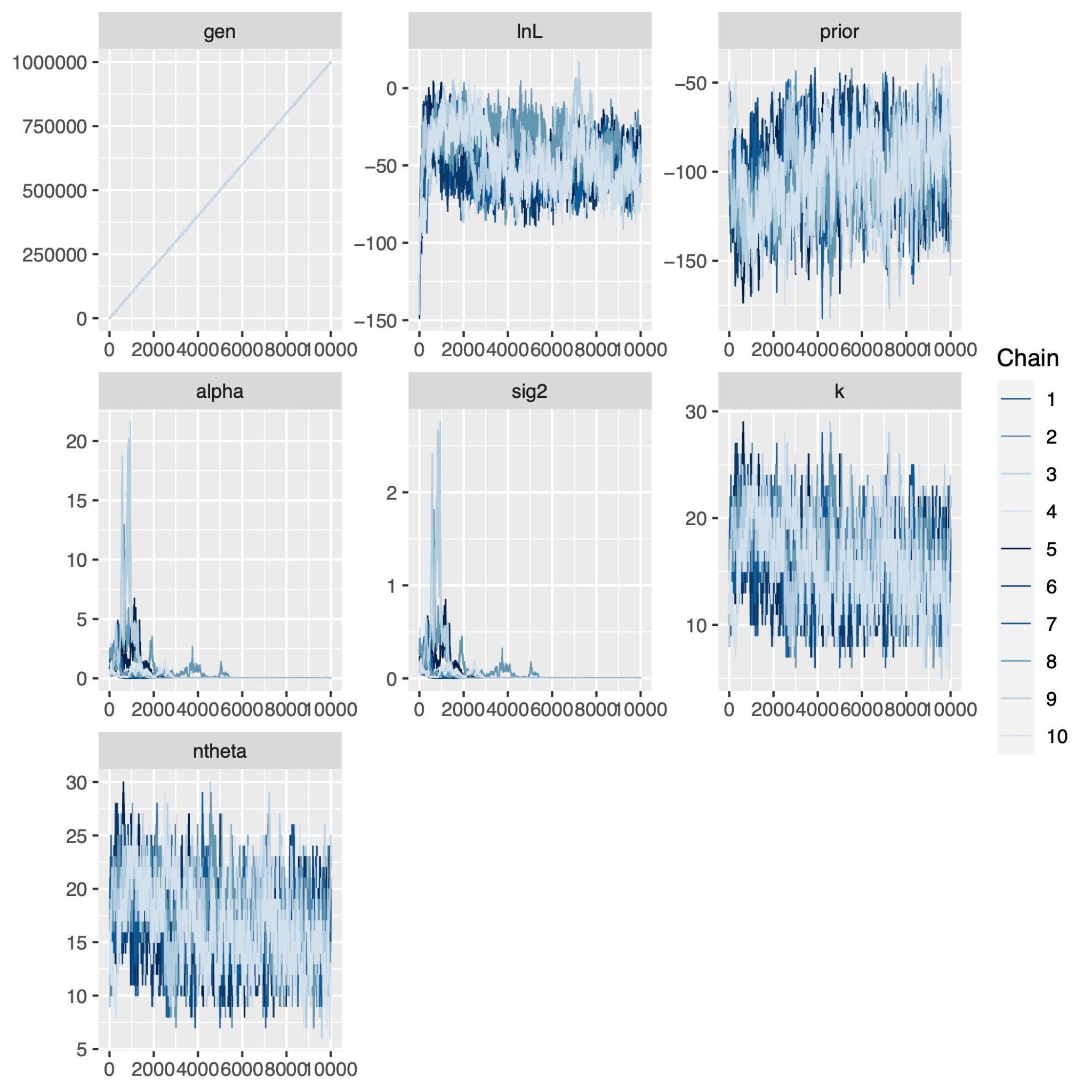

**Figure SF5 Trace output for the PME using Bayesian multiregime OU modeling, generated from 10 chains.** Curated dataset from Wolff et al. (2022).

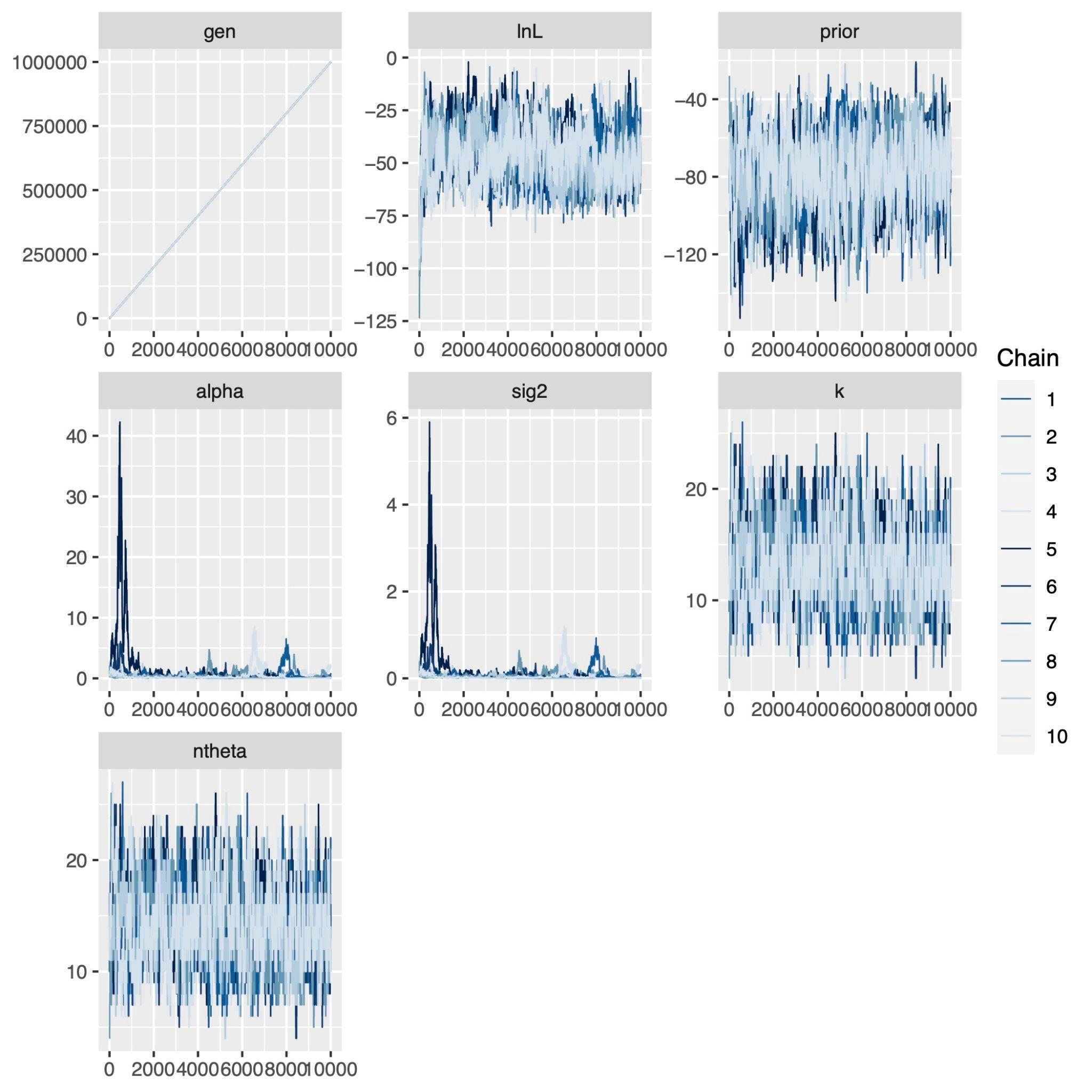

**Figure SF6 Trace output for the PLE model using Bayesian multiregime OU modeling, generated from 10 chains.** Curated dataset from Wolff et al. (2022).

*[bayouAllometry_AME_CW_1e+06_PP_0.2.pdf*]*

**Figure SF7 Prediction of allometric shifts in AME diameter, generated by Bayesian multiregime Ornstein–Uhlenbeck (OU) modelling. The associated posterior probability is 0.2.** Curated dataset from Wolff et al. (2022).

*[bayouAllometry_ALE_CW_1e+06_PP_0.2.pdf*]*

**Figure SF8 Prediction of allometric shifts in ALE diameter, generated by Bayesian multiregime Ornstein–Uhlenbeck (OU) modelling. The associated posterior probability is 0.2.** Curated dataset from Wolff et al. (2022).

*[bayouAllometry_PME_CW_1e+06_PP_0.2.pdf*]*

**Figure SF9 Prediction of allometric shifts in PME diameter, generated by Bayesian multiregime Ornstein–Uhlenbeck (OU) modelling. The associated posterior probability is 0.2.** Curated dataset from Wolff et al. (2022).

*[bayouAllometry_PLE_CW_1e+06_PP_0.2.pdf*]*

**Figure SF10 Prediction of allometric shifts in PLE diameter, generated by Bayesian multiregime Ornstein–Uhlenbeck (OU) modelling. The associated posterior probability is 0.2.** Curated dataset from Wolff et al. (2022).

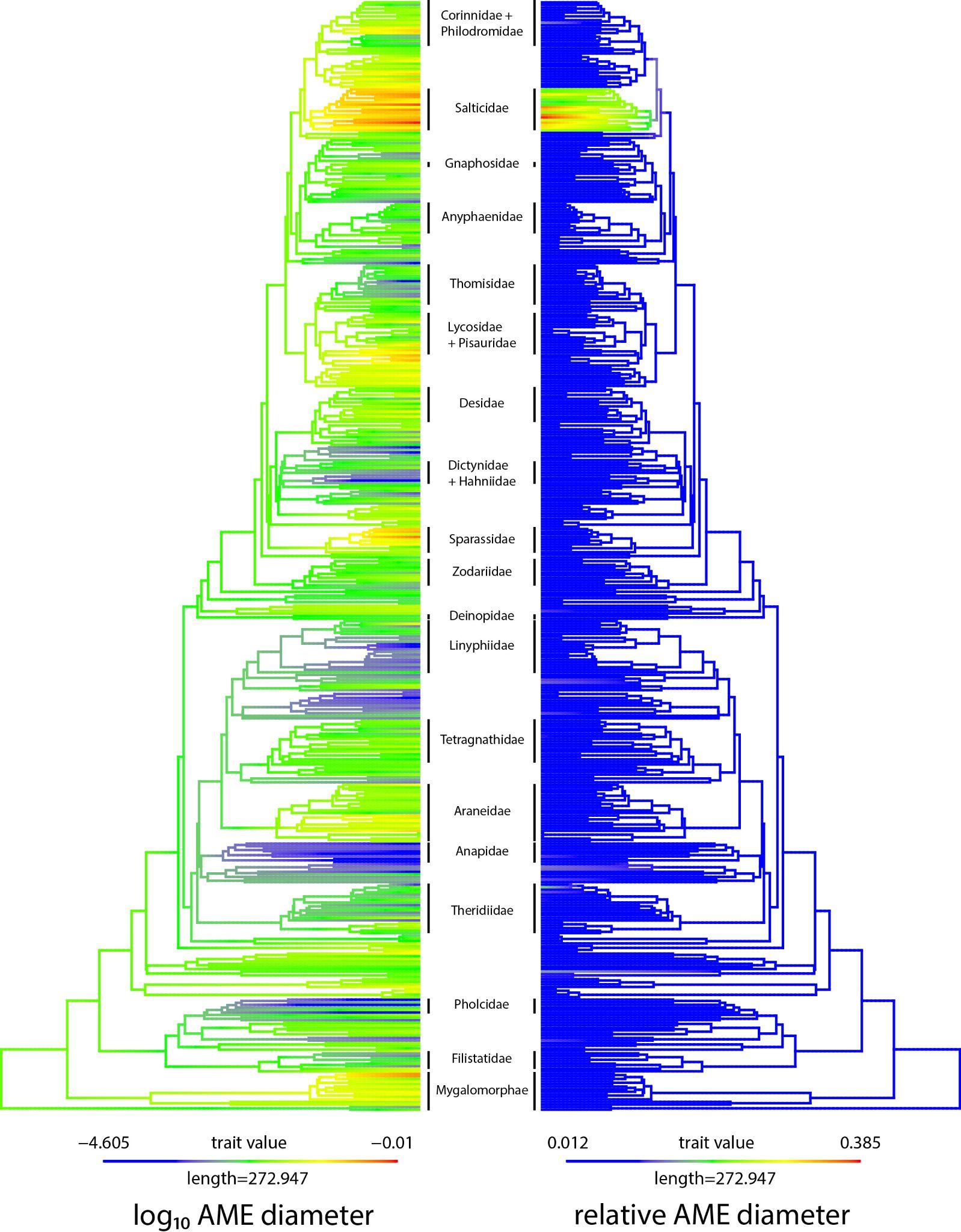

**Figure SF11 Ancestral state reconstruction of log-transformed and relative AME diameters across spiders.** Curated dataset from Wolff et al. (2022). Position of major families labelled.

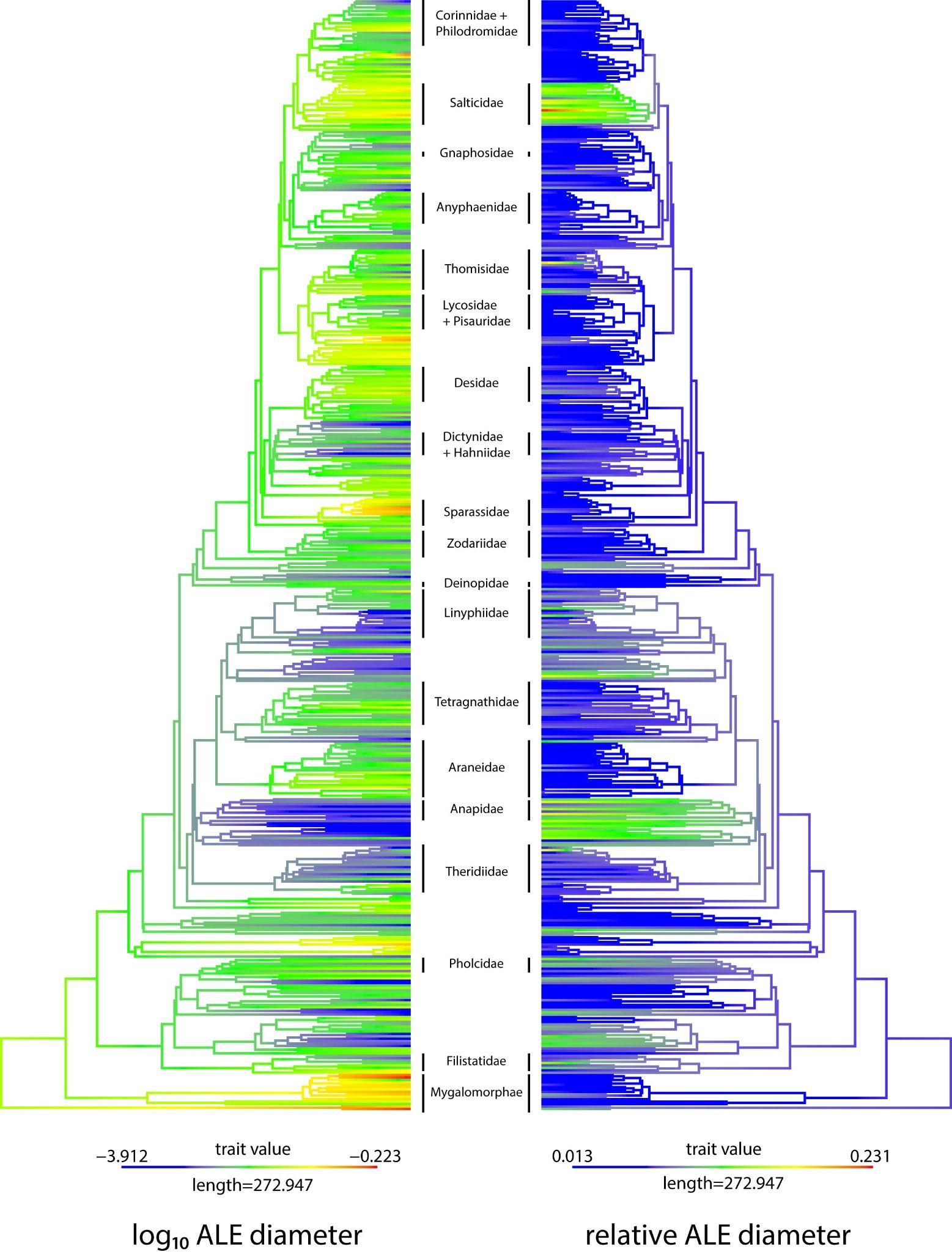

**Figure SF12 Ancestral state reconstruction of log-transformed and relative ALE diameters across spiders.** Curated dataset from Wolff et al. (2022). Position of major families labelled.

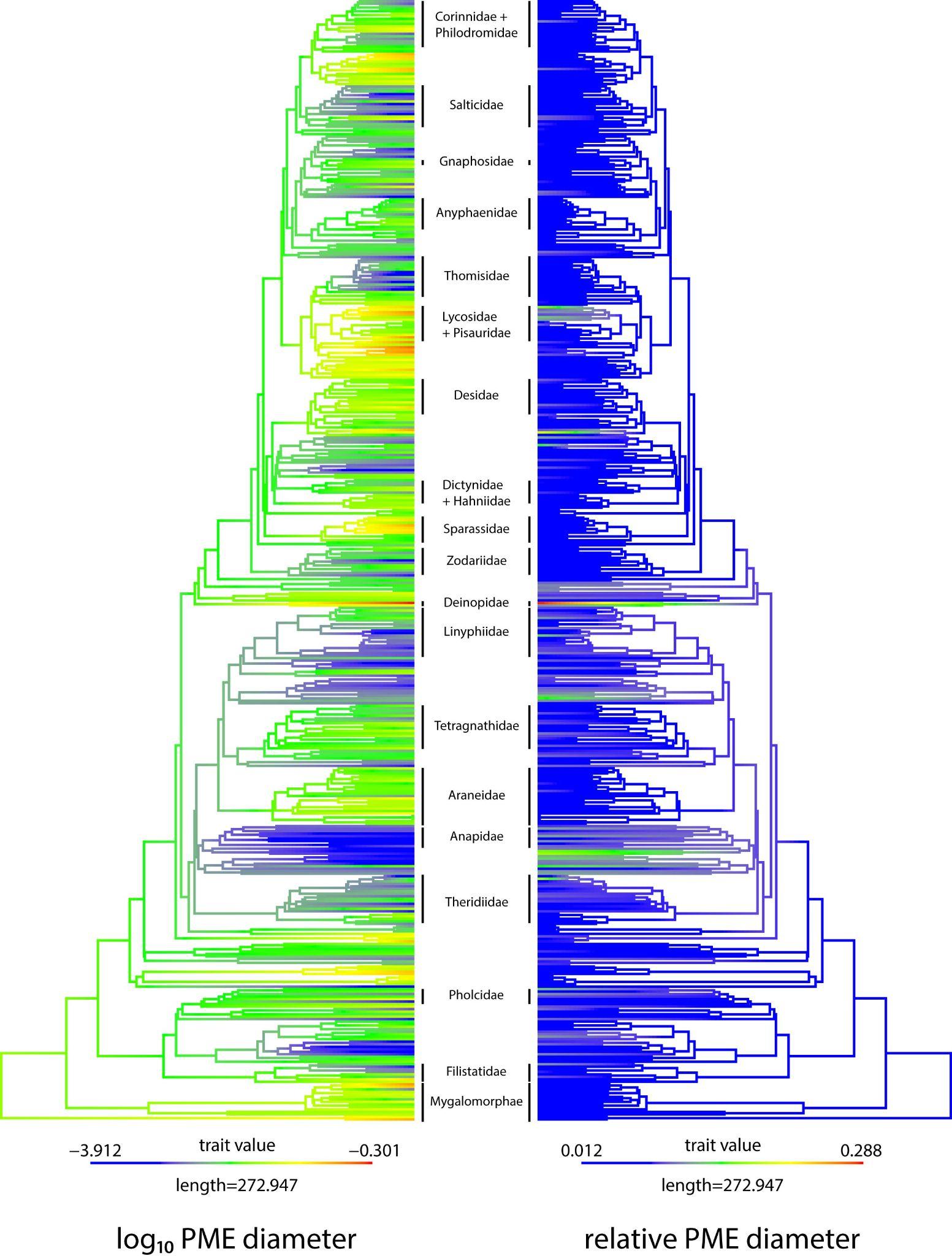

**Figure SF13 Ancestral state reconstruction of log-transformed and relative PME diameters across spiders.** Curated dataset from Wolff et al. (2022). Position of major families labelled.

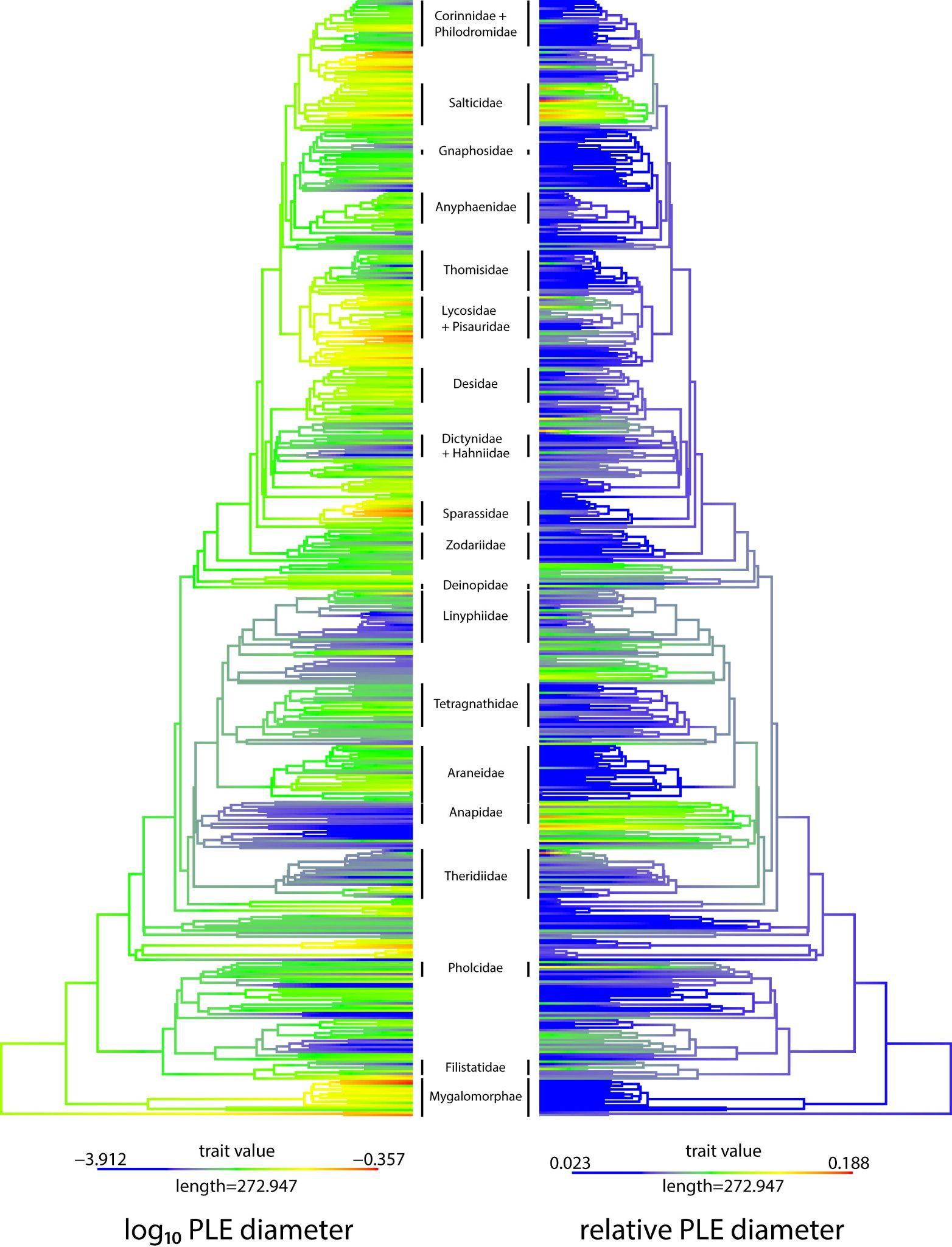

**Figure SF14 Ancestral state reconstruction of log-transformed and relative PLE diameters across spiders.** Curated dataset from Wolff et al. (2022). Position of major families labelled.
