## Supplementary material for "Complex allometric relationships and ecological factors shape the development and evolution of eye size in the modular visual system of spiders": PGLS.Rmd

````
---
title: "WolffData"
author: "LSR"
date: "2023-01-20"
output: html_document
---

```{r setup, include=FALSE}
knitr::opts_chunk$set(echo = TRUE, cache = TRUE)
```

## PGLS analysis of the comparative dataset published by Wolff et al. 2022:

Wolff, J. O., K. Wierucka, G. B. Paterno, J. A. Coddington, G. Hormiga, M. B. J. Kelly, M. E. Herberstein, and M. J. Ramírez. 2022. Stabilized morphological evolution of spiders despite mosaic changes in foraging ecology. Syst. Biol. 0:1–17.

The original dataset contained morphometric data (n=1968 entries) including eye diameters and carapace widths, ecological data (n=829) including hunting guilds, and a time-calibrated maximum-likelihood phylogeny of 828 species. We extracted entries with all eyes, carapace width, and ecological data recorded. Where duplicate entries were present for a given species, we took mean eye diameters and carapace widths. Species missing from the phylogeny were removed. The resulting dataset and phylogeny contained single entries for 474 species. 

Eyes that are absent (e.g. AMEs in Dysdera) were scored diameter of 0 in the original dataset, but these were replaced by NAs for the purposes of these analyses. These data are contained in WolffDataNAs-Dec23.txt. The original data (with 0s) are contained in WolffData-Dec23.txt. The curated phylogeny is contained in WolffRed22Sept.nwk 

The approach to this analysis was based on previous work by Thomas et al. 2020:

Thomas, K. N., D. J. Gower, R. C. Bell, M. K. Fujita, R. K. Schott, and J. W. Streicher. 2020. Eye size and investment in frogs and toads correlate with adult habitat, activity pattern and breeding ecology: Anuran eye size and investment. Proc. R. Soc. B Biol. Sci. 287. ##

```{r}

library(caper)
library(dplyr)
library(phytools)

#read the reduced dataset
WolffDataRedDupsGone <- read.delim("~/WolffData-Dec23.txt") 

#change ecological data to characters
class(WolffDataRedDupsGone$builder) = "character"
class(WolffDataRedDupsGone$orbweb) = "character"
class(WolffDataRedDupsGone$hanging) = "character"
class(WolffDataRedDupsGone$guild) = "character"
class(WolffDataRedDupsGone$var) = "double"
class(WolffDataRedDupsGone$visual.pursuit) = "character"
class(WolffDataRedDupsGone$visual.incl.static) = "character"
class(WolffDataRedDupsGone$active) = "character"

#read the phylogeny
tree<-read.tree("~/WolffRed22Sept.nwk")

#check for data that don't match the tips
datadiff<-setdiff(WolffDataRedDupsGone$terminal, tree$tip.label)
#and remove these from the dataset
data<-WolffDataRedDupsGone[!WolffDataRedDupsGone$terminal %in% datadiff,]

#make comparative object for Caper
df<-comparative.data(phy=tree, data=data, na.omit = FALSE, names.col = terminal, vcv = TRUE, warn.dropped = TRUE)


#with 0 diameters as missing values, affects smallest and variance calculations
WolffDataRedDups0sGone <- read.delim("~/WolffDataNAs-Dec23.txt") 

#change ecological data to characters
class(WolffDataRedDups0sGone$builder) = "character"
class(WolffDataRedDups0sGone$orbweb) = "character"
class(WolffDataRedDups0sGone$hanging) = "character"
class(WolffDataRedDups0sGone$guild) = "character"
class(WolffDataRedDups0sGone$var) = "double"
class(WolffDataRedDupsGone$visual.pursuit) = "character"
class(WolffDataRedDupsGone$visual.incl.static) = "character"
class(WolffDataRedDupsGone$active) = "character"

#and remove the extra tips from the dataset
dataNAs<-WolffDataRedDups0sGone[!WolffDataRedDups0sGone$terminal %in% datadiff,]
#make comparative object for Caper
dfNAs<-comparative.data(phy=tree, data=dataNAs, na.omit = FALSE, names.col = terminal, vcv = TRUE, warn.dropped = TRUE)


```

#PGLS on CW as only predictor (residuals used for comparison of investment between ecological groupings)

Eye pairs analysed one at a time, including only species that have the eye pair

1) AMEs

```{r}


#identify the species lacking AMEs
AME0<-rownames(df$data)[df$data$AME==0]
#remove from tree
AMEtree<-drop.tip(tree, AME0, trim.internal = TRUE)
#remove from data
AMEdata<-data[!data$terminal %in% AME0,]
#make new comparative object
AMEcomp<-comparative.data(phy=AMEtree, data=AMEdata, na.omit = FALSE, names.col = terminal, vcv = TRUE, warn.dropped = TRUE)

#run PGLS 1
pglsAME<-pgls(log(AME) ~ log(CW), AMEcomp, lambda = 'ML')
summary(pglsAME)

plot.pgls(pglsAME)

#are there any phylogenetic outliers?

#extract phylogenetic residuals
presAME<-pglsAME$phyres
#for some reason $residuals takes the rownames with it, but $phyres doesn't?
rownames(presAME)<-rownames(pglsAME$residuals)

#studentise
stpresAME<-presAME/sqrt(var(presAME))[1]
#identify species with studentised residuals >|3|
AMEout<-rownames(stpresAME)[(abs(stpresAME)>3)]

#now these need to be removed and the PGLS run again 

#remove from tree
AMEtree2<-drop.tip(AMEtree, AMEout, trim.internal = TRUE)
#remove from data
AMEdata2<-AMEdata[!AMEdata$terminal %in% AMEout,]
#make new comparative object
AMEcomp2<-comparative.data(phy=AMEtree2, data=AMEdata2, na.omit = FALSE, names.col = terminal, vcv = TRUE, warn.dropped = TRUE)

#run PGLS 2
pglsAME2<-pgls(log(AME) ~ log(CW), AMEcomp2, lambda = 'ML')
summary(pglsAME2)

#are there any phylogenetic outliers?


#extract phylogenetic residuals
presAME<-pglsAME2$phyres
#for some reason $residuals takes the rownames with it, but $phyres doesn't?
rownames(presAME)<-rownames(pglsAME2$residuals)

#studentise
stpresAME<-presAME/sqrt(var(presAME))[1]
#identify species with studentised residuals >|3|
AMEout2<-rownames(stpresAME)[(abs(stpresAME)>3)]

#now these need to be removed and the PGLS run again 

#remove from tree
AMEtree3<-drop.tip(AMEtree2, AMEout2, trim.internal = TRUE)
#remove from data
AMEdata3<-AMEdata2[!AMEdata2$terminal %in% AMEout2,]
#make new comparative object
AMEcomp3<-comparative.data(phy=AMEtree3, data=AMEdata3, na.omit = FALSE, names.col = terminal, vcv = TRUE, warn.dropped = TRUE)

#run PGLS 3
pglsAME3<-pgls(log(AME) ~ log(CW), AMEcomp3, lambda = 'ML')
summary(pglsAME3)

#are there any phylogenetic outliers?

#extract phylogenetic residuals
presAME<-pglsAME3$phyres
#for some reason $residuals takes the rownames with it, but $phyres doesn't?
rownames(presAME)<-rownames(pglsAME3$residuals)

#studentise
stpresAME<-presAME/sqrt(var(presAME))[1]
#identify species with studentised residuals >|3|
AMEout3<-rownames(stpresAME)[(abs(stpresAME)>3)]

#now these need to be removed and the PGLS run again 

#remove from tree
AMEtree4<-drop.tip(AMEtree3, AMEout3, trim.internal = TRUE)
#remove from data
AMEdata4<-AMEdata3[!AMEdata3$terminal %in% AMEout3,]
#make new comparative object
AMEcomp4<-comparative.data(phy=AMEtree4, data=AMEdata4, na.omit = FALSE, names.col = terminal, vcv = TRUE, warn.dropped = TRUE)


#run PGLS 4
pglsAME4<-pgls(log(AME) ~ log(CW), AMEcomp4, lambda = 'ML')
summary(pglsAME4)

#are there any phylogenetic outliers?

#extract phylogenetic residuals
presAME<-pglsAME4$phyres
#for some reason $residuals takes the rownames with it, but $phyres doesn't?
rownames(presAME)<-rownames(pglsAME4$residuals)

#studentise
stpresAME<-presAME/sqrt(var(presAME))[1]
#identify species with studentised residuals >|3|
AMEout4<-rownames(stpresAME)[(abs(stpresAME)>3)]

#now these need to be removed and the PGLS run again 

#remove from tree
AMEtree5<-drop.tip(AMEtree4, AMEout4, trim.internal = TRUE)
#remove from data
AMEdata5<-AMEdata4[!AMEdata4$terminal %in% AMEout4,]
#make new comparative object
AMEcomp5<-comparative.data(phy=AMEtree5, data=AMEdata5, na.omit = FALSE, names.col = terminal, vcv = TRUE, warn.dropped = TRUE)

#run PGLS 5
pglsAME5<-pgls(log(AME) ~ log(CW), AMEcomp5, lambda = 'ML')
summary(pglsAME5)

#are there any phylogenetic outliers?

#extract phylogenetic residuals
presAME<-pglsAME5$phyres
#for some reason $residuals takes the rownames with it, but $phyres doesn't?
rownames(presAME)<-rownames(pglsAME5$residuals)

#studentise
stpresAME<-presAME/sqrt(var(presAME))[1]
AMEout5<-rownames(stpresAME)[(abs(stpresAME)>3)]
AMEout5

#once that's done, we can use $residuals to look at the actual investment

#extract residuals (not phylogenetic residuals!)
resAME<-pglsAME5$residuals
#Studentise
stresAME <- resAME/sqrt(var(resAME))[1]
#are there any outliers? i.e. studentised residuals >|3|?
rownames(stresAME)[(abs(stresAME)>3)]
#sense check - for the AMEs this should be a bunch of salticids

#add the stres as a new column to the original data

#make sure they're in the right order first!
stresAME <- stresAME[order(row.names(stresAME)),]
spider.AME.res <- cbind (AMEdata5, stresAME = stresAME)

spider.AME.res

```

ALEs

```{r}

#identify the species lacking AMEs
ALE0<-rownames(df$data)[df$data$ALE==0]

#remove from tree
ALEtree<-drop.tip(tree, ALE0, trim.internal = TRUE)
#remove from data
ALEdata<-data[!data$terminal %in% ALE0,]
#make new comparative object
ALEcomp<-comparative.data(phy=ALEtree, data=ALEdata, na.omit = FALSE, names.col = terminal, vcv = TRUE, warn.dropped = TRUE)

#run PGLS 1
pglsALE<-pgls(log(ALE) ~ log(CW), ALEcomp, lambda = 'ML')
summary(pglsALE)

plot.pgls(pglsALE)

#are there any phylogenetic outliers?

#extract phylogenetic residuals
presALE<-pglsALE$phyres
#for some reason $residuals takes the rownames with it, but $phyres doesn't?
rownames(presALE)<-rownames(pglsALE$residuals)

#studentise
stpresALE<-presALE/sqrt(var(presALE))[1]
#identify species with studentised residuals >|3|
ALEout<-rownames(stpresALE)[(abs(stpresALE)>3)]

#now these need to be removed and the PGLS run again 

#remove from tree
ALEtree2<-drop.tip(tree, ALEout, trim.internal = TRUE)
#remove from data
ALEdata2<-ALEdata[!ALEdata$terminal %in% ALEout,]
#make new comparative object
ALEcomp2<-comparative.data(phy=ALEtree2, data=ALEdata2, na.omit = FALSE, names.col = terminal, vcv = TRUE, warn.dropped = TRUE)

#run PGLS 2
pglsALE2<-pgls(log(ALE) ~ log(CW), ALEcomp2, lambda = 'ML')
summary(pglsALE2)

#are there any phylogenetic outliers?

#extract phylogenetic residuals
presALE<-pglsALE2$phyres
#for some reason $residuals takes the rownames with it, but $phyres doesn't?
rownames(presALE)<-rownames(pglsALE2$residuals)

#studentise
stpresALE<-presALE/sqrt(var(presALE))[1]
#identify species with studentised residuals >|3|
ALEout2<-rownames(stpresALE)[(abs(stpresALE)>3)]

#now these need to be removed and the PGLS run again 

#remove from tree
ALEtree3<-drop.tip(ALEtree2, ALEout2, trim.internal = TRUE)
#remove from data
ALEdata3<-ALEdata2[!ALEdata2$terminal %in% ALEout2,]
#make new comparative object
ALEcomp3<-comparative.data(phy=ALEtree3, data=ALEdata3, na.omit = FALSE, names.col = terminal, vcv = TRUE, warn.dropped = TRUE)

#run PGLS 3
pglsALE3<-pgls(log(ALE) ~ log(CW), ALEcomp3, lambda = 'ML')
summary(pglsALE3)

#are there any phylogenetic outliers?

#extract phylogenetic residuals
presALE<-pglsALE3$phyres
#for some reason $residuals takes the rownames with it, but $phyres doesn't?
rownames(presALE)<-rownames(pglsALE3$residuals)

#studentise
stpresALE<-presALE/sqrt(var(presALE))[1]
#identify species with studentised residuals >|3|
ALEout3<-rownames(stpresALE)[(abs(stpresALE)>3)]

#now these need to be removed and the PGLS run again 

#remove from tree
ALEtree4<-drop.tip(ALEtree3, ALEout3, trim.internal = TRUE)
#remove from data
ALEdata4<-ALEdata3[!ALEdata3$terminal %in% ALEout3,]
#make new comparative object
ALEcomp4<-comparative.data(phy=ALEtree4, data=ALEdata4, na.omit = FALSE, names.col = terminal, vcv = TRUE, warn.dropped = TRUE)

#run PGLS 4
pglsALE4<-pgls(log(ALE) ~ log(CW), ALEcomp4, lambda = 'ML')
summary(pglsALE4)

#are there any phylogenetic outliers?

#extract phylogenetic residuals
presALE<-pglsALE4$phyres
#for some reason $residuals takes the rownames with it, but $phyres doesn't?
rownames(presALE)<-rownames(pglsALE4$residuals)

#studentise
stpresALE<-presALE/sqrt(var(presALE))[1]
#identify species with studentised residuals >|3|
ALEout4<-rownames(stpresALE)[(abs(stpresALE)>3)]

#now these need to be removed and the PGLS run again 

#remove from tree
ALEtree5<-drop.tip(ALEtree4, ALEout4, trim.internal = TRUE)
#remove from data
ALEdata5<-ALEdata4[!ALEdata4$terminal %in% ALEout4,]
#make new comparative object
ALEcomp5<-comparative.data(phy=ALEtree5, data=ALEdata5, na.omit = FALSE, names.col = terminal, vcv = TRUE, warn.dropped = TRUE)

#run PGLS 5
pglsALE5<-pgls(log(ALE) ~ log(CW), ALEcomp5, lambda = 'ML')
summary(pglsALE5)

#are there any phylogenetic outliers?

#extract phylogenetic residuals
presALE<-pglsALE5$phyres
#for some reason $residuals takes the rownames with it, but $phyres doesn't?
rownames(presALE)<-rownames(pglsALE5$residuals)

#studentise
stpresALE<-presALE/sqrt(var(presALE))[1]
#identify species with studentised residuals >|3|
ALEout5<-rownames(stpresALE)[(abs(stpresALE)>3)]

#once that's done, we can use $residuals to look at the actual investment

#extract residuals (not phylogenetic residuals!)
resALE<-pglsALE5$residuals
#Studentise
stresALE <- resALE/sqrt(var(resALE))[1]
#identify species with studentised residuals >|3|
rownames(stresALE)[(abs(stresALE)>3)]


#add the stres as a new column to the original data

#make sure they're in the right order first!
stresALE <- stresALE[order(row.names(stresALE)),]
spider.ALE.res <- cbind (ALEdata5, stresALE = stresALE)

spider.ALE.res

```

PMEs

```{r}

#identify the species lacking PMEs
PME0<-rownames(df$data)[df$data$PME==0]

#remove from tree
PMEtree<-drop.tip(tree, PME0, trim.internal = TRUE)
#remove from data
PMEdata<-data[!data$terminal %in% PME0,]
#make new comparative object
PMEcomp<-comparative.data(phy=PMEtree, data=PMEdata, na.omit = FALSE, names.col = terminal, vcv = TRUE, warn.dropped = TRUE)

#run PGLS 1
pglsPME<-pgls(log(PME) ~ log(CW), PMEcomp, lambda = 'ML')
summary(pglsPME)

plot.pgls(pglsPME)

#are there any phylogenetic outliers?

#extract phylogenetic residuals
presPME<-pglsPME$phyres
#for some reason $residuals takes the rownames with it, but $phyres doesn't?
rownames(presPME)<-rownames(pglsPME$residuals)

#studentise
stpresPME<-presPME/sqrt(var(presPME))[1]
#identify species with studentised residuals >|3|
PMEout<-rownames(stpresPME)[(abs(stpresPME)>3)]

#now these need to be removed and the PGLS run again 

#remove from tree
PMEtree2<-drop.tip(PMEtree, PMEout, trim.internal = TRUE)
#remove from data
PMEdata2<-PMEdata[!PMEdata$terminal %in% PMEout,]
#make new comparative object
PMEcomp2<-comparative.data(phy=PMEtree2, data=PMEdata2, na.omit = FALSE, names.col = terminal, vcv = TRUE, warn.dropped = TRUE)

#run PGLS 2
pglsPME2<-pgls(log(PME) ~ log(CW), PMEcomp2, lambda = 'ML')
summary(pglsPME2)

#are there any phylogenetic outliers?

#extract phylogenetic residuals
presPME<-pglsPME2$phyres
#for some reason $residuals takes the rownames with it, but $phyres doesn't?
rownames(presPME)<-rownames(pglsPME2$residuals)

#studentise
stpresPME<-presPME/sqrt(var(presPME))[1]
#identify species with studentised residuals >|3|
PMEout2<-rownames(stpresPME)[(abs(stpresPME)>3)]

#now these need to be removed and the PGLS run again 

#remove from tree
PMEtree3<-drop.tip(PMEtree2, PMEout2, trim.internal = TRUE)
#remove from data
PMEdata3<-PMEdata2[!PMEdata2$terminal %in% PMEout2,]
#make new comparative object
PMEcomp3<-comparative.data(phy=PMEtree3, data=PMEdata3, na.omit = FALSE, names.col = terminal, vcv = TRUE, warn.dropped = TRUE)

#run PGLS 3
pglsPME3<-pgls(log(PME) ~ log(CW), PMEcomp3, lambda = 'ML')
summary(pglsPME3)

#are there any phylogenetic outliers?

#extract phylogenetic residuals
presPME<-pglsPME3$phyres
#for some reason $residuals takes the rownames with it, but $phyres doesn't?
rownames(presPME)<-rownames(pglsPME3$residuals)

#studentise
stpresPME<-presPME/sqrt(var(presPME))[1]
#identify species with studentised residuals >|3|
PMEout3<-rownames(stpresPME)[(abs(stpresPME)>3)]

#now these need to be removed and the PGLS run again 

#remove from tree
PMEtree4<-drop.tip(PMEtree3, PMEout3, trim.internal = TRUE)
#remove from data
PMEdata4<-PMEdata3[!PMEdata3$terminal %in% PMEout3,]
#make new comparative object
PMEcomp4<-comparative.data(phy=PMEtree4, data=PMEdata4, na.omit = FALSE, names.col = terminal, vcv = TRUE, warn.dropped = TRUE)

#run PGLS 4
pglsPME4<-pgls(log(PME) ~ log(CW), PMEcomp4, lambda = 'ML')
summary(pglsPME4)

#are there any phylogenetic outliers?

#extract phylogenetic residuals
presPME<-pglsPME4$phyres
#for some reason $residuals takes the rownames with it, but $phyres doesn't?
rownames(presPME)<-rownames(pglsPME4$residuals)

#studentise
stpresPME<-presPME/sqrt(var(presPME))[1]
#identify species with studentised residuals >|3|
PMEout4<-rownames(stpresPME)[(abs(stpresPME)>3)]

#now these need to be removed and the PGLS run again 

#remove from tree
PMEtree5<-drop.tip(PMEtree4, PMEout4, trim.internal = TRUE)
#remove from data
PMEdata5<-PMEdata4[!PMEdata4$terminal %in% PMEout4,]
#make new comparative object
PMEcomp5<-comparative.data(phy=PMEtree5, data=PMEdata5, na.omit = FALSE, names.col = terminal, vcv = TRUE, warn.dropped = TRUE)

#run PGLS 5
pglsPME5<-pgls(log(PME) ~ log(CW), PMEcomp5, lambda = 'ML')
summary(pglsPME5)

#are there any phylogenetic outliers?

#extract phylogenetic residuals
presPME<-pglsPME5$phyres
#for some reason $residuals takes the rownames with it, but $phyres doesn't?
rownames(presPME)<-rownames(pglsPME5$residuals)

#studentise
stpresPME<-presPME/sqrt(var(presPME))[1]
#identify species with studentised residuals >|3|
PMEout5<-rownames(stpresPME)[(abs(stpresPME)>3)]

#once that's done, we can use $residuals to look at the actual investment

#extract residuals (not phylogenetic residuals!)
resPME<-pglsPME5$residuals
#Studentise
stresPME <- resPME/sqrt(var(resPME))[1]
#identify species with studentised residuals >|3|
rownames(stresPME)[(abs(stresPME)>3)]


#add the stres as a new column to the original data

#make sure they're in the right order first!
stresPME <- stresPME[order(row.names(stresPME)),]
spider.PME.res <- cbind (PMEdata5, stpresPME = stpresPME)

spider.PME.res

```

PLEs

```{r}

#identify the species lacking PLEs
PLE0<-rownames(df$data)[df$data$PLE==0]

#remove from tree
PLEtree<-drop.tip(tree, PLE0, trim.internal = TRUE)
#remove from data
PLEdata<-data[!data$terminal %in% PLE0,]
#make new comparative object
PLEcomp<-comparative.data(phy=PLEtree, data=PLEdata, na.omit = FALSE, names.col = terminal, vcv = TRUE, warn.dropped = TRUE)

#run PGLS 1
pglsPLE<-pgls(log(PLE) ~ log(CW), PLEcomp, lambda = 'ML')
summary(pglsPLE)

plot.pgls(pglsPLE)

#are there any phylogenetic outliers?

#extract phylogenetic residuals
presPLE<-pglsPLE$phyres
#for some reason $residuals takes the rownames with it, but $phyres doesn't?
rownames(presPLE)<-rownames(pglsPLE$residuals)

#studentise
stpresPLE<-presPLE/sqrt(var(presPLE))[1]
#identify species with studentised residuals >|3|
PLEout<-rownames(stpresPLE)[(abs(stpresPLE)>3)]

#now these need to be removed and the PGLS run again 

#remove from tree
PLEtree2<-drop.tip(PLEtree, PLEout, trim.internal = TRUE)
#remove from data
PLEdata2<-PLEdata[!PLEdata$terminal %in% PLEout,]
#make new comparative object
PLEcomp2<-comparative.data(phy=PLEtree2, data=PLEdata2, na.omit = FALSE, names.col = terminal, vcv = TRUE, warn.dropped = TRUE)

#run PGLS 2
pglsPLE2<-pgls(log(PLE) ~ log(CW), PLEcomp2, lambda = 'ML')
summary(pglsPLE2)

#are there any phylogenetic outliers?

#extract phylogenetic residuals
presPLE<-pglsPLE2$phyres
#for some reason $residuals takes the rownames with it, but $phyres doesn't?
rownames(presPLE)<-rownames(pglsPLE2$residuals)

#studentise
stpresPLE<-presPLE/sqrt(var(presPLE))[1]
#identify species with studentised residuals >|3|
PLEout2<-rownames(stpresPLE)[(abs(stpresPLE)>3)]

#now these need to be removed and the PGLS run again 

#remove from tree
PLEtree3<-drop.tip(PLEtree2, PLEout2, trim.internal = TRUE)
#remove from data
PLEdata3<-PLEdata2[!PLEdata2$terminal %in% PLEout2,]
#make new comparative object
PLEcomp3<-comparative.data(phy=PLEtree3, data=PLEdata3, na.omit = FALSE, names.col = terminal, vcv = TRUE, warn.dropped = TRUE)

#run PGLS 3
pglsPLE3<-pgls(log(PLE) ~ log(CW), PLEcomp3, lambda = 'ML')
summary(pglsPLE3)

#are there any phylogenetic outliers?

#extract phylogenetic residuals
presPLE<-pglsPLE3$phyres
#for some reason $residuals takes the rownames with it, but $phyres doesn't?
rownames(presPLE)<-rownames(pglsPLE3$residuals)

#studentise
stpresPLE<-presPLE/sqrt(var(presPLE))[1]
#identify species with studentised residuals >|3|
PLEout3<-rownames(stpresPLE)[(abs(stpresPLE)>3)]

#now these need to be removed and the PGLS run again 

#remove from tree
PLEtree4<-drop.tip(PLEtree3, PLEout3, trim.internal = TRUE)
#remove from data
PLEdata4<-PLEdata3[!PLEdata3$terminal %in% PLEout3,]
#make new comparative object
PLEcomp4<-comparative.data(phy=PLEtree4, data=PLEdata4, na.omit = FALSE, names.col = terminal, vcv = TRUE, warn.dropped = TRUE)

#run PGLS 4
pglsPLE4<-pgls(log(PLE) ~ log(CW), PLEcomp4, lambda = 'ML')
summary(pglsPLE4)

#are there any phylogenetic outliers?

#extract phylogenetic residuals
presPLE<-pglsPLE4$phyres
#for some reason $residuals takes the rownames with it, but $phyres doesn't?
rownames(presPLE)<-rownames(pglsPLE4$residuals)

#studentise
stpresPLE<-presPLE/sqrt(var(presPLE))[1]
#identify species with studentised residuals >|3|
PLEout4<-rownames(stpresPLE)[(abs(stpresPLE)>3)]

#now these need to be removed and the PGLS run again 

#remove from tree
PLEtree5<-drop.tip(PLEtree4, PLEout4, trim.internal = TRUE)
#remove from data
PLEdata5<-PLEdata4[!PLEdata4$terminal %in% PLEout4,]
#make new comparative object
PLEcomp5<-comparative.data(phy=PLEtree5, data=PLEdata5, na.omit = FALSE, names.col = terminal, vcv = TRUE, warn.dropped = TRUE)

#run PGLS 5
pglsPLE5<-pgls(log(PLE) ~ log(CW), PLEcomp5, lambda = 'ML')
summary(pglsPLE5)

#are there any phylogenetic outliers?

#extract phylogenetic residuals
presPLE<-pglsPLE5$phyres
#for some reason $residuals takes the rownames with it, but $phyres doesn't?
rownames(presPLE)<-rownames(pglsPLE5$residuals)

#studentise
stpresPLE<-presPLE/sqrt(var(presPLE))[1]
#identify species with studentised residuals >|3|
PLEout5<-rownames(stpresPLE)[(abs(stpresPLE)>3)]

#once that's done, we can use $residuals to look at the actual investment

#extract residuals (not phylogenetic residuals!)
resPLE<-pglsPLE5$residuals
#studentise
stresPLE <- resPLE/sqrt(var(resPLE))[1]
#identify species with studentised residuals >|3|
rownames(stresPLE)[(abs(stresPLE)>3)]

#add the stres as a new column to the original data

#make sure they're in the right order first!
stresPLE <- stresPLE[order(row.names(stresPLE)),]
spider.PLE.res <- cbind (PLEdata5, stresPLE = stresPLE)

spider.PLE.res

```

Largest - this uses the whole comparative dataset

```{r}

pglsLargest<-pgls(log(Largest) ~ log(CW), df, lambda = 'ML')
summary(pglsLargest)

#are there any phylogenetic outliers?

#extract phylogenetic residuals
presLargest<-pglsLargest$phyres
#for some reason $residuals takes the rownames with it, but $phyres doesn't?
rownames(presLargest)<-rownames(pglsLargest$residuals)

#studentise
stpresLargest<-presLargest/sqrt(var(presLargest))[1]
#identify species with studentised residuals >|3|
Largestout<-rownames(stpresLargest)[(abs(stpresLargest)>3)]

#now these need to be removed and the PGLS run again 

#remove from tree
Largesttree2<-drop.tip(tree, Largestout, trim.internal = TRUE)
#remove from data
Largestdata2<-data[!data$terminal %in% Largestout,]
#make new comparative object
Largestcomp2<-comparative.data(phy=Largesttree2, data=Largestdata2, na.omit = FALSE, names.col = terminal, vcv = TRUE, warn.dropped = TRUE)

#run PGLS 2
pglsLargest2<-pgls(log(Largest) ~ log(CW), Largestcomp2, lambda = 'ML')
summary(pglsLargest2)

#are there any phylogenetic outliers?

#extract phylogenetic residuals
presLargest<-pglsLargest2$phyres
#for some reason $residuals takes the rownames with it, but $phyres doesn't?
rownames(presLargest)<-rownames(pglsLargest2$residuals)

#studentise
stpresLargest<-presLargest/sqrt(var(presLargest))[1]
#identify species with studentised residuals >|3|
Largestout2<-rownames(stpresLargest)[(abs(stpresLargest)>3)]

#now these need to be removed and the PGLS run again 

#remove from tree
Largesttree3<-drop.tip(Largesttree2, Largestout2, trim.internal = TRUE)
#remove from data
Largestdata3<-Largestdata2[!Largestdata2$terminal %in% Largestout2,]
#make new comparative object
Largestcomp3<-comparative.data(phy=Largesttree3, data=Largestdata3, na.omit = FALSE, names.col = terminal, vcv = TRUE, warn.dropped = TRUE)

#run PGLS 3
pglsLargest3<-pgls(log(Largest) ~ log(CW), Largestcomp3, lambda = 'ML')
summary(pglsLargest3)

#are there any phylogenetic outliers?

#extract phylogenetic residuals
presLargest<-pglsLargest3$phyres
#for some reason $residuals takes the rownames with it, but $phyres doesn't?
rownames(presLargest)<-rownames(pglsLargest3$residuals)

#studentise
stpresLargest<-presLargest/sqrt(var(presLargest))[1]
#identify species with studentised residuals >|3|
Largestout3<-rownames(stpresLargest)[(abs(stpresLargest)>3)]

#now these need to be removed and the PGLS run again 

#remove from tree
Largesttree4<-drop.tip(Largesttree3, Largestout3, trim.internal = TRUE)
#remove from data
Largestdata4<-Largestdata3[!Largestdata3$terminal %in% Largestout3,]
#make new comparative object
Largestcomp4<-comparative.data(phy=Largesttree4, data=Largestdata4, na.omit = FALSE, names.col = terminal, vcv = TRUE, warn.dropped = TRUE)

#run PGLS 4
pglsLargest4<-pgls(log(Largest) ~ log(CW), Largestcomp4, lambda = 'ML')
summary(pglsLargest4)

#are there any phylogenetic outliers?

#extract phylogenetic residuals
presLargest<-pglsLargest4$phyres
#for some reason $residuals takes the rownames with it, but $phyres doesn't?
rownames(presLargest)<-rownames(pglsLargest4$residuals)

#studentise
stpresLargest<-presLargest/sqrt(var(presLargest))[1]
#identify species with studentised residuals >|3|
Largestout4<-rownames(stpresLargest)[(abs(stpresLargest)>3)]

#now these need to be removed and the PGLS run again 

#remove from tree
Largesttree5<-drop.tip(Largesttree4, Largestout4, trim.internal = TRUE)
#remove from data
Largestdata5<-Largestdata4[!Largestdata4$terminal %in% Largestout4,]
#make new comparative object
Largestcomp5<-comparative.data(phy=Largesttree5, data=Largestdata5, na.omit = FALSE, names.col = terminal, vcv = TRUE, warn.dropped = TRUE)

#run PGLS 5
pglsLargest5<-pgls(log(Largest) ~ log(CW), Largestcomp5, lambda = 'ML')
summary(pglsLargest5)

#are there any phylogenetic outliers?

#extract phylogenetic residuals
presLargest<-pglsLargest5$phyres
#for some reason $residuals takes the rownames with it, but $phyres doesn't?
rownames(presLargest)<-rownames(pglsLargest5$residuals)

#studentise
stpresLargest<-presLargest/sqrt(var(presLargest))[1]
#identify species with studentised residuals >|3|
Largestout5<-rownames(stpresLargest)[(abs(stpresLargest)>3)]

#once that's done, we can use $residuals to look at the actual investment

#extract residuals (not phylogenetic residuals!)
resLargest<-pglsLargest5$residuals
#studentise
stresLargest <- resLargest/sqrt(var(resLargest))[1]
#identify species with studentised residuals >|3|
rownames(stresLargest)[(abs(stresLargest)>3)]

#add the stres as a new column to the original data

#make sure they're in the right order first!
stresLargest <- stresLargest[order(row.names(stresLargest)),]
spider.Largest.res <- cbind (Largestdata5, stresLargest = stresLargest)

spider.Largest.res

```

Smallest

```{r}

#identify the species lacking any eye pair
Small0<-rownames(df$data)[df$data$Smallest==0]

#remove from tree
Smallesttree<-drop.tip(tree, Small0, trim.internal = TRUE)
#remove from data
Smallestdata<-data[!data$terminal %in% Small0,]
#make new comparative object
Smallestcomp<-comparative.data(phy=Smallesttree, data=Smallestdata, na.omit = FALSE, names.col = terminal, vcv = TRUE, warn.dropped = TRUE)

#run PGLS 1
pglsSmallest<-pgls(log(Smallest) ~ log(CW), Smallestcomp, lambda = 'ML')
summary(pglsSmallest)

plot.pgls(pglsSmallest)

#are there any phylogenetic outliers?

#extract phylogenetic residuals
presSmallest<-pglsSmallest$phyres
#for some reason $residuals takes the rownames with it, but $phyres doesn't?
rownames(presSmallest)<-rownames(pglsSmallest$residuals)

#studentise
stpresSmallest<-presSmallest/sqrt(var(presSmallest))[1]
#identify species with studentised residuals >|3|
Smallestout<-rownames(stpresSmallest)[(abs(stpresSmallest)>3)]

#now these need to be removed and the PGLS run again 

#remove from tree
Smallesttree2<-drop.tip(Smallesttree, Smallestout, trim.internal = TRUE)
#remove from data
Smallestdata2<-Smallestdata[!Smallestdata$terminal %in% Smallestout,]
#make new comparative object
Smallestcomp2<-comparative.data(phy=Smallesttree2, data=Smallestdata2, na.omit = FALSE, names.col = terminal, vcv = TRUE, warn.dropped = TRUE)

#run PGLS 2
pglsSmallest2<-pgls(log(Smallest) ~ log(CW), Smallestcomp2, lambda = 'ML')
summary(pglsSmallest2)

#are there any phylogenetic outliers?

#extract phylogenetic residuals
presSmallest<-pglsSmallest2$phyres
#for some reason $residuals takes the rownames with it, but $phyres doesn't?
rownames(presSmallest)<-rownames(pglsSmallest2$residuals)

#studentise
stpresSmallest<-presSmallest/sqrt(var(presSmallest))[1]
#identify species with studentised residuals >|3|
Smallestout2<-rownames(stpresSmallest)[(abs(stpresSmallest)>3)]

#now these need to be removed and the PGLS run again 

#remove from tree
Smallesttree3<-drop.tip(Smallesttree2, Smallestout2, trim.internal = TRUE)
#remove from data
Smallestdata3<-Smallestdata2[!Smallestdata2$terminal %in% Smallestout2,]
#make new comparative object
Smallestcomp3<-comparative.data(phy=Smallesttree3, data=Smallestdata3, na.omit = FALSE, names.col = terminal, vcv = TRUE, warn.dropped = TRUE)

#run PGLS 3
pglsSmallest3<-pgls(log(Smallest) ~ log(CW), Smallestcomp3, lambda = 'ML')
summary(pglsSmallest3)

#are there any phylogenetic outliers?

#extract phylogenetic residuals
presSmallest<-pglsSmallest3$phyres
#for some reason $residuals takes the rownames with it, but $phyres doesn't?
rownames(presSmallest)<-rownames(pglsSmallest3$residuals)

#studentise
stpresSmallest<-presSmallest/sqrt(var(presSmallest))[1]
#identify species with studentised residuals >|3|
Smallestout3<-rownames(stpresSmallest)[(abs(stpresSmallest)>3)]

#now these need to be removed and the PGLS run again 

#remove from tree
Smallesttree4<-drop.tip(Smallesttree3, Smallestout3, trim.internal = TRUE)
#remove from data
Smallestdata4<-Smallestdata3[!Smallestdata3$terminal %in% Smallestout3,]
#make new comparative object
Smallestcomp4<-comparative.data(phy=Smallesttree4, data=Smallestdata4, na.omit = FALSE, names.col = terminal, vcv = TRUE, warn.dropped = TRUE)

#run PGLS 4
pglsSmallest4<-pgls(log(Smallest) ~ log(CW), Smallestcomp4, lambda = 'ML')
summary(pglsSmallest4)

#are there any phylogenetic outliers?

#extract phylogenetic residuals
presSmallest<-pglsSmallest4$phyres
#for some reason $residuals takes the rownames with it, but $phyres doesn't?
rownames(presSmallest)<-rownames(pglsSmallest4$residuals)

#studentise
stpresSmallest<-presSmallest/sqrt(var(presSmallest))[1]
#identify species with studentised residuals >|3|
Smallestout4<-rownames(stpresSmallest)[(abs(stpresSmallest)>3)]

#now these need to be removed and the PGLS run again 

#remove from tree
Smallesttree5<-drop.tip(Smallesttree4, Smallestout4, trim.internal = TRUE)
#remove from data
Smallestdata5<-Smallestdata4[!Smallestdata4$terminal %in% Smallestout4,]
#make new comparative object
Smallestcomp5<-comparative.data(phy=Smallesttree5, data=Smallestdata5, na.omit = FALSE, names.col = terminal, vcv = TRUE, warn.dropped = TRUE)

#run PGLS 5
pglsSmallest5<-pgls(log(Smallest) ~ log(CW), Smallestcomp5, lambda = 'ML')
summary(pglsSmallest5)

#are there any phylogenetic outliers?

#extract phylogenetic residuals
presSmallest<-pglsSmallest5$phyres
#for some reason $residuals takes the rownames with it, but $phyres doesn't?
rownames(presSmallest)<-rownames(pglsSmallest5$residuals)

#residuals
stpresSmallest<-presSmallest/sqrt(var(presSmallest))[1]
#identify species with studentised residuals >|3|
Smallestout5<-rownames(stpresSmallest)[(abs(stpresSmallest)>3)]

#once that's done, we can use $residuals to look at the actual investment

#extract residuals (not phylogenetic residuals!)
resSmallest<-pglsSmallest5$residuals
#studentise
stresSmallest <- resSmallest/sqrt(var(resSmallest))[1]
#identify species with studentised residuals >|3|
rownames(stresSmallest)[(abs(stresSmallest)>3)]

#add the stres as a new column to the original data

#make sure they're in the right order first!
stresSmallest <- stresSmallest[order(row.names(stresSmallest)),]
spider.Smallest.res <- cbind (Smallestdata5, stresSmallest = stresSmallest)

spider.Smallest.res


```

And using NAs dataset (so, smallest of the existing eye pairs for ALL species)

```{r}

#run  PGLS 1
pglsSmallestNAs<-pgls(log(Smallest) ~ log(CW), dfNAs, lambda = 'ML')
summary(pglsSmallestNAs)
#fixed from before via na.omit = FALSE when creating comparative object

plot.pgls(pglsSmallestNAs)

#are there any phylogenetic outliers?

#extract phylogenetic residuals
presSmallestNAs<-pglsSmallestNAs$phyres
#for some reason $residuals takes the rownames with it, but $phyres doesn't?
rownames(presSmallestNAs)<-rownames(pglsSmallestNAs$residuals)

#studentise
stpresSmallestNAs<-presSmallestNAs/sqrt(var(presSmallestNAs))[1]
#identify species with studentised residuals >|3|
SmallestNAsout<-rownames(stpresSmallestNAs)[(abs(stpresSmallestNAs)>3)]

#now these need to be removed and the PGLS run again 

#remove from tree
SmallestNAstree2<-drop.tip(SmallestNAstree, SmallestNAsout, trim.internal = TRUE)
#remove from data
SmallestNAsdata2<-SmallestNAsdata[!SmallestNAsdata$terminal %in% SmallestNAsout,]
#make new comparative object
SmallestNAscomp2<-comparative.data(phy=SmallestNAstree2, data=SmallestNAsdata2, na.omit = FALSE, names.col = terminal, vcv = TRUE, warn.dropped = TRUE)

#run PGLS 2
pglsSmallestNAs2<-pgls(log(SmallestNAs) ~ log(CW), SmallestNAscomp2, lambda = 'ML')
summary(pglsSmallestNAs2)

#are there any phylogenetic outliers?

#extract phylogenetic residuals
presSmallestNAs<-pglsSmallestNAs2$phyres
#for some reason $residuals takes the rownames with it, but $phyres doesn't?
rownames(presSmallestNAs)<-rownames(pglsSmallestNAs2$residuals)

#studentise
stpresSmallestNAs<-presSmallestNAs/sqrt(var(presSmallestNAs))[1]
#identify species with studentised residuals >|3|
SmallestNAsout2<-rownames(stpresSmallestNAs)[(abs(stpresSmallestNAs)>3)]

#now these need to be removed and the PGLS run again 

#remove from tree
SmallestNAstree3<-drop.tip(SmallestNAstree2, SmallestNAsout2, trim.internal = TRUE)
#remove from data
SmallestNAsdata3<-SmallestNAsdata2[!SmallestNAsdata2$terminal %in% SmallestNAsout2,]
#make new comparative object
SmallestNAscomp3<-comparative.data(phy=SmallestNAstree3, data=SmallestNAsdata3, na.omit = FALSE, names.col = terminal, vcv = TRUE, warn.dropped = TRUE)

#run PGLS 3
pglsSmallestNAs3<-pgls(log(SmallestNAs) ~ log(CW), SmallestNAscomp3, lambda = 'ML')
summary(pglsSmallestNAs3)

#are there any phylogenetic outliers?

#extract phylogenetic residuals
presSmallestNAs<-pglsSmallestNAs3$phyres
#for some reason $residuals takes the rownames with it, but $phyres doesn't?
rownames(presSmallestNAs)<-rownames(pglsSmallestNAs3$residuals)

#studentise
stpresSmallestNAs<-presSmallestNAs/sqrt(var(presSmallestNAs))[1]
#identify species with studentised residuals >|3|
SmallestNAsout3<-rownames(stpresSmallestNAs)[(abs(stpresSmallestNAs)>3)]

#now these need to be removed and the PGLS run again 

#remove from tree
SmallestNAstree4<-drop.tip(SmallestNAstree3, SmallestNAsout3, trim.internal = TRUE)
#remove from data
SmallestNAsdata4<-SmallestNAsdata3[!SmallestNAsdata3$terminal %in% SmallestNAsout3,]
#make new comparative object
SmallestNAscomp4<-comparative.data(phy=SmallestNAstree4, data=SmallestNAsdata4, na.omit = FALSE, names.col = terminal, vcv = TRUE, warn.dropped = TRUE)

#run PGLS 4
pglsSmallestNAs4<-pgls(log(SmallestNAs) ~ log(CW), SmallestNAscomp4, lambda = 'ML')
summary(pglsSmallestNAs4)

#are there any phylogenetic outliers?

#extract phylogenetic residuals
presSmallestNAs<-pglsSmallestNAs4$phyres
#for some reason $residuals takes the rownames with it, but $phyres doesn't?
rownames(presSmallestNAs)<-rownames(pglsSmallestNAs4$residuals)

#studentise
stpresSmallestNAs<-presSmallestNAs/sqrt(var(presSmallestNAs))[1]
#identify species with studentised residuals >|3|
SmallestNAsout4<-rownames(stpresSmallestNAs)[(abs(stpresSmallestNAs)>3)]

#now these need to be removed and the PGLS run again 

#remove from tree
SmallestNAstree5<-drop.tip(SmallestNAstree4, SmallestNAsout4, trim.internal = TRUE)
#remove from data
SmallestNAsdata5<-SmallestNAsdata4[!SmallestNAsdata4$terminal %in% SmallestNAsout4,]
#make new comparative object
SmallestNAscomp5<-comparative.data(phy=SmallestNAstree5, data=SmallestNAsdata5, na.omit = FALSE, names.col = terminal, vcv = TRUE, warn.dropped = TRUE)

#run PGLS 5
pglsSmallestNAs5<-pgls(log(SmallestNAs) ~ log(CW), SmallestNAscomp5, lambda = 'ML')
summary(pglsSmallestNAs5)

#are there any phylogenetic outliers?

#extract phylogenetic residuals
presSmallestNAs<-pglsSmallestNAs5$phyres
#for some reason $residuals takes the rownames with it, but $phyres doesn't?
rownames(presSmallestNAs)<-rownames(pglsSmallestNAs5$residuals)

#residuals
stpresSmallestNAs<-presSmallestNAs/sqrt(var(presSmallestNAs))[1]
#identify species with studentised residuals >|3|
SmallestNAsout5<-rownames(stpresSmallestNAs)[(abs(stpresSmallestNAs)>3)]

#once that's done, we can use $residuals to look at the actual investment

#extract residuals (not phylogenetic residuals!)
resSmallestNAs<-pglsSmallestNAs5$residuals
#studentise
stresSmallestNAs <- resSmallestNAs/sqrt(var(resSmallestNAs))[1]
#identify species with studentised residuals >|3|
rownames(stresSmallestNAs)[(abs(stresSmallestNAs)>3)]

#add the stres as a new column to the original data

#make sure they're in the right order first!
stresSmallestNAs <- stresSmallestNAs[order(row.names(stresSmallestNAs)),]
spider.SmallestNAs.res <- cbind (SmallestNAsdata5, stresSmallestNAs = stresSmallestNAs)

spider.SmallestNAs.res


```

Variance (calculated on relative diameters, where absent eyes are NAs, not =0)

```{r}

#run PGLS 1
pglsrVarNAs<-pgls(rvar ~ log(CW), dfNAs, lambda = 'ML')
summary(pglsrVarNAs)

#are there any phylogenetic outliers?

#extract phylogenetic residuals
presrVarNAs<-pglsrVarNAs$phyres
#for some reason $residuals takes the rownames with it, but $phyres doesn't?
rownames(presrVarNAs)<-rownames(pglsrVarNAs$residuals)

#studentise
stpresrVarNAs<-presrVarNAs/sqrt(var(presrVarNAs))[1]
#identify species with studentised residuals >|3|
rVarNAsout<-rownames(stpresrVarNAs)[(abs(stpresrVarNAs)>3)]

#now these need to be removed and the PGLS run again 

#remove from tree
rVarNAstree2<-drop.tip(tree, rVarNAsout, trim.internal = TRUE)
#remove from data
rVarNAsdata2<-data[!data$terminal %in% rVarNAsout,]
#make new comparative object
rVarNAscomp2<-comparative.data(phy=rVarNAstree2, data=rVarNAsdata2, na.omit = FALSE, names.col = terminal, vcv = TRUE, warn.dropped = TRUE)

#run PGLS 2
pglsrVarNAs2<-pgls(rvar ~ log(CW), rVarNAscomp2, lambda = 'ML')

summary(pglsrVarNAs2)

#are there any phylogenetic outliers?

#extract phylogenetic residuals
presrVarNAs<-pglsrVarNAs2$phyres
#for some reason $residuals takes the rownames with it, but $phyres doesn't?
rownames(presrVarNAs)<-rownames(pglsrVarNAs2$residuals)

#studentise
stpresrVarNAs<-presrVarNAs/sqrt(var(presrVarNAs))[1]
#identify species with studentised residuals >|3|
rVarNAsout2<-rownames(stpresrVarNAs)[(abs(stpresrVarNAs)>3)]

#now these need to be removed and the PGLS run again 

#remove from tree
rVarNAstree3<-drop.tip(rVarNAstree2, rVarNAsout2, trim.internal = TRUE)
#remove from data
rVarNAsdata3<-data[!data$terminal %in% rVarNAsout2,]
#make new comparative object
rVarNAscomp3<-comparative.data(phy=rVarNAstree3, data=rVarNAsdata3, na.omit = FALSE, names.col = terminal, vcv = TRUE, warn.dropped = TRUE)

#run PGLS 2
pglsrVarNAs3<-pgls(rvar ~ log(CW), rVarNAscomp3, lambda = 'ML')
summary(pglsrVarNAs3)

#are there any phylogenetic outliers?

#extract phylogenetic residuals
presrVarNAs<-pglsrVarNAs3$phyres
#for some reason $residuals takes the rownames with it, but $phyres doesn't?
rownames(presrVarNAs)<-rownames(pglsrVarNAs3$residuals)

#studentise
stpresrVarNAs<-presrVarNAs/sqrt(var(presrVarNAs))[1]
#identify species with studentised residuals >|3|
rVarNAsout3<-rownames(stpresrVarNAs)[(abs(stpresrVarNAs)>3)]

#now these need to be removed and the PGLS run again 

#remove from tree
rVarNAstree4<-drop.tip(rVarNAstree3, rVarNAsout3, trim.internal = TRUE)
#remove from data
rVarNAsdata4<-data[!data$terminal %in% rVarNAsout3,]
#make new comparative object
rVarNAscomp4<-comparative.data(phy=rVarNAstree4, data=rVarNAsdata4, na.omit = FALSE, names.col = terminal, vcv = TRUE, warn.dropped = TRUE)

#run PGLS 4
pglsrVarNAs4<-pgls(rvar ~ log(CW), rVarNAscomp4, lambda = 'ML')
summary(pglsrVarNAs4)

#are there any phylogenetic outliers?

#extract phylogenetic residuals
presrVarNAs<-pglsrVarNAs4$phyres
#for some reason $residuals takes the rownames with it, but $phyres doesn't?
rownames(presrVarNAs)<-rownames(pglsrVarNAs4$residuals)

#studentise
stpresrVarNAs<-presrVarNAs/sqrt(var(presrVarNAs))[1]
#identify species with studentised residuals >|3|
rVarNAsout4<-rownames(stpresrVarNAs)[(abs(stpresrVarNAs)>3)]

#now these need to be removed and the PGLS run again 

#remove from tree
rVarNAstree5<-drop.tip(rVarNAstree4, rVarNAsout4, trim.internal = TRUE)
#remove from data
rVarNAsdata5<-data[!data$terminal %in% rVarNAsout4,]
#make new comparative object
rVarNAscomp5<-comparative.data(phy=rVarNAstree5, data=rVarNAsdata5, na.omit = FALSE, names.col = terminal, vcv = TRUE, warn.dropped = TRUE)

#run PGLS 5
pglsrVarNAs5<-pgls(rvar ~ log(CW), rVarNAscomp5, lambda = 'ML')
summary(pglsrVarNAs5)

#are there any phylogenetic outliers?

#extract phylogenetic residuals
presrVarNAs<-pglsrVarNAs5$phyres
#identify species with studentised residuals >|3|
rownames(presrVarNAs)<-rownames(pglsrVarNAs5$residuals)

#studentise
stpresrVarNAs<-presrVarNAs/sqrt(var(presrVarNAs))[1]
#identify species with studentised residuals >|3|
rVarNAsout5<-rownames(stpresrVarNAs)[(abs(stpresrVarNAs)>3)]

#once that's done, we can use $residuals to look at the actual investment

#extract residuals (not phylogenetic residuals!)
resrVarNAs<-pglsrVarNAs5$residuals
#studentise
stresrVarNAs <- resrVarNAs/sqrt(var(resrVarNAs))[1]

#identify species with studentised residuals >|3|
rownames(stresrVarNAs)[(abs(stresrVarNAs)>3)]

#add the stres as a new column to the original data

#make sure they're in the right order first!
stresrVarNAs <- stresrVarNAs[order(row.names(stresrVarNAs)),]
spider.rVarNAs.res <- cbind (rVarNAsdata5, stresrVarNAs = stresrVarNAs)

spider.rVarNAs.res

```

So, following Thomas et al. 2020, we can take the residuals from these first PGLS, with just log(CW) as an explanatory variable, and use these to analyse the impacts of ecological factors via Kruskal Wallis.

```{r dev='pdf'}

kruskal.test(spider.AME.res$stresAME, spider.AME.res$visual.pursuit)
kruskal.test(spider.AME.res$stresAME, spider.AME.res$visual.incl.static)
kruskal.test(spider.AME.res$stresAME, spider.AME.res$builder)
kruskal.test(spider.AME.res$stresAME, spider.AME.res$active)

#ALEs
kruskal.test(spider.ALE.res$stresALE, spider.ALE.res$visual.pursuit)
kruskal.test(spider.ALE.res$stresALE, spider.ALE.res$visual.incl.static)
kruskal.test(spider.ALE.res$stresALE, spider.ALE.res$builder)
kruskal.test(spider.ALE.res$stresALE, spider.ALE.res$active)

#PMEs
kruskal.test(spider.PME.res$stresPME, spider.PME.res$visual.pursuit)
kruskal.test(spider.PME.res$stresPME, spider.PME.res$visual.incl.static)
kruskal.test(spider.PME.res$stresPME, spider.PME.res$builder)
kruskal.test(spider.PME.res$stresPME, spider.PME.res$active)

#PLEs
kruskal.test(spider.PLE.res$stresPLE, spider.PLE.res$visual.pursuit)
kruskal.test(spider.PLE.res$stresPLE, spider.PLE.res$visual.incl.static)
kruskal.test(spider.PLE.res$stresPLE, spider.PLE.res$builder)
kruskal.test(spider.PLE.res$stresPLE, spider.PLE.res$active)

#Largest
kruskal.test(spider.Largest.res$stresLargest, spider.Largest.res$visual.pursuit)
kruskal.test(spider.Largest.res$stresLargest,spider.Largest.res$visual.incl.static)
kruskal.test(spider.Largest.res$stresLargest, spider.Largest.res$builder)
kruskal.test(spider.Largest.res$stresLargest, spider.Largest.res$active)

#Smallest NAs
kruskal.test(spider.SmallestNAs.res$stresSmallestNAs,spider.SmallestNAs.res$visual.pursuit)
kruskal.test(spider.SmallestNAs.res$stresSmallestNAs, spider.SmallestNAs.res$visual.incl.static)
kruskal.test(spider.SmallestNAs.res$stresSmallestNAs, spider.SmallestNAs.res$builder)
kruskal.test(spider.SmallestNAs.res$stresSmallestNAs, spider.SmallestNAs.res$active)

#variance
kruskal.test(spider.rVarNAs.res$stresrVarNAs, spider.rVarNAs.res$visual.pursuit)
kruskal.test(spider.rVarNAs.res$stresrVarNAs, spider.rVarNAs.res$visual.incl.static)
kruskal.test(spider.rVarNAs.res$stresrVarNAs, spider.rVarNAs.res$builder)
kruskal.test(spider.rVarNAs.res$stresrVarNAs, spider.rVarNAs.res$active)

```

Plots for figures 3-5

Figure 3, visual pursuit hunting


```{r}

#AME
#identify tips that don't match the reduced dataset
diff<-setdiff(AMEtree5$tip.label, rownames(resAME))
#remove these
AMEtree6<-drop.tip(AMEtree5,diff,trim.internal = TRUE)

#colour tips
AMEdata5<-AMEdata5 %>%
  mutate(vpcolour=case_when(visual.pursuit=="1" ~ "palevioletred", visual.pursuit=="0" ~ "aquamarine", is.na(visual.pursuit) == TRUE ~ "black"))

vpcol<-as.data.frame(AMEdata5$vpcolour)
rownames(vpcol)<-AMEdata5$terminal
vpcol<-vpcol[order(match(rownames(vpcol), AMEtree6$tip.label)),,]

#plot tree - might need to zoom in to see the colours! Or they're not visible (on my tiny screen, at least)
plotTree.wBars(AMEtree6, resAME[,1], type = "fan", col=vpcol)

#ALE
#identify tips that don't match the reduced dataset
diff<-setdiff(ALEtree5$tip.label, rownames(resALE))
#remove these
ALEtree6<-drop.tip(ALEtree5,diff,trim.internal = TRUE)

#colour tips
ALEdata5<-ALEdata5 %>%
  mutate(vpcolour=case_when(visual.pursuit=="1" ~ "palevioletred", visual.pursuit=="0" ~ "aquamarine", is.na(visual.pursuit) == TRUE ~ "black"))

vpcol<-as.data.frame(ALEdata5$vpcolour)
rownames(vpcol)<-ALEdata5$terminal
vpcol<-vpcol[order(match(rownames(vpcol), ALEtree6$tip.label)),,]

#plot tree - might need to zoom in to see the colours! Or they're not visible (on my tiny screen, at least)
plotTree.wBars(ALEtree6, resALE[,1], type = "fan", col=vpcol)

#PME
#identify tips that don't match the reduced dataset
diff<-setdiff(PMEtree5$tip.label, rownames(resPME))
#remove these
PMEtree6<-drop.tip(PMEtree5,diff,trim.internal = TRUE)

#colour tips
PMEdata5<-PMEdata5 %>%
  mutate(vpcolour=case_when(visual.pursuit=="1" ~ "palevioletred", visual.pursuit=="0" ~ "aquamarine", is.na(visual.pursuit) == TRUE ~ "black"))

vpcol<-as.data.frame(PMEdata5$vpcolour)
rownames(vpcol)<-PMEdata5$terminal
vpcol<-vpcol[order(match(rownames(vpcol), PMEtree6$tip.label)),,]

#plot tree - might need to zoom in to see the colours! Or they're not visible (on my tiny screen, at least)
plotTree.wBars(PMEtree6, resPME[,1], type = "fan", col=vpcol)

#PLE
#identify tips that don't match the reduced dataset
diff<-setdiff(PLEtree5$tip.label, rownames(resPLE))
#remove these
PLEtree6<-drop.tip(PLEtree5,diff,trim.internal = TRUE)

#colour tips
PLEdata5<-PLEdata5 %>%
  mutate(vpcolour=case_when(visual.pursuit=="1" ~ "palevioletred", visual.pursuit=="0" ~ "aquamarine", is.na(visual.pursuit) == TRUE ~ "black"))

vpcol<-as.data.frame(PLEdata5$vpcolour)
rownames(vpcol)<-PLEdata5$terminal
vpcol<-vpcol[order(match(rownames(vpcol), PLEtree6$tip.label)),,]

#plot tree - might need to zoom in to see the colours! Or they're not visible (on my tiny screen, at least)
plotTree.wBars(PLEtree6, resPLE[,1], type = "fan", col=vpcol)

#largest
#identify tips that don't match the reduced dataset
diff<-setdiff(Largesttree5$tip.label, rownames(resLargest))
#remove these
Largesttree6<-drop.tip(Largesttree5,diff,trim.internal = TRUE)

#colour tips
Largestdata5<-Largestdata5 %>%
  mutate(vpcolour=case_when(visual.pursuit=="1" ~ "palevioletred", visual.pursuit=="0" ~ "aquamarine", is.na(visual.pursuit) == TRUE ~ "black"))

vpcol<-as.data.frame(Largestdata5$vpcolour)
rownames(vpcol)<-Largestdata5$terminal
vpcol<-vpcol[order(match(rownames(vpcol), Largesttree6$tip.label)),,]

#plot tree - might need to zoom in to see the colours! Or they're not visible (on my tiny screen, at least)
plotTree.wBars(Largesttree6, resLargest[,1], type = "fan", col=vpcol)

#smallestNAs

#identify tips that don't match the reduced dataset
diff<-setdiff(SmallestNAstree5$tip.label, rownames(resSmallestNAs))
#remove these
SmallestNAstree6<-drop.tip(SmallestNAstree5,diff,trim.internal = TRUE)

#colour tips
SmallestNAsdata5<-SmallestNAsdata5 %>%
  mutate(vpcolour=case_when(visual.pursuit=="1" ~ "palevioletred", visual.pursuit=="0" ~ "aquamarine", is.na(visual.pursuit) == TRUE ~ "black"))

vpcol<-as.data.frame(SmallestNAsdata5$vpcolour)
rownames(vpcol)<-SmallestNAsdata5$terminal
vpcol<-vpcol[order(match(rownames(vpcol), SmallestNAstree6$tip.label)),,]

#plot tree - might need to zoom in to see the colours! Or they're not visible (on my tiny screen, at least)
plotTree.wBars(SmallestNAstree6, resSmallestNAs[,1], type = "fan", col=vpcol)

#rVarNAs
#identify tips that don't match the reduced dataset
diff<-setdiff(rVarNAstree5$tip.label, rownames(resrVarNAs))
#remove these
rVarNAstree6<-drop.tip(rVarNAstree5,diff,trim.internal = TRUE)

#colour tips
rVarNAsdata5<-rVarNAsdata5 %>%
  mutate(vpcolour=case_when(visual.pursuit=="1" ~ "palevioletred", visual.pursuit=="0" ~ "aquamarine", is.na(visual.pursuit) == TRUE ~ "black"))

vpcol<-as.data.frame(rVarNAsdata5$vpcolour)
rownames(vpcol)<-rVarNAsdata5$terminal
vpcol<-vpcol[order(match(rownames(vpcol), rVarNAstree6$tip.label)),,]

#plot tree - might need to zoom in to see the colours! Or they're not visible (on my tiny screen, at least)
plotTree.wBars(rVarNAstree6, resrVarNAs[,1], type = "fan", col=vpcol)

#box plots 
boxplot(spider.AME.res$stresAME ~ spider.AME.res$visual.pursuit)
boxplot(spider.ALE.res$stresALE ~ spider.ALE.res$visual.pursuit)
boxplot(spider.PME.res$stresPME ~ spider.PME.res$visual.pursuit)
boxplot(spider.PLE.res$stresPLE ~ spider.PLE.res$visual.pursuit)
boxplot(spider.Largest.res$stresLargest ~ spider.Largest.res$visual.pursuit)
boxplot(spider.SmallestNAs.res$stresSmallestNAs ~ spider.SmallestNAs.res$visual.pursuit)
boxplot(spider.rVarNAs.res$stresrVarNAs ~ spider.rVarNAs.res$visual.pursuit)

```

Figure 4, web building

```{r}

#AME
#identify tips that don't match the reduced dataset
diff<-setdiff(AMEtree5$tip.label, rownames(resAME))
#remove these
AMEtree6<-drop.tip(AMEtree5,diff,trim.internal = TRUE)

#colour tips
AMEdata5<-AMEdata5 %>%
  mutate(bcolour=case_when(builder=="1" ~ "darkorange1", builder=="0" ~ "dodgerblue", is.na(builder) == TRUE ~ "black"))

bcol<-as.data.frame(AMEdata5$bcolour)
rownames(bcol)<-AMEdata5$terminal
bcol<-bcol[order(match(rownames(bcol), AMEtree6$tip.label)),,]

#plot tree - might need to zoom in to see the colours! Or they're not visible (on my tiny screen, at least)
plotTree.wBars(AMEtree6, resAME[,1], type = "fan", col=bcol)

#ALE
#identify tips that don't match the reduced dataset
diff<-setdiff(ALEtree5$tip.label, rownames(resALE))
#remove these
ALEtree6<-drop.tip(ALEtree5,diff,trim.internal = TRUE)

#colour tips
ALEdata5<-ALEdata5 %>%
  mutate(bcolour=case_when(builder=="1" ~ "darkorange1", builder=="0" ~ "dodgerblue", is.na(builder) == TRUE ~ "black"))

bcol<-as.data.frame(ALEdata5$bcolour)
rownames(bcol)<-ALEdata5$terminal
bcol<-bcol[order(match(rownames(bcol), ALEtree6$tip.label)),,]

#plot tree - might need to zoom in to see the colours! Or they're not visible (on my tiny screen, at least)
plotTree.wBars(ALEtree6, resALE[,1], type = "fan", col=bcol)

#PME
#identify tips that don't match the reduced dataset
diff<-setdiff(PMEtree5$tip.label, rownames(resPME))
diff

PMEtree6<-drop.tip(PMEtree5,diff,trim.internal = TRUE)

#colour tips
PMEdata5<-PMEdata5 %>%
  mutate(bcolour=case_when(builder=="1" ~ "darkorange1", builder=="0" ~ "dodgerblue", is.na(builder) == TRUE ~ "black"))

bcol<-as.data.frame(PMEdata5$bcolour)
rownames(bcol)<-PMEdata5$terminal
bcol<-bcol[order(match(rownames(bcol), PMEtree6$tip.label)),,]

#plot tree - might need to zoom in to see the colours! Or they're not visible (on my tiny screen, at least)
plotTree.wBars(PMEtree6, resPME[,1], type = "fan", col=bcol)

#PLE
#identify tips that don't match the reduced dataset
diff<-setdiff(PLEtree5$tip.label, rownames(resPLE))
#remove these
PLEtree6<-drop.tip(PLEtree5,diff,trim.internal = TRUE)

#colour tips
PLEdata5<-PLEdata5 %>%
  mutate(bcolour=case_when(builder=="1" ~ "darkorange1", builder=="0" ~ "dodgerblue", is.na(builder) == TRUE ~ "black"))

bcol<-as.data.frame(PLEdata5$bcolour)
rownames(bcol)<-PLEdata5$terminal
bcol<-bcol[order(match(rownames(bcol), PLEtree6$tip.label)),,]

#plot tree - might need to zoom in to see the colours! Or they're not visible (on my tiny screen, at least)
plotTree.wBars(PLEtree6, resPLE[,1], type = "fan", col=bcol)

#largest
#identify tips that don't match the reduced dataset
diff<-setdiff(Largesttree5$tip.label, rownames(resLargest))
#remove these
Largesttree6<-drop.tip(Largesttree5,diff,trim.internal = TRUE)

#colour tips
Largestdata5<-Largestdata5 %>%
  mutate(bcolour=case_when(builder=="1" ~ "darkorange1", builder=="0" ~ "dodgerblue", is.na(builder) == TRUE ~ "black"))

bcol<-as.data.frame(Largestdata5$bcolour)
rownames(bcol)<-Largestdata5$terminal
bcol<-bcol[order(match(rownames(bcol), Largesttree6$tip.label)),,]

#plot tree - might need to zoom in to see the colours! Or they're not visible (on my tiny screen, at least)
plotTree.wBars(Largesttree6, resLargest[,1], type = "fan", col=bcol)

#smallestNAs

#identify tips that don't match the reduced dataset
diff<-setdiff(SmallestNAstree5$tip.label, rownames(resSmallestNAs))
#remove these
SmallestNAstree6<-drop.tip(SmallestNAstree5,diff,trim.internal = TRUE)

#colour tips
SmallestNAsdata5<-SmallestNAsdata5 %>%
  mutate(bcolour=case_when(builder=="1" ~ "darkorange1", builder=="0" ~ "dodgerblue", is.na(builder) == TRUE ~ "black"))

bcol<-as.data.frame(SmallestNAsdata5$bcolour)
rownames(bcol)<-SmallestNAsdata5$terminal
bcol<-bcol[order(match(rownames(bcol), SmallestNAstree6$tip.label)),,]

#plot tree - might need to zoom in to see the colours! Or they're not visible (on my tiny screen, at least)
plotTree.wBars(SmallestNAstree6, resSmallestNAs[,1], type = "fan", col=bcol)

#rVarNAs
#identify tips that don't match the reduced dataset
diff<-setdiff(rVarNAstree5$tip.label, rownames(resrVarNAs))
#remove these
rVarNAstree6<-drop.tip(rVarNAstree5,diff,trim.internal = TRUE)

#colour tips
rVarNAsdata5<-rVarNAsdata5 %>%
  mutate(bcolour=case_when(builder=="1" ~ "darkorange1", builder=="0" ~ "dodgerblue", is.na(builder) == TRUE ~ "black"))

bcol<-as.data.frame(rVarNAsdata5$bcolour)
rownames(bcol)<-rVarNAsdata5$terminal
bcol<-bcol[order(match(rownames(bcol), rVarNAstree6$tip.label)),,]

#plot tree - might need to zoom in to see the colours! Or they're not visible (on my tiny screen, at least)
plotTree.wBars(rVarNAstree6, resrVarNAs[,1], type = "fan", col=bcol)

#box plots 
boxplot(spider.AME.res$stresAME ~ spider.AME.res$builder)
boxplot(spider.ALE.res$stresALE ~ spider.ALE.res$builder)
boxplot(spider.PME.res$stresPME ~ spider.PME.res$builder)
boxplot(spider.PLE.res$stresPLE ~ spider.PLE.res$builder)
boxplot(spider.Largest.res$stresLargest ~ spider.Largest.res$builder)
boxplot(spider.SmallestNAs.res$stresSmallestNAs ~ spider.SmallestNAs.res$builder)
boxplot(spider.rVarNAs.res$stresrVarNAs ~ spider.rVarNAs.res$builder)

```

Figure 5, activity

```{r}

#AME
#identify tips that don't match the reduced dataset
diff<-setdiff(AMEtree5$tip.label, rownames(resAME))

#remove these
AMEtree6<-drop.tip(AMEtree5,diff,trim.internal = TRUE)

#colour tips
AMEdata5<-AMEdata5 %>%
  mutate(actcolour=case_when(active=="1" ~ "darkgoldenrod1", active=="0" ~ "blueviolet", is.na(active) == TRUE ~ "black"))

actcol<-as.data.frame(AMEdata5$actcolour)
rownames(actcol)<-AMEdata5$terminal
actcol<-actcol[order(match(rownames(actcol), AMEtree6$tip.label)),,]

#plot tree - might need to zoom in to see the colours! Or they're not visible (on my tiny screen, at least)
plotTree.wBars(AMEtree6, resAME[,1], type = "fan", col=actcol)

#ALE
#identify tips that don't match the reduced dataset
diff<-setdiff(ALEtree5$tip.label, rownames(resALE))

#remove these
ALEtree6<-drop.tip(ALEtree5,diff,trim.internal = TRUE)

#colour tips
ALEdata5<-ALEdata5 %>%
  mutate(actcolour=case_when(active=="1" ~ "darkgoldenrod1", active=="0" ~ "blueviolet", is.na(active) == TRUE ~ "black"))

actcol<-as.data.frame(ALEdata5$actcolour)
rownames(actcol)<-ALEdata5$terminal
actcol<-actcol[order(match(rownames(actcol), ALEtree6$tip.label)),,]

#plot tree - might need to zoom in to see the colours! Or they're not visible (on my tiny screen, at least)
plotTree.wBars(ALEtree6, resALE[,1], type = "fan", col=actcol)

#PME
#identify tips that don't match the reduced dataset
diff<-setdiff(PMEtree5$tip.label, rownames(resPME))

#remove these
PMEtree6<-drop.tip(PMEtree5,diff,trim.internal = TRUE)

#colour tips
PMEdata5<-PMEdata5 %>%
  mutate(actcolour=case_when(active=="1" ~ "darkgoldenrod1", active=="0" ~ "blueviolet", is.na(active) == TRUE ~ "black"))

actcol<-as.data.frame(PMEdata5$actcolour)
rownames(actcol)<-PMEdata5$terminal
actcol<-actcol[order(match(rownames(actcol), PMEtree6$tip.label)),,]

#plot tree - might need to zoom in to see the colours! Or they're not visible (on my tiny screen, at least)
plotTree.wBars(PMEtree6, resPME[,1], type = "fan", col=actcol)

#PLE
#identify tips that don't match the reduced dataset
diff<-setdiff(PLEtree5$tip.label, rownames(resPLE))

#remove these
PLEtree6<-drop.tip(PLEtree5,diff,trim.internal = TRUE)

#colour tips
PLEdata5<-PLEdata5 %>%
  mutate(actcolour=case_when(active=="1" ~ "darkgoldenrod1", active=="0" ~ "blueviolet", is.na(active) == TRUE ~ "black"))

actcol<-as.data.frame(PLEdata5$actcolour)
rownames(actcol)<-PLEdata5$terminal
actcol<-actcol[order(match(rownames(actcol), PLEtree6$tip.label)),,]

#plot tree - might need to zoom in to see the colours! Or they're not visible (on my tiny screen, at least)
plotTree.wBars(PLEtree6, resPLE[,1], type = "fan", col=actcol)

#largest
#identify tips that don't match the reduced dataset
diff<-setdiff(Largesttree5$tip.label, rownames(resLargest))
#remove these 
Largesttree6<-drop.tip(Largesttree5,diff,trim.internal = TRUE)

#colour tips
Largestdata5<-Largestdata5 %>%
  mutate(actcolour=case_when(active=="1" ~ "darkgoldenrod1", active=="0" ~ "blueviolet", is.na(active) == TRUE ~ "black"))

actcol<-as.data.frame(Largestdata5$actcolour)
rownames(actcol)<-Largestdata5$terminal
actcol<-actcol[order(match(rownames(actcol), Largesttree6$tip.label)),,]

#plot tree - might need to zoom in to see the colours! Or they're not visible (on my tiny screen, at least)
plotTree.wBars(Largesttree6, resLargest[,1], type = "fan", col=actcol)

#smallestNAs

#identify tips that don't match the reduced dataset
diff<-setdiff(SmallestNAstree5$tip.label, rownames(resSmallestNAs))

#remove these
SmallestNAstree6<-drop.tip(SmallestNAstree5,diff,trim.internal = TRUE)

#colour tips
SmallestNAsdata5<-SmallestNAsdata5 %>%
  mutate(actcolour=case_when(active=="1" ~ "darkgoldenrod1", active=="0" ~ "blueviolet", is.na(active) == TRUE ~ "black"))

actcol<-as.data.frame(SmallestNAsdata5$actcolour)
rownames(actcol)<-SmallestNAsdata5$terminal
actcol<-actcol[order(match(rownames(actcol), SmallestNAstree6$tip.label)),,]

#plot tree - might need to zoom in to see the colours! Or they're not visible (on my tiny screen, at least)
plotTree.wBars(SmallestNAstree6, resSmallestNAs[,1], type = "fan", col=actcol)

#rVarNAs
#identify tips that don't match the reduced dataset
diff<-setdiff(rVarNAstree4$tip.label, rownames(resrVarNAs))

#remove these
rVarNAstree6<-drop.tip(rVarNAstree5,diff,trim.internal = TRUE)

#colour tips
rVarNAsdata5<-rVarNAsdata5 %>%
  mutate(actcolour=case_when(active=="1" ~ "darkgoldenrod1", active=="0" ~ "blueviolet", is.na(active) == TRUE ~ "black"))

actcol<-as.data.frame(rVarNAsdata5$actcolour)
rownames(actcol)<-rVarNAsdata5$terminal
actcol<-actcol[order(match(rownames(actcol), rVarNAstree6$tip.label)),,]

#plot tree - might need to zoom in to see the colours! Or they're not visible (on my tiny screen, at least)
plotTree.wBars(rVarNAstree6, resrVarNAs[,1], type = "fan", col=actcol)

#box plots 
boxplot(spider.AME.res$stresAME ~ spider.AME.res$active)
boxplot(spider.ALE.res$stresALE ~ spider.ALE.res$active)
boxplot(spider.PME.res$stresPME ~ spider.PME.res$active)
boxplot(spider.PLE.res$stresPLE ~ spider.PLE.res$active)
boxplot(spider.Largest.res$stresLargest ~ spider.Largest.res$active)
boxplot(spider.SmallestNAs.res$stresSmallestNAs ~ spider.SmallestNAs.res$active)
boxplot(spider.rVarNAs.res$stresrVarNAs ~ spider.rVarNAs.res$active)

```

Not included as a figure, but same for Visual incl static

```{r}

#AME
#identify tips that don't match the reduced dataset
diff<-setdiff(AMEtree5$tip.label, rownames(resAME))

#remove these
AMEtree6<-drop.tip(AMEtree5,diff,trim.internal = TRUE)

#colour tips
AMEdata5<-AMEdata5 %>%
  mutate(vscolour=case_when(visual.incl.static=="1" ~ "deeppink", visual.incl.static=="0" ~ "darkturquoise", is.na(visual.incl.static) == TRUE ~ "black"))

vscol<-as.data.frame(AMEdata5$vscolour)
rownames(vscol)<-AMEdata5$terminal
vscol<-vscol[order(match(rownames(vscol), AMEtree6$tip.label)),,]

#plot tree - might need to zoom in to see the colours! Or they're not visible (on my tiny screen, at least)
plotTree.wBars(AMEtree6, resAME[,1], type = "fan", col=vscol)

#ALE
#identify tips that don't match the reduced dataset
diff<-setdiff(ALEtree5$tip.label, rownames(resALE))

#remove these
ALEtree6<-drop.tip(ALEtree5,diff,trim.internal = TRUE)

#colour tips
ALEdata5<-ALEdata5 %>%
  mutate(vscolour=case_when(visual.incl.static=="1" ~ "deeppink", visual.incl.static=="0" ~ "darkturquoise", is.na(visual.incl.static) == TRUE ~ "black"))

vscol<-as.data.frame(ALEdata5$vscolour)
rownames(vscol)<-ALEdata5$terminal
vscol<-vscol[order(match(rownames(vscol), ALEtree6$tip.label)),,]

#plot tree - might need to zoom in to see the colours! Or they're not visible (on my tiny screen, at least)
plotTree.wBars(ALEtree6, resALE[,1], type = "fan", col=vscol)

#PME
#identify tips that don't match the reduced dataset
diff<-setdiff(PMEtree5$tip.label, rownames(resPME))
#remove these
PMEtree6<-drop.tip(PMEtree5,diff,trim.internal = TRUE)

#colour tips
PMEdata5<-PMEdata5 %>%
  mutate(vscolour=case_when(visual.incl.static=="1" ~ "deeppink", visual.incl.static=="0" ~ "darkturquoise", is.na(visual.incl.static) == TRUE ~ "black"))

vscol<-as.data.frame(PMEdata5$vscolour)
rownames(vscol)<-PMEdata5$terminal
vscol<-vscol[order(match(rownames(vscol), PMEtree6$tip.label)),,]

#plot tree - might need to zoom in to see the colours! Or they're not visible (on my tiny screen, at least)
plotTree.wBars(PMEtree6, resPME[,1], type = "fan", col=vscol)

#PLE
#identify tips that don't match the reduced dataset
diff<-setdiff(PLEtree5$tip.label, rownames(resPLE))
#remove these
PLEtree6<-drop.tip(PLEtree5,diff,trim.internal = TRUE)

#colour tips
PLEdata5<-PLEdata5 %>%
  mutate(vscolour=case_when(visual.incl.static=="1" ~ "deeppink", visual.incl.static=="0" ~ "darkturquoise", is.na(visual.incl.static) == TRUE ~ "black"))

vscol<-as.data.frame(PLEdata5$vscolour)
rownames(vscol)<-PLEdata5$terminal
vscol<-vscol[order(match(rownames(vscol), PLEtree6$tip.label)),,]

#plot tree - might need to zoom in to see the colours! Or they're not visible (on my tiny screen, at least)
plotTree.wBars(PLEtree6, resPLE[,1], type = "fan", col=vscol)

#largest
#identify tips that don't match the reduced dataset
diff<-setdiff(Largesttree5$tip.label, rownames(resLargest))
#remove these
Largesttree6<-drop.tip(Largesttree5,diff,trim.internal = TRUE)

#colour tips
Largestdata5<-Largestdata5 %>%
  mutate(vscolour=case_when(visual.incl.static=="1" ~ "deeppink", visual.incl.static=="0" ~ "darkturquoise", is.na(visual.incl.static) == TRUE ~ "black"))

vscol<-as.data.frame(Largestdata5$vscolour)
rownames(vscol)<-Largestdata5$terminal
vscol<-vscol[order(match(rownames(vscol), Largesttree6$tip.label)),,]

#plot tree - might need to zoom in to see the colours! Or they're not visible (on my tiny screen, at least)
plotTree.wBars(Largesttree6, resLargest[,1], type = "fan", col=vscol)

#smallestNAs

#identify tips that don't match the reduced dataset
diff<-setdiff(SmallestNAstree5$tip.label, rownames(resSmallestNAs))
#remove these
SmallestNAstree6<-drop.tip(SmallestNAstree5,diff,trim.internal = TRUE)

#colour tips
SmallestNAsdata5<-SmallestNAsdata5 %>%
  mutate(vscolour=case_when(visual.incl.static=="1" ~ "deeppink", visual.incl.static=="0" ~ "darkturquoise", is.na(visual.incl.static) == TRUE ~ "black"))

vscol<-as.data.frame(SmallestNAsdata5$vscolour)
rownames(vscol)<-SmallestNAsdata5$terminal
vscol<-vscol[order(match(rownames(vscol), SmallestNAstree6$tip.label)),,]

#plot tree - might need to zoom in to see the colours! Or they're not visible (on my tiny screen, at least)
plotTree.wBars(SmallestNAstree6, resSmallestNAs[,1], type = "fan", col=vscol)

#rVarNAs
#identify tips that don't match the reduced dataset
diff<-setdiff(rVarNAstree5$tip.label, rownames(resrVarNAs))
#remove these
rVarNAstree6<-drop.tip(rVarNAstree5,diff,trim.internal = TRUE)

#colour tips
rVarNAsdata5<-rVarNAsdata5 %>%
  mutate(vscolour=case_when(visual.incl.static=="1" ~ "deeppink", visual.incl.static=="0" ~ "darkturquoise", is.na(visual.incl.static) == TRUE ~ "black"))

vscol<-as.data.frame(rVarNAsdata5$vscolour)
rownames(vscol)<-rVarNAsdata5$terminal
vscol<-vscol[order(match(rownames(vscol), rVarNAstree6$tip.label)),,]

#plot tree - might need to zoom in to see the colours! Or they're not visible (on my tiny screen, at least)
plotTree.wBars(rVarNAstree6, resrVarNAs[,1], type = "fan", col=vscol)

#box plots 
boxplot(spider.AME.res$stresAME ~ spider.AME.res$visual.incl.static)
boxplot(spider.ALE.res$stresALE ~ spider.ALE.res$visual.incl.static)
boxplot(spider.PME.res$stresPME ~ spider.PME.res$visual.incl.static)
boxplot(spider.PLE.res$stresPLE ~ spider.PLE.res$visual.incl.static)
boxplot(spider.Largest.res$stresLargest ~ spider.Largest.res$visual.incl.static)
boxplot(spider.SmallestNAs.res$stresSmallestNAs ~ spider.SmallestNAs.res$visual.incl.static)
boxplot(spider.rVarNAs.res$stresrVarNAs ~ spider.rVarNAs.res$visual.incl.static)


```


Alternative approach: using ecological factors as covariates (phylogenetic ANCOVA, after Thomas et al. 2020)
NB. uses datasets with phylogenetic outliers already removed, from fifth iterations above.

1. visual pursuit element to hunting known

```{r}

pglsAMEV<-pgls(log(AME) ~ log(CW) * visual.pursuit, AMEcomp5, lambda = 'ML')
summary(pglsAMEV)

pglsALEV<-pgls(log(ALE) ~ log(CW)* visual.pursuit, ALEcomp5, lambda = 'ML')
summary(pglsALEV)

pglsPMEV<-pgls(log(PME) ~ log(CW)* visual.pursuit, PMEcomp3, lambda = 'ML')
summary(pglsPMEV)

pglsPLEV<-pgls(log(PLE) ~ log(CW)* visual.pursuit, PLEcomp5, lambda = 'ML')
summary(pglsPLEV)

pglsLargestV<-pgls(log(Largest) ~ log(CW)* visual.pursuit, Largestcomp5, lambda = 'ML')
summary(pglsLargestV)

pglsSmallestNAsV<-pgls(log(Smallest) ~ log(CW)* visual.pursuit, SmallestNAscomp5, lambda = 'ML')
summary(pglsSmallestNAsV)

pglsrVarNAsV<-pgls(rvar ~ log(CW) * visual.pursuit, rVarNAscomp3, lambda = 'ML')
summary(pglsrVarNAsV)

#for variance, also tried without CW
pglsrVarNAsVOnly<-pgls(rvar ~ visual.pursuit, rVarNAscomp3, lambda = 'ML')
summary(pglsrVarNAsVOnly)

#and is CW affected by factor? Phylo. logistic regression
pglsCWVOnly<-pgls(log(CW) ~ visual.pursuit, Largestcomp5, lambda = 'ML')
summary(pglsCWVOnly)

```

2.All species with a visual element to hunting, including static visual sit-and-wait

```{r}

pglsAMEVs<-pgls(log(AME) ~ log(CW) * visual.incl.static, AMEcomp5, lambda = 'ML')
summary(pglsAMEVs)

pglsALEVs<-pgls(log(ALE) ~ log(CW)* visual.incl.static, ALEcomp5, lambda = 'ML')
summary(pglsALEVs)

pglsPMEVs<-pgls(log(PME) ~ log(CW)* visual.incl.static, PMEcomp3, lambda = 'ML')
summary(pglsPMEVs)

pglsPLEVs<-pgls(log(PLE) ~ log(CW)* visual.incl.static, PLEcomp5, lambda = 'ML')
summary(pglsPLEVs)

pglsLargestVs<-pgls(log(Largest) ~ log(CW)* visual.incl.static, Largestcomp5, lambda = 'ML')
summary(pglsLargestVs)

pglsSmallestVs<-pgls(log(Smallest) ~ log(CW)* visual.incl.static, Smallestcomp5, lambda = 'ML')
summary(pglsSmallestVs)

pglsSmallestNAsVs<-pgls(log(Smallest) ~ log(CW)* visual.incl.static, SmallestNAscomp5, lambda = 'ML')
summary(pglsSmallestNAsVs)

pglsrVarNAsVs<-pgls(rvar ~ log(CW) * visual.incl.static, rVarNAscomp3, lambda = 'ML')
summary(pglsrVarNAsVs)

#for variance, also tried without CW
pglsrVarNAsVOnly<-pgls(rvar ~ visual.incl.static, rVarNAscomp3, lambda = 'ML')
summary(pglsrVarNAsVOnly)

#and is CW affected by factor? Phylo. logistic regression
pglsCWVsOnly<-pgls(log(CW) ~ visual.incl.static, Largestcomp5, lambda = 'ML')
summary(pglsCWVsOnly)

```

3.  Web building

```{r}

pglsAMEB<-pgls(log(AME) ~ log(CW) * builder, AMEcomp5, lambda = 'ML')
summary(pglsAMEB)

pglsALEB<-pgls(log(ALE) ~ log(CW)* builder, ALEcomp5, lambda = 'ML')
summary(pglsALEB)

pglsPMEB<-pgls(log(PME) ~ log(CW)* builder, PMEcomp3, lambda = 'ML')
summary(pglsPMEB)

pglsPLEB<-pgls(log(PLE) ~ log(CW)* builder, PLEcomp5, lambda = 'ML')
summary(pglsPLEB)

pglsLargestB<-pgls(log(Largest) ~ log(CW)* builder, Largestcomp5, lambda = 'ML')
summary(pglsLargestB)

pglsSmallestNAsB<-pgls(log(Smallest) ~ log(CW)* builder, SmallestNAscomp5, lambda = 'ML')
summary(pglsSmallestNAsB)

pglsrVarNAsB<-pgls(rvar ~ log(CW) * builder, rVarNAscomp3, lambda = 'ML')
summary(pglsrVarNAsB)

#for variance, also tried without CW
pglsrVarNAsBOnly<-pgls(rvar ~ builder, rVarNAscomp3, lambda = 'ML')
summary(pglsrVarNAsBOnly)

#and is CW affected by factor? Phylo. logistic regression
pglsCWBOnly<-pgls(log(CW) ~ builder, Largestcomp5, lambda = 'ML')
summary(pglsCWBOnly)

```

4.  activity

```{r}

pglsAMEA<-pgls(log(AME) ~ log(CW) * active, AMEcomp5, lambda = 'ML')
summary(pglsAMEA)

pglsALEA<-pgls(log(ALE) ~ log(CW)* active, ALEcomp5, lambda = 'ML')
summary(pglsALEA)

pglsPMEA<-pgls(log(PME) ~ log(CW)* active, PMEcomp3, lambda = 'ML')
summary(pglsPMEA)

pglsPLEA<-pgls(log(PLE) ~ log(CW)* active, PLEcomp5, lambda = 'ML')
summary(pglsPLEA)

pglsLargestA<-pgls(log(Largest) ~ log(CW)* active, Largestcomp5, lambda = 'ML')
summary(pglsLargestA)

pglsSmallestNAsA<-pgls(log(Smallest) ~ log(CW)* active, SmallestNAscomp5, lambda = 'ML')
summary(pglsSmallestNAsA)

pglsrVarNAsA<-pgls(rvar ~ log(CW) * active, rVarNAscomp3, lambda = 'ML')
summary(pglsrVarNAsA)

#for variance, also tried without CW
pglsrVarNAsAOnly<-pgls(rvar ~ active, rVarNAscomp3, lambda = 'ML')
summary(pglsrVarNAsAOnly)

#and is CW affected by factor? Phylo. logistic regression
pglsCWAOnly<-pgls(log(CW) ~ active, Largestcomp5, lambda = 'ML')
summary(pglsCWAOnly)

```
````
