## Supplementary material for "Complex allometric relationships and ecological factors shape the development and evolution of eye size in the modular visual system of spiders": README.txt

Chong et al. Supplementary Materials

1) Datasets

ln_total.csv = measured eye diameters and carapace widths from OUMNH collections for static allometry. These data are called in static_allometry.Rmd
WolffData-Dec23.txt = curated dataset from Wolff et al. 2022 Syst. Biol. used for phylogenetic allometry. Absent eyes are assigned a diameter of 0. These data are called in PGLS.Rmd
WolffDataNAs-Dec23.txt = curated dataset from Wolff et al. 2022 Syst. Biol. used for phylogenetic allometry. Absent eyes are NAs. These data are called in PGLS.Rmd
WolffRed22Sept.nwk = curated phylogenetic tree from Wolff et al. 2022 Syst. Biol. used for phylogenetic allometry. These data are called in PGLS.Rmd

2) Analyses

static_allometry.Rmd = analysis of measured eye diameters and carapace widths from OUMNH collections for static allometry. Calls ln_total.csv
PGLS.Rmd = analysis of curated Wolff et al. 2022 dataset for phylogenetic allometry. Calls WolffData-Dec23.txt, WolffDataNAs-Dec23.txt, and WolffRed22Sept.nwk
ancestral_state_analysis.Rmd = analysis of curated Wolff et al. 2022 dataset for ancestral state reconstruction. Calls WolffDataNAs-Dec23.txt and WolffRed22Sept.nwk
Bayou Rmd.zip = contains datasets, analyses, and results of Bayou allometric shift estimation

3) Results

Supplementary_tables.xls = contains supplementary tables referred to in the manuscript.
