## Supplementary material for "Complex allometric relationships and ecological factors shape the development and evolution of eye size in the modular visual system of spiders": static_allometry.Rmd

````
---
title: "R Analysis"
output: html_notebook
editor_options: 
  chunk_output_type: console
---

```{r, echo=FALSE}
#Required packages
pacman::p_load(tidyverse,cowplot,RColorBrewer, dplyr,lme4, emmeans, effects, forcats, lmerTest, ggplot2, emmeans, broom.mixed, rlist, ape, nlme, geiger, ade4, gpubr, bayou, magrittr, tidyr, caper, phytools,tidyr)
#reading in data
total <-read.csv('ln_total.csv')
total <- as_tibble(total)
```

Making the Caper file

```{r}
#creates a grouped data frame based on the columns Family, Species, and Number. 
group_number <- group_by(total, Family, Species, Number)
#calculates mean and log carapace width
specimen_mean <- summarise(group_number, mean_CW = mean(CW), log_CW = mean(log_CW), na.rm = TRUE)

species_mean <- specimen_mean %>% group_by(Family, Species) %>%
  summarise(CARAPACE = mean(mean_CW),logCW = mean(log_CW), na.rm = TRUE)

#creates a grouped data frame that has mean and log eye pair diameter
eyepairs <- group_by(total, Family, Species, Eye.pair)
abs_eyepairs<- summarise(eyepairs, abs = mean(Absolute), rel = mean(Relative), logEP = mean(log_EP), na.rm = TRUE, .groups = 'drop')

#merge the carapace and eye pair means into one dataframe
total_data <- merge(species_mean, abs_eyepairs, by = c("Family", "Species"))

#clean up dataframe
tot <- total_data[,-5]
tot <- tot[,-9]

tot <- as_tibble(tot)

df <- tot %>%
  pivot_wider(names_from =Eye.pair, values_from = c(abs, rel, logEP))
#save the caper file to be used later
write.csv(df,"caper_draft.csv" )
```

# **Eye size and body size raw data**

```{r}
#create a box and whisker plot showing absolute eye size separated per family
family_plot <- ggplot(total, aes(x=Family, y=log_EP, fill = Eye.pair)) + geom_boxplot() + scale_fill_manual(values =c('#eded15','#c360dc', '#e20505', '#09ef09')) + labs(x = "Family", y = 'Eye diameter') + 
  guides(fill=guide_legend(title="Eye pair")) + theme_light() 
ggsave(family_plot, file=paste0("Absolute.pdf"), width = 44.45, 
         height = 27.78, units = "cm", dpi=300)

#create a box and whisker plot showing relative eye size separated per family
relative_diameter <- ggplot(total, aes(x=Family, y= Relative, fill = Eye.pair)) + geom_boxplot() + 
  scale_fill_manual(values =c('#eded15','#c360dc', '#e20505', '#09ef09')) + labs(x = "Family", y = 'Relative eye diameter') + 
  guides(fill=guide_legend(title="Eye pair")) + theme_light()
ggsave(relative_diameter, file=paste0("Relative.pdf"), width = 44.45, 
         height = 27.78, units = "cm", dpi=300)

#create a box and whisker plot showing carapace width separated per family
carapace <- ggplot(total, aes(x=Family, y= CW)) + geom_boxplot() + labs(y= "Carapace width") + theme_light()
ggsave(carapace, file=paste0("Carapace.pdf"), width = 44.45, 
         height = 27.78, units = "cm", dpi=300)
```


#Running Anovas
```{r}
library(tidyverse)
library(ggpubr)
library(rstatix)

##########PHOLCID ############
fam_dat = total_df %>% filter(Family == "Pholcid")
#check for outliers
outliers <- fam_dat %>% group_by(Eye.pair) %>%
  identify_outliers(log_EP)
fam_dat <- fam_dat %>% anti_join(outliers, by = c("Eye.pair", "log_EP"))
fam_dat %>% group_by(Eye.pair) %>%
  identify_outliers(log_EP)

fam_dat <- as.data.frame(fam_dat)
##Analyzing the ANOVA model residuals 
#Build the linear model
model  <- lm(log_EP ~ Eye.pair, data = fam_dat)
# Create a QQ plot of residuals
ggqqplot(residuals(model))
model <- aov(log_EP ~ Eye.pair, data = fam_dat)
# Compute Shapiro-Wilk test of normality
shapiro_test(residuals(model))

#Check normality for each group separately - was unconvinced by pholcid
fam_dat %>% group_by(Eye.pair) %>%
  shapiro_test(log_EP)
ggqqplot(fam_dat, "log_EP", facet.by = "Eye.pair")

#Homogneity of variance
#The residuals versus fits plot can be used to check the homogeneity of variances.
plot(model, 1)

#compute
res.aov <- aov(log_EP ~ Eye.pair, data = fam_dat)
res.aov
report(res.aov)

tukey_result <- TukeyHSD(res.aov)
tukey_result

###########ARANEIDAE #########
fam_dat = total_df %>% filter(Family == "Araneidae")
#check for outliers
outliers <- fam_dat %>% group_by(Eye.pair) %>%
  identify_outliers(log_EP)
fam_dat <- fam_dat %>% anti_join(outliers, by = c("Eye.pair", "log_EP"))
fam_dat %>% group_by(Eye.pair) %>%
  identify_outliers(log_EP)

fam_dat <- as.data.frame(fam_dat)
#check for normality
##Analyzing the ANOVA model residuals 
#Build the linear model
model  <- lm(log_EP ~ Eye.pair, data = fam_dat)
# Create a QQ plot of residuals
ggqqplot(residuals(model))
model <- aov(log_EP ~ Eye.pair, data = fam_dat)
# Compute Shapiro-Wilk test of normality
shapiro_test(residuals(model))
#DOES NOT MEET NORMALITY ASSUMPTION

#Homogneity of variance
#The residuals versus fits plot can be used to check the homogeneity of variances.
plot(model, 1)

#compute
logEP <- as.numeric(fam_dat$log_EP)
eyepair <- factor(c(fam_dat$Eye.pair))
KWmodel <- kruskal.test(logEP~eyepair)
KWmodel
#significant, run Dunns
dunn_result <- dunnTest(logEP ~ eyepair,method="bonferroni")
dunn_result
##########GNAPHOSIDAE ###########
fam_dat = total_df %>% filter(Family == "Gnaphosidae")
#check for outliers
outliers <- fam_dat %>% group_by(Eye.pair) %>%
  identify_outliers(log_EP)
fam_dat <- fam_dat %>% anti_join(outliers, by = c("Eye.pair", "log_EP"))
fam_dat %>% group_by(Eye.pair) %>%
  identify_outliers(log_EP)

fam_dat <- as.data.frame(fam_dat)
#check for normality
##Analyzing the ANOVA model residuals 
#Build the linear model
model  <- lm(log_EP ~ Eye.pair, data = fam_dat)
# Create a QQ plot of residuals
ggqqplot(residuals(model))
model <- aov(log_EP ~ Eye.pair, data = fam_dat)
# Compute Shapiro-Wilk test of normality
shapiro_test(residuals(model))
#DOES NOT MEET NORMALITY ASSUMPTION

#Homogneity of variance
#The residuals versus fits plot can be used to check the homogeneity of variances.
plot(model, 1)
#compute
logEP <- as.numeric(fam_dat$log_EP)
eyepair <- factor(c(fam_dat$Eye.pair))
KWmodel <- kruskal.test(logEP~eyepair)
KWmodel
#significant, run Dunns
dunn_result <- dunnTest(logEP ~ eyepair,method="bonferroni")
dunn_result
###########THERIDIIDAE #########
fam_dat = total_df %>% filter(Family == "Theridiidae")
#check for outliers
outliers <- fam_dat %>% group_by(Eye.pair) %>%
  identify_outliers(log_EP)
fam_dat <- fam_dat %>% anti_join(outliers, by = c("Eye.pair", "log_EP"))
fam_dat %>% group_by(Eye.pair) %>%
  identify_outliers(log_EP)

fam_dat <- as.data.frame(fam_dat)
#check for normality
##Analyzing the ANOVA model residuals 
#Build the linear model
model  <- lm(log_EP ~ Eye.pair, data = fam_dat)
# Create a QQ plot of residuals
ggqqplot(residuals(model))
model <- aov(log_EP ~ Eye.pair, data = fam_dat)
# Compute Shapiro-Wilk test of normality
shapiro_test(residuals(model))
#DOES NOT MEET NORMALITY ASSUMPTION

#Homogneity of variance
#The residuals versus fits plot can be used to check the homogeneity of variances.
plot(model, 1)

#compute
logEP <- as.numeric(fam_dat$log_EP)
eyepair <- factor(c(fam_dat$Eye.pair))
KWmodel <- kruskal.test(logEP~eyepair)
KWmodel
#significant, run Dunns
dunn_result <- dunnTest(logEP ~ eyepair,method="bonferroni")
dunn_result

###########SALTICID#############
fam_dat = total_df %>% filter(Family == "Salticidae")

#check for outliers
outliers <- fam_dat %>% group_by(Eye.pair) %>%
  identify_outliers(log_EP)
fam_dat <- fam_dat %>% anti_join(outliers, by = c("Eye.pair", "log_EP"))
fam_dat %>% group_by(Eye.pair) %>%
  identify_outliers(log_EP)

fam_dat <- as.data.frame(fam_dat)
#check for normality
##Analyzing the ANOVA model residuals 
#Build the linear model
model  <- lm(log_EP ~ Eye.pair, data = fam_dat)
# Create a QQ plot of residuals
ggqqplot(residuals(model))
model <- aov(log_EP ~ Eye.pair, data = fam_dat)
# Compute Shapiro-Wilk test of normality
shapiro_test(residuals(model))
#DOES NOT MEET NORMALITY ASSUMPTION

#Homogneity of variance
#The residuals versus fits plot can be used to check the homogeneity of variances.
plot(model, 1)

#compute
logEP <- as.numeric(fam_dat$log_EP)
eyepair <- factor(c(fam_dat$Eye.pair))
KWmodel <- kruskal.test(logEP~eyepair)
KWmodel
#significant, run Dunns
dunn_result <- dunnTest(logEP ~ eyepair,method="bonferroni")
dunn_result

##############################LYNIPHID##################################################
fam_dat = total_df %>% filter(Family == "Lyniphiadesidae")

#check for outliers
outliers <- fam_dat %>% group_by(Eye.pair) %>%
  identify_outliers(log_EP)
fam_dat <- fam_dat %>% anti_join(outliers, by = c("Eye.pair", "log_EP"))
fam_dat %>% group_by(Eye.pair) %>%
  identify_outliers(log_EP)

#check for normality
##Analyzing the ANOVA model residuals 
#Build the linear model
model  <- lm(log_EP ~ Eye.pair, data = fam_dat)
# Create a QQ plot of residuals
ggqqplot(residuals(model))
model <- aov(log_EP ~ Eye.pair, data = fam_dat)
# Compute Shapiro-Wilk test of normality
shapiro_test(residuals(model))
#In the QQ plot, as all the points fall approximately along the reference line, we can assume normality. This conclusion is supported by the Shapiro-Wilk test. The p-value is not significant (p = 0.128), so we can assume normality.

#Homogneity of variance
#The residuals versus fits plot can be used to check the homogeneity of variances.
plot(model, 1)
#In the plot above, there is no evident relationships between residuals and fitted values (the mean of each groups), which is good. So, we can assume the homogeneity of variances.

#compute
res.aov <- aov(log_EP ~ Eye.pair, data = fam_dat)
res.aov
report(res.aov)

tukey_result <- TukeyHSD(res.aov)
tukey_result

##########Thomisid ############
fam_dat = total_df %>% filter(Family == "Thomisidae")
#check for outliers
outliers <- fam_dat %>% group_by(Eye.pair) %>%identify_outliers(log_EP)
fam_dat <- fam_dat %>% anti_join(outliers, by = c("Eye.pair", "log_EP"))
fam_dat %>% group_by(Eye.pair) %>%
  identify_outliers(log_EP)

fam_dat <- as.data.frame(fam_dat)
##Analyzing the ANOVA model residuals 
#Build the linear model
model  <- lm(log_EP ~ Eye.pair, data = fam_dat)
# Create a QQ plot of residuals
ggqqplot(residuals(model))
model <- aov(log_EP ~ Eye.pair, data = fam_dat)
# Compute Shapiro-Wilk test of normality
shapiro_test(residuals(model))
#DOES NOT MEET NORMALACY TEST

#compute
logEP <- as.numeric(fam_dat$log_EP)
eyepair <- factor(c(fam_dat$Eye.pair))
KWmodel <- kruskal.test(logEP~eyepair)
KWmodel
#significant, run Dunns
#library(FSA)
dunn_result <- dunnTest(logEP ~ eyepair,method="bonferroni")
dunn_result

##########LYCOSIDAE############
fam_dat = total_df %>% filter(Family == "Lycosidae")
#check for outliers
outliers <- fam_dat %>% group_by(Eye.pair) %>%identify_outliers(log_EP)
fam_dat <- fam_dat %>% anti_join(outliers, by = c("Eye.pair", "log_EP"))
fam_dat %>% group_by(Eye.pair) %>%
  identify_outliers(log_EP)

fam_dat <- as.data.frame(fam_dat)
##Analyzing the ANOVA model residuals 
#Build the linear model
model  <- lm(log_EP ~ Eye.pair, data = fam_dat)
# Create a QQ plot of residuals
ggqqplot(residuals(model))
model <- aov(log_EP ~ Eye.pair, data = fam_dat)
# Compute Shapiro-Wilk test of normality
shapiro_test(residuals(model))

#Check normality for each group separately - was unconvinced by pholcid
fam_dat %>% group_by(Eye.pair) %>%
  shapiro_test(log_EP)
ggqqplot(fam_dat, "log_EP", facet.by = "Eye.pair")

#Homogneity of variance
#The residuals versus fits plot can be used to check the homogeneity of variances.
plot(model, 1)

#compute
res.aov <- aov(log_EP ~ Eye.pair, data = fam_dat)
res.aov
report(res.aov)

tukey_result <- TukeyHSD(res.aov)
tukey_result
```

# **Static allometry**

### Analysis (within species comparison)

##### Running OLS analysis on the Species data

```{r}
#read in the data and remove A. artema as there is too few data points
total <-read.csv('ln_total.csv')
total <- as_tibble(total)
total <- total %>%  filter(Species != "A. artema")

#create a list of species names and eye pair names
b <- total$Species[!duplicated(total$Species)]
c <- total$Eye.pair[!duplicated(total$Eye.pair)]
#create an empty data frame
ols_table <- data.frame()

for(i in b){ #for each species
  for(j in c){ #for each eye pair
#create a subset of the total data for that specific species + eye pair
test <- subset(total, Species == i & Eye.pair == j)
#run OLS
ols <- lm(formula = log_EP~log_CW, data = test)
#extract the confidence intervals
UpperCI <- ols$coefficients["log_CW"] + (1.96 * coef(summary(ols))["log_CW", "Std. Error"])
LowerCI <- ols$coefficients["log_CW"] - (1.96 * coef(summary(ols))["log_CW", "Std. Error"])
#add this to the empty dataframe
summaries <- summary(ols)
summaries <- summaries$coefficients
summaries <- as.data.frame(summaries)
ols_table <- rbind(ols_table, summaries)
  }
}
ols_table <- as.data.frame(ols_table)

#create a list of species to use to combine with the OLS csv in a datatable
species_lis <- list()
for(i in b){
  lis <- list(i)
  species_lis <- rbind(species_lis, i)
  species_lis <-rbind(species_lis, i)
  species_lis <-rbind(species_lis, i)
  species_lis <-rbind(species_lis, i)
  species_lis <- rbind(species_lis, i)
  species_lis <-rbind(species_lis, i)
  species_lis <-rbind(species_lis, i)
  species_lis <-rbind(species_lis, i)
}
species_lis <- as.data.frame(species_lis$V1)
write.csv(species_lis, "specieslis_ols.csv", row.names=FALSE)

write.csv(ols_table, "species_olstable.csv", row.names=FALSE)


#Extract R^2 values
r2_table <- data.frame()

for(i in b){ #for each species
  for(j in c){ #for each eye pair
#create a subset of the total data for that specific species + eye pair
test <- subset(total, Species == i & Eye.pair == j)
#run OLS
ols <- lm(formula = log_EP~log_CW, data = test)
#add this to the empty dataframe
summaries <- summary(ols)
summaries <- summaries$r.squared
to_add <- list(i, j, summaries)
r2_table <- rbind(r2_table,to_add)
  }
}
write.csv(r2_table,"ols_r2.csv", row.names=FALSE)
```

##### **Running linear mixed model for each species**

```{r}
total <-read.csv('ln_total.csv')
ontogenetic_df <- data.frame(matrix(ncol = 0, nrow = 8))
species_list <- c(total$Species)
#!duplicated(species_list)
species_list <-species_list[!duplicated(species_list)]
#A. artema only has one sample, so remove 
species_list <-species_list[species_list != "A. artema"]
species_list <- as.list(species_list)
#loop to run the lmer for each species
for (i in 1:39){
  filter = species_list[i]
  print(filter)
  filter_data = total %>% filter(Species == filter)
  x = lmer(log_EP ~ log_CW * Eye.pair + (1|Number),REML=F,
                           na.action = na.omit, data = filter_data)
  print(summary(x))
  sutemp<- as.data.frame(coef(summary(x)))
  ontogenetic_df <- rbind(ontogenetic_df, temp)
}

multiplied_names <- t(as.data.frame(c(rep(species_list, each = 8))))
write.csv(ontogenetic_df, file = "onto_table.csv")
write.csv(multiplied_names, file = "multiple_names.csv")
```

Extracting slopes using emmtrends:
```{r}
species_lmm_emtrends <- data.frame()
for (i in 1:39){
  filter = species_list[i]
  filter_data = total %>% filter(Species == filter)
  namess <- rep(filter, times = 4)
  x = lmer(log_EP ~ log_CW * Eye.pair + (1|Number),REML=F,
                           na.action = na.omit, data = filter_data)
  comps<-emtrends(x, 'Eye.pair', var = 'log_CW')
  temp_table <- as.data.frame.array(summary(comps))
  temp_table <-cbind(temp_table, namess[1])
  colnames(temp_table)[7] <- "species"
  species_lmm_emtrends <- rbind(species_lmm_emtrends, temp_table)
}
write.csv(species_lmm_emtrends,file='species_lmm_emtrends.csv')
```

Extracting the slope and residuals from the linear mixed models

```{r}
#remove A. artema as there is not enough samples per species
total <- total %>%  filter(Species != "A. artema")
#order the dataset by species
total <- total[order(total$Species),]
#make a list of species
species_list <- c(total$Species)
species_list <-species_list[!duplicated(species_list)]
#Remove A. artema
species_list <-species_list[species_list != "A. artema"]
species_list <- as.list(species_list)
total_df <- data.frame(total)

#create a dataframe for the slopes
slopes_df <- data.frame(matrix(ncol = 0, nrow = 8))
#create an empty list for residuals
residuals <- list()
#rsquared <- list()
#loop to run the lmer for each species
for (i in 1:39){
  filter = species_list[i]
  print(filter)
  filter_data = total_df %>% filter(Species == filter)
  x = lmer(log_EP ~ log_CW * Eye.pair + (1|Number),REML=F,
                           na.action = na.omit, data = filter_data)
  comps<-emtrends(x,'Eye.pair',var = 'log_CW')
  print(summary(x))
  temp<- coef(summary(x))
  slopes_df <- cbind(slopes_df, temp[,1])
  colnames(slopes_df)[i] = filter
  print(pairs(comps))
  residuals <- list.append(residuals,x@sigma)
}

#create a dataframe for the slopes 
slopes_df <- t(slopes_df)
species_vector <- unlist(species_list)
#create a dataframe for the residuals
residuals_vector <- unlist(residuals)
residual_df <-data.frame(species_vector, residuals_vector)

write.table(slopes_df,file ="slope_df.csv")
write.table(residual_df, file = "residual_df.csv")
```

Creating a boxplot of the residuals

```{r}
residuals <- read.csv('residuals.csv')

residual_plot <- ggplot(residuals, aes(x=Family, y=R.2, fill = Eye.pair)) + geom_boxplot() + scale_fill_manual(values =c('#eded15','#c360dc', '#e20505', '#09ef09')) + labs(x = "Family", y = 'r^2') + 
  guides(fill=guide_legend(title="Eye pair")) + theme_light() + scale_y_continuous(limits = c(-0, 0.053))
residual_plot  

ggsave(residual_plot, file=paste0("residual.pdf"), width = 44.45, 
         height = 27.78, units = "cm", dpi=300)
```

### Figures

**Graphs for each eye pair per family with species highlighted**

```{r}
#create a list of all the spider families
family_list <- c(total$Family)
!duplicated(family_list)
family_list <- family_list [!duplicated(family_list)]
family_list <- as.list(family_list)

#run a loop for each eye pair per family
for(j in family_list){
  fam_dat = total_df %>% filter(Family == j)
  species_list <- c(fam_dat$Species)
 # !duplicated(species_list)
  species_list <-species_list[!duplicated(species_list)]
  species_list <- as.list(species_list)
  fam_df <- data.frame(matrix(ncol = 7, nrow = 0))
  for (i in species_list){
    name <- rep(i, times = 20)
    name <- unlist(name)
    dat = fam_dat %>% filter(Species == i)
    x = lmer(log_EP ~ log_CW * Eye.pair + (1|Number),REML=F,
                           na.action = na.omit, data = dat)
  
    effects_allom <- effects::effect(term = "log_CW * Eye.pair", mod = x)
    x_allom <- data.frame(effects_allom)
    temp <- cbind(x_allom, name)
    fam_df <-rbind(fam_df, temp)
    temp<- data.frame(matrix(ncol = 7, nrow = 0))
    }
  colnames(fam_df)[7] ="Spec"
  fam_plot <- ggplot(data = fam_dat, aes(x = log_CW, y = log_EP,lty = Species, col = Eye.pair)) +   
  geom_point(size = 2,alpha = 0.6, aes(shape = Species)) +
  facet_wrap(~Eye.pair,scales = "free") +
  geom_line(data=fam_df, aes(x = log_CW,y=fit, lty = Spec, col = Eye.pair),lwd=1.5)+
  ylab("Log Eye Pair") + xlab("Log Carapace Width") + scale_linetype(name = "Eye Pair") +
  scale_color_manual(values=c('#09ef09','#c360dc', '#e20505', '#eded15'))  + theme_light() + xlim(5, 8.50)+ylim(3.20, 6.6)
  ggsave(fam_plot, file=paste0("plot_", j,".pdf"), width = 44.45, 
         height = 27.78, units = "cm", dpi=300)
  }
  
```

### Analysis (family level)

Set up, create a family dataset that contains the regression line information + extract slope + intercept

```{r}
total <-read.csv('total.csv')
total <- as_tibble(total)
total <- total %>%  filter(Species != "A. artema")
total_df <- data.frame(total)
#A. artema only has one sample, so remove 
family_list <- c(total$Family)
family_list <- family_list [!duplicated(family_list)]
family_list <- as.list(family_list)

#extract model information 
fam_df <- data.frame(matrix(ncol = 7, nrow = 0))
for(j in family_list){
  fam_dat = total_df %>% filter(Family == j)
  species_list <- c(fam_dat$Species)
  species_list <-species_list[!duplicated(species_list)]
  species_list <- as.list(species_list)
  for (i in species_list){
    name <- rep(i, times = 20)
    name <- unlist(name)
    dat = fam_dat %>% filter(Species == i)
    x = lmer(log_EP ~ log_CW * Eye.pair + (1|Number),REML=F,
             na.action = na.omit, data = dat)
    
    effects_allom <- effects::effect(term = "log_CW * Eye.pair", mod = x)
    x_allom <- data.frame(effects_allom)
    temp <- cbind(x_allom, name)
    fam_df <-rbind(fam_df, temp)
    temp<- data.frame(matrix(ncol = 7, nrow = 0))
  }
}
write.csv(fam_df,file='fam_df.csv')

#for each species, for each eye pair, create an equation, run this for each species to find residuals
test_df <-read.csv("testdf.csv")
model = lm(test_df$fit ~ test_df$log_CW)

summary_df <- data.frame(species = character(), eye_pair = character(),
                         intercept = numeric(), slope = numeric(), stringsAsFactors = FALSE)

specieslist <- c(fam_df$name)
specieslist <- specieslist[!duplicated(specieslist)]
specieslist <- as.list(specieslist)
ep_list <- list("ALE", "AME", "PLE", "PME")
for(a in specieslist){
  for(b in ep_list){
    temp_df <- fam_df %>% filter(name == a)
    temp_df <- temp_df %>% filter(Eye.pair == b)
    model = lm(temp_df$fit ~ temp_df$log_CW)
    new_row <- data.frame(species = a , eye_pair = b,
                          intercept = model$coefficients[1], slope = model$coefficients[2], stringsAsFactors = FALSE)
    summary_df <- rbind(summary_df, new_row)
  }
}

write.csv(summary_df,file='summary_df.csv')
```

**Running mixed model**

```{r}
##set up datasets for the families
lycosid <- total %>% filter(Family == 'Lycosidae')
therid <- total %>% filter(Family == 'Theridiidae')
thomisid <- total %>% filter(Family == 'Thomisidae')
lyniphid<- total %>% filter(Family == 'Lyniphiadesidae')
araneid<- total %>% filter(Family == 'Araneidae')
gnaphosid<- total %>% filter(Family == 'Gnaphosidae')
salticid<- total %>% filter(Family == "Salticidae")

#LYCOSIDS
#Full, maximum likelihood
lycosid_ML_full<-lmer(log_EP ~ log_CW * Eye.pair * Species+ (1|Number),REML=F,na.action = na.omit, data = lycosid)
#Reduced, maximum likelihood
lycosid_ML_red<-lmer(log_EP ~ (log_CW + Eye.pair + Species )^2+(1|Number),REML=F,na.action = na.omit,data = lycosid)
#Testing the significance of fixed effects
##Satterthwaite’s method for estimating fixed effects finds a significant three-way interaction.
anova(lycosid_ML_full, lycosid_ML_red)
#Full has a smaller AIC
anova(lycosid_ML_full)
report(anova(lycosid_ML_full))

summary(emtrends(lycosid_ML_full, 'Eye.pair', var = 'log_CW'))

#THERID
dat = therid
#Full, maximum likelihood
spider_ML_full<-lmer(log_EP ~ log_CW * Eye.pair * Species+ (1|Number),REML=F,na.action = na.omit, data = dat)
#Reduced, maximum likelihood
spider_ML_red<-lmer(log_EP ~ (log_CW + Eye.pair + Species )^2+(1|Number),REML=F,na.action = na.omit,data = dat)
#Testing the significance of fixed effects
##Satterthwaite’s method for estimating fixed effects finds a significant three-way interaction.
anova(spider_ML_full, spider_ML_red)
#Full has a smaller AIC
anova(spider_ML_full)
report(anova(spider_ML_full))
summary(emtrends(spider_ML_full, 'Eye.pair', var = 'log_CW'))
#Thomisid
dat = thomisid
#Full, maximum likelihood
spider_ML_full<-lmer(log_EP ~ log_CW * Eye.pair * Species+ (1|Number),REML=F,na.action = na.omit, data = dat)
#Reduced, maximum likelihood
spider_ML_red<-lmer(log_EP ~ (log_CW + Eye.pair + Species )^2+(1|Number),REML=F,na.action = na.omit,data = dat)
#Testing the significance of fixed effects
##Satterthwaite’s method for estimating fixed effects finds a significant three-way interaction.
anova(spider_ML_full, spider_ML_red)
#Full has a smaller AIC
anova(spider_ML_full)

report(anova(spider_ML_full))
summary(emtrends(spider_ML_full, 'Eye.pair', var = 'log_CW'))
#lyniphid
dat = lyniphid
#Full, maximum likelihood
spider_ML_full<-lmer(log_EP ~ log_CW * Eye.pair * Species+ (1|Number),REML=F,na.action = na.omit, data = dat)
#Reduced, maximum likelihood
spider_ML_red<-lmer(log_EP ~ (log_CW + Eye.pair + Species )^2+(1|Number),REML=F,na.action = na.omit,data = dat)
#Testing the significance of fixed effects
##Satterthwaite’s method for estimating fixed effects finds a significant three-way interaction.
anova(spider_ML_full, spider_ML_red)
#Full has a smaller AIC
anova(spider_ML_full)
report(anova(spider_ML_full))
summary(emtrends(spider_ML_full, 'Eye.pair', var = 'log_CW'))
#araneid
dat = araneid
#Full, maximum likelihood
spider_ML_full<-lmer(log_EP ~ log_CW * Eye.pair * Species+ (1|Number),REML=F,na.action = na.omit, data = dat)
#Reduced, maximum likelihood
spider_ML_red<-lmer(log_EP ~ (log_CW + Eye.pair + Species )^2+(1|Number),REML=F,na.action = na.omit,data = dat)
#Testing the significance of fixed effects
##Satterthwaite’s method for estimating fixed effects finds a significant three-way interaction.
anova(spider_ML_full, spider_ML_red)
#Full has a smaller AIC
anova(spider_ML_full)

report(anova(spider_ML_full))
summary(emtrends(spider_ML_full, 'Eye.pair', var = 'log_CW'))
#gnaphosid
dat = gnaphosid
#Full, maximum likelihood
spider_ML_full<-lmer(log_EP ~ log_CW * Eye.pair * Species+ (1|Number),REML=F,na.action = na.omit, data = dat)
#Reduced, maximum likelihood
spider_ML_red<-lmer(log_EP ~ (log_CW + Eye.pair + Species )^2+(1|Number),REML=F,na.action = na.omit,data = dat)
#Testing the significance of fixed effects
##Satterthwaite’s method for estimating fixed effects finds a significant three-way interaction.
anova(spider_ML_full, spider_ML_red)
#Full has a smaller AIC
anova(spider_ML_red)
report(anova(spider_ML_red))
summary(emtrends(spider_ML_full, 'Eye.pair', var = 'log_CW'))
#salticid
dat = salticid

spider_ML_full<-lmer(log_EP ~ log_CW * Eye.pair * Species+ (1|Number),REML=F,na.action = na.omit, data = dat)
#Reduced, maximum likelihood
spider_ML_red<-lmer(log_EP ~ (log_CW + Eye.pair + Species )^2+(1|Number),REML=F,na.action = na.omit,data = dat)
#Testing the significance of fixed effects
##Satterthwaite’s method for estimating fixed effects finds a significant three-way interaction.
anova(spider_ML_full, spider_ML_red)
#Full has a smaller AIC
anova(spider_ML_red)

report(anova(spider_ML_red))
summary(emtrends(spider_ML_red, 'Eye.pair', var = 'log_CW'))
```

## Getting the comparisons:

Between eye pairs within a given species:

```{r}
total <-read.csv('ln_total.csv')
total <- as_tibble(total)
total <- total %>%  filter(Species != "A. artema")
total <- total[order(total$Species),]
species_list <- c(total$Species)
#!duplicated(species_list)
species_list <-species_list[!duplicated(species_list)]
#A. artema only has one sample, so remove 
species_list <-species_list[species_list != "A. artema"]
species_list <- as.list(species_list)
total_df <- data.frame(total)

for (i in 1:39){
  filter = species_list[i]
  filter_data = total_df %>% filter(Species == filter)
  x = lmer(log_EP ~ log_CW * Eye.pair + (1|Number),REML=F,
                           na.action = na.omit, data = filter_data)
  comps<-contrast(emtrends(x, 'Eye.pair', var = 'log_CW'), method = "pairwise")
  file <- paste("within_species_comparison/",filter,".csv")
  write.csv(comps, file, row.names = FALSE)
}
```

Do we see the same trends at the family level? Aka, are species within a family showing the same dynamics?

```{r}
family_list <- c(total$Family)
family_list <- family_list [!duplicated(family_list)]
family_list <- as.list(family_list)

for (j in 1:8){
  filt = family_list[j]
  fam_dat = total_df %>% filter(Family == filt)
  x = lmer(log_EP ~ log_CW * Eye.pair + (1|Species/Number),REML=F,
                           na.action = na.omit, data = fam_dat)
  comps<-contrast(emtrends(x, 'Eye.pair', var = 'log_CW'), method = "pairwise")
  file <- paste("within_family_comparison/",filt,".csv")
  write.csv(comps, file, row.names = FALSE)
}
```
````
